# Structural basis for multiple binding modes of antimalarial ligands targeting *Plasmodium falciparum* NCR1

**DOI:** 10.1101/2025.11.11.687791

**Authors:** Aaron Kohling, Coralie Boulet, Mounaf Al Makhlouf, Basavraj Khanppnavar, Ya Chen, Emma Ganga, Johannes Kirchmair, Thomas Stockner, Mathieu Brochet, Volodymyr M. Korkhov

## Abstract

PfNCR1 is a *Plasmodium falciparum* cholesterol transporter at the plasma membrane–parasitophorous vacuole interface, which has recently emerged as a promising antimalarial target. Despite an immense interest in development of novel antimalarial compounds targeting PfNCR1, the molecular mechanism of PfNCR1 inhibition remains elusive. Here, we report cryo-EM structures of PfNCR1 in its apo state and bound to three inhibitors: MMV009108, MMV019662 and MMV028038. MMV009108 binds to the “neck” site at the ectodomain-membrane domain interface. MMV028038 displaces the sterol at the ectodomain “ecto” site. Remarkably, MMV019662 binds both sites: it associates near the bound sterol molecule at the ecto site and targets the neck site, thereby altering the sterol-sensing domain conformation. Importantly, we identify a novel antimalarial compound, G856-4236, which targets the ecto site exclusively. These four distinct modes of PfNCR1 inhibition advance our understanding of its conformational plasticity and establish a framework for rational drug discovery targeting PfNCR1 and related transporters.

## Introduction

Malaria is a mosquito-borne disease that affects a staggering number of people worldwide, with 280 million cases and 610 000 deaths globally in 2024 ^1^. The causative agent of malaria is a unicellular eukaryotic parasite of the *Plasmodium* genus, which belongs to the phylum of Apicomplexa. *P. falciparum* is the most virulent and accounts for the majority of malaria-associated deaths ^1^. The parasite undergoes a complex life cycle involving multiple developmental stages ^2^. Following transmission through the bite of an infected *Anopheles* mosquito, *P. falciparum* sporozoites enter the human blood-stream and migrate to the liver. There, they invade hepatocytes, replicate extensively, and release thousands of merozoites into the bloodstream. Once in circulation, merozoites invade red blood cells (RBCs), initiating the asexual blood stages (ABS), which are responsible for the clinical manifestations of malaria including fever, anemia, organ dysfunction, and potentially death. During ABS replication, the parasite progresses through distinct intraerythrocytic stages: ring, trophozoite, and schizont. Mature schizonts rupture to release up to 32 new merozoites, perpetuating the cycle. In parallel, a subset of parasites commits to sexual differentiation, forming male and female gametocytes. These gametocytes are taken up by mosquitoes during a blood meal, where they mature into gametes and undergo fertilization. The resulting zygote develops into a motile ookinete, which traverses the midgut epithelium to form an oocyst on the outer gut wall, where it undergoes sporogony to produce thousands of sporozoites that eventually migrate to the mosquito’s salivary glands completing the cycle and enabling transmission to a new human host ^2^.

Currently available vaccines targeting *P. falciparum* have limited efficacy and do not yet offer sufficient protection against malaria ^3^, which is why we still need to heavily rely on antimalarial drugs. Despite the availability of drugs for both prophylaxis and treatment, malaria remains a major global health challenge owing to drug resistance, limited healthcare access, and the differential susceptibility of the parasite’s life-cycle stages to antimalarial therapies. Several of the currently available antimalarial drugs are used either individually or in combination. These include: mefloquine, artemisinin (and its derivatives), atovaquone, pyronaridine, proguanil and several others ^4^. All of these drugs target the symptomatic ABS of *P. falciparum* within the human host. A subset also targets liver stages, and only a minority are effective against gametocytes ^5^. This reduced susceptibility of gametocytes may stem from their distinct biology, including lower metabolic activity and altered drug permeability ^6^. This represents a major limitation of current control strategies, as patients undergoing treatment may remain infectious for a period of time. To effectively interrupt malaria transmission, there is a pressing need for new drugs that target gametocytes.

Primary mechanisms of action have been proposed for many of available drugs. For example, mefloquine has been shown to act on the *P. falciparum* 80S ribosome, specifically inhibiting protein synthesis in the parasite ^7^. Atovaquone inhibits mitochondrial electron transport by targeting the *P. falciparum* cytochrome bc1 ^8^. Moreover, some of the drugs may utilize multiple modes of action. For example, it has been suggested that artemisinin and its derivatives could act via multiple potentially connected molecular pathways, from heme detoxification ^9^, to covalent protein modification ^10^, proteasome inhibition ^11^, inhibition of phosphatidyninositol-3-kinase (PfPI3K) ^12^. Similarly, pyronaridine has been found to interfere with b-hematin formation, in addition to intercalating into DNA and inhibiting DNA topoisomerase 2 ^13^. The antimalarial drug proguanil is a precursor for its presumed active metabolite cycloguanil, which is a dihydrofolate reductase (DHFR) inhibitor ^14^. Remarkably, proguanil has been suggested to synergize with atovaquone in the absence of cycloguanil ^15^, and was found to potentiate atovaquone’s ability to collapse the mitochondrial membrane potential ^16^, indicating that the mechanism of proguanil action is complex. While several drugs show efficacy in preventing and treating malaria, the emergence of *P. falciparum* strains resistant to antimalarial drug treatment ^17^ highlights the necessity of further development of therapeutic approaches with new drug targets and modes of action to retain efficacy across a broad range of existing and emerging parasite strains.

A recent study identified the *P. falciparum* Niemann-Pick type C1 (NPC1)-related protein (PfNCR1, PF3D7_0107500) as a promising new antimalarial drug target (**Figure 1A**) ^18^. Mutations of PfNCR1 identified by *in vitro* evolution and whole genome analysis were found to alter susceptibility of *P. falciparum* to three antimalarial compounds in the Medicines for Malaria Venture (MMV) Malaria Box library, MMV009108 (hereafter referred to as Compound 1), MMV028038 (Compound 2) and MMV019662 (Compound 3) (**Figure 1B**). PfNCR1 localizes to the plasma membrane of *P. falciparum* at sites of contact with the parasitophorous vacuole (PV), a membrane compartment formed upon invasion that surrounds the parasite and provides a specialized interface with the host cell. At these contact sites, PfNCR1 is thought to regulate cholesterol exchange between membranes ^19^. Recent work has shown that PfNCR1 acts as a cholesterol exporter, removing cholesterol from the parasite plasma membrane ^20^. Inhibition of this function by MMV compounds targeting PfNCR1 likely explains their antimalarial activity ^18^.

**Figure 1.**
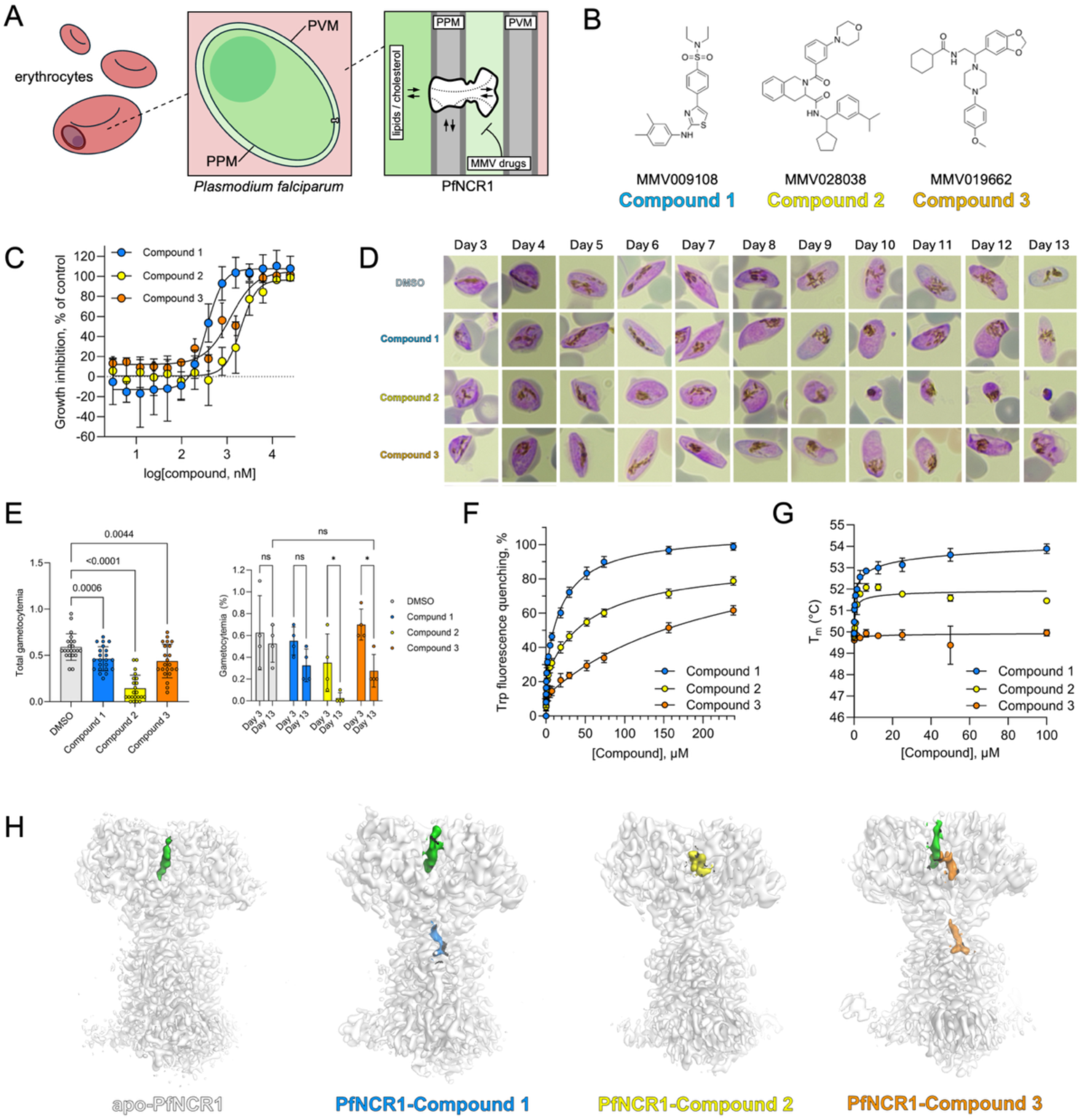
Cryo-EM reconstructions of PfNCR1 in apo-and MMV compound-bound states. **A.** A sketch illustrating the cholesterol export pathway involving PfNCR1 at the interface between the *Plasmodium* plasma membrane (PPM) and the parasitophorous vacuole membrane (PVM). **B.** Three MMV compounds used in this study: MMV009108 (Compound 1), MMV028038 (Compound 2), MMV019662 (Compound 3). **C.** *P. falciparum* asexual growth inhibition assay over 72 h: EC_50_ for Compounds 1, 2 and 3 = 0.404, 2.37 and 1.18 µM respectively. **D-E**. Gametocytogenesis was induced, and parasites exposed to 3 x EC_90_ of Compound 1, 2 or 3 (*i.e.* 3 µM, 15 µM and 15µM) or the DMSO vehicle control the following day. Gametocytemia was counted every day and parasites imaged on a thin blood smear. **F-G.** PfNCR1 Trp fluorescence quenching-based ligand binding assay (F) and thermostability assays (G) in the presence of the three MMV compounds. **H.** Cryo-EM maps of apo-PfNCR1 and the three ligand-bound states, Compound 1-(blue), Compound 2-(yellow), and Compound 3 (orange)-bound states. A bound sterol (CHS) is green.

PfNCR1 belongs to the Resistance-Nodulation-Cell Division (RND) family of transporters. Members of this family act in prokaryotic organisms as multidrug resistance transporters (AcrB ^21^) or transporters for sterol-like molecules, hopanoids (HpnN ^22^). In eukaryotes, including humans, some of the RND family members are involved in membrane homeostasis-and signaling-related tasks. For example, the namesake of PfNCR1, NPC1, is a lysosomal cholesterol transporter that is associated with the Niemann Pick disease, type C1, which is a lysosomal cholesterol storage disease ^23,24^. The NPC1-like protein, NPC1L1, is an NPC1 homologue that is important for intestinal cholesterol absorption ^25^. Other eukaryotic RND proteins, such as DISP1 (Dispatched in *Drosophila melanogaster*) and PTCH1 (Patched in *Drosophila melanogaster*), are key players in the hedgehog (Hh) signaling pathway ^26^. DISP1 is necessary for the release of the Hh morphogen from the Hh-producing cells, whereas PTCH1 acts as one of the key Hh receptors. The ligand of these proteins, Hh, is a small protein modified by a palmitoyl and a cholesteryl moiety ^26^. Both PTCH1 ^27^ and DISP1 ^28^ interact with the cholesteryl group of the Hh ligand, and PTCH1 has been suggested to act as a cholesterol transporter moving the sterol from the inner leaflet of the lipid bilayer and thus preventing Hh pathway activation ^26^. Thus, RND transporters play important roles in cell survival, lipid homeostasis and signaling.

Structurally, the RND protein family shares a polytopic membrane domain of 12-13 transmembrane (TM) helices and two extracellular ectodomains (ECD). All eukaryotic RND transporters contain a Sterol-Sensing Domain (SSD) ^29^. The SSD was originally proposed to be formed by the TM2-6 helical bundles of the SSD-containing proteins. However, in light of recent structural insights, the SSD was suggested to include two additional conserved structural motifs: the N-terminal transverse helix (TH), which is an amphipathic helix at the cytosol-membrane interface, and the TM1 that immediately follows the TH ^29^. The SSD regions in the RND transporters are similar to the SSDs originally identified in the membrane proteins involved in cholesterol metabolism control, SCAP ^30^ and HMG CoA reductase ^29,31^. While the SSD regions in SCAP and HMG CoA reductase appear to have a function in sensing the abundance of cholesterol in the lipid bilayer, these domains in the RND transporters are presumed to play an important role in cholesterol translocation. A widely accepted hypothesis is that the SSD provides an optimal surface for protein-sterol interactions and for guiding the sterols towards the sterol translocation pathway within the ECD region. This notion is supported by structural evidence indicating that the SSD can serve as a site of sterol binding and/or enrichment in NPC1L1 ^32^, Dispatched ^33^ or PTCH1 ^27^. However, how exactly SSDs facilitate the movement of sterols remains to be demonstrated.

Motivated by the need to characterize PfNCR1 both as a representative RND transporter and a putative drug target, we set out to determine its structure and obtain insights into the modes of action of the three MMV compounds known to act on PfNCR1. The results of our structural and biophysical investigations of PfNCR1-small molecule interactions, detailed below, open new opportunities for rational discovery of antimalarial drugs targeting PfNCR1.

## Results

### Biological activity of the MMV compounds

Three MMV compounds have previously been described to exert activity against *P. falciparum* ABS by targeting PfNCR1 (**Figure 1A-B**) ^18^. We first confirmed these compounds inhibited parasite asexual growth with EC50 values in the micromolar range: 0.4, 2.4 and 1.2 µM, respectively (**Figure 1C**). We next tested the activity of these compounds against gametocytes, which has not been assessed previously. Transcriptomic and proteomic data indicate that PfNCR1 is expressed in gametocytes, albeit at lower levels than in ABS ^34,35^, raising the possibility that PfNCR1 inhibition may also affect gametocytes and thereby contribute to blocking parasite transmission to mosquitoes. We therefore sought to assess the transmission-blocking potential of MMV009108 (Compound 1), MMV028038 (Compound 2) and MMV019662 (Compound 3). To do so, we first assessed the expression of PfNCR1 in gametocytes. Using *P. falciparum* expressing GFP-tagged PfNCR1 ^18^, we demonstrated both by flow cytometry and western blot analyses that PfNCR1-GFP is indeed expressed in gametocytes, although at lower levels than in ABS (Figure S1-2). To examine the effect of compounds 1-3 on gametocytogenesis, gametocytes were exposed throughout their 13-day developmental period to each compound at three times the EC_90_ concentration determined for ABS, *i.e.* 3 µM, 15 µM and 15 µM respectively for compound 1, 2 and 3 (**Figure 1D, E**). All compounds altered gametocytes morphology from day 6 onwards, with compound 2 displaying the most pronounced impacts (**Figure 1D**). The typical elongation of stage IV gametocytes was lost upon treatment with compound 2 and further development was abolished. With compounds 1 and 3, stage V gametocytes were rounder than the DMSO-treated parasites and appeared abnormal by day 13. Importantly, all three compounds significantly reduced overall gametocytemia (**Figure 1E**), with compound 2 exhibiting again the strongest effect. Overall, compounds 1-3 target *P. falciparum* ABS as well as gametocyte formation, which may be beneficial for developing new drugs that prevent transmission to mosquitoes.

### *In vitro* analysis of PfNCR1-ligand interactions

To characterize protein-ligand interactions *in vitro*, we expressed PfNCR1 in Freestyle 293 cells, purified the protein using affinity and size exclusion chromatography in detergent (glyco-diosgenin, GDN) (Figure S3) and performed tryptophan fluorescence quenching and thermostability assays. The tryptophan quenching-based binding assays revealed the apparent K_d_ values in the micromolar range for the three compounds (**Figure 1F**). Interestingly, while Compound 1 and Compound 2 stabilized the protein by ∼4°C and ∼2°C, respectively, Compound 3 showed no thermostabilization of the purified PfNCR1 (**Figure 1G**).

Nevertheless, we proceeded to analyze the three purified PfNCR1-ligand complexes, along with the apo-PfNCR1 in the absence of added compounds, using cryo-EM and single particle analysis. We prepared cryo-EM grids with the purified PfNCR1 in the presence of Compound 1, 2 or 3, at a final drug concentration of 1 mM. Single particle cryo-EM analysis resulted in four 3D reconstructions: apo-PfNCR1 and Compound 1-, Compound 2-and Compound 3-bound states, at 3.04 Å, 3.89 Å, 3.69 Å and 3.73 Å resolution, respectively (**Figure 1H**, Figure S4-12).

### Cryo-EM structure of apo-PfNCR1

The structure of the apo-state of the protein was consistent with the prediction by AlphaFold2 ^36^, as well as with the structure of PfNCR1 published recently (Figure S13) ^20^. Several loops of the protein were not well resolved, but the 12-TM bundle and the two ECD lobes were clearly defined (**Figure 2A**). Interestingly, the ECD features a density consistent with a bound sterol molecule, which we interpreted as cholesterol hemisuccinate (CHS), a cholesterol analogue that was added during sample preparation to stabilize the protein for structural studies (detailed in “Materials and Methods”). Unambiguous placement of the CHS molecule was challenging at the resolution of our 3D reconstruction. We therefore validated the orientation of the best four CHS orientations using 5 independent repeats of 0.5 μs long molecular dynamics (MD) simulations for each of the four orientations (**Figure 2B**, Figure S14). Of these, the two orientations of CHS with the polar headgroup (linked to the hemisuccinate moiety) oriented towards the core of the ECD showed higher stability and consistency with the cryo-EM density in explored conformational space and deviating less from the starting cryo-EM structure (RMSD of pose 1: 4.42±0.90 Å; pose 3: 4.14±0.85 Å), indicating that pose 3 leads to the most stable trajectories. In both orientation, cholesterol rotated around its long axis, but remained within the experimentally derived density. In contrast, the CHS with the inverted orientation deviated from the cryo-EM density, especially towards the conduit (RMSD of pose 2: 5.05±1.63 Å; pose 4: 5.11±1.43 Å), where the cryo-EM density and the simulation derived density did not overlap. The 2D RMSD plots (Figure S14) show a much higher consistency across the 5 repeats. These 10 consistent simulations also showed convergence towards a common orientation, supporting the ligand placement. We find that the cholesterol moiety remains dynamic, but confined, which is consistent with the limited local resolution in the cryo-EM density for CHS. This observed flexibility may reflect a functional requirement for a relatively loose binding of cholesterol to its binding site to permit cholesterol translocation by PfNCR1. The originally modelled sterol conformation with the lowest root-mean-square deviation of CHS was thus selected. This conformation is similar to the conformation of the cholesterol molecule in the structure of PfNCR1 purified in GDN without added CHS (with endogenous recombinant host cell cholesterol co-isolated with the purified protein sample) ^20^, with only a slight deviation in the sterol orientation (Figure S13).

**Figure 2.**
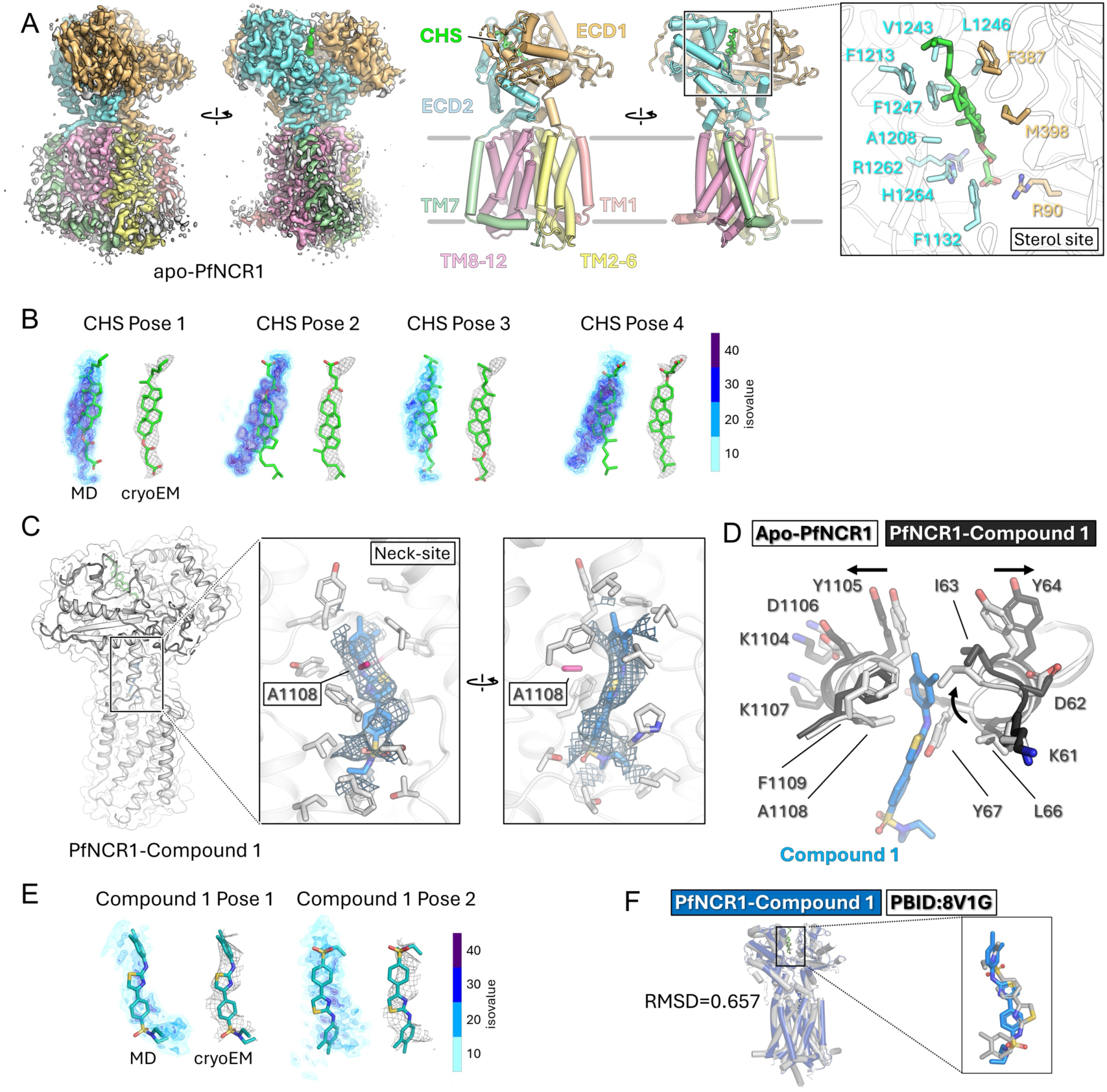
Cryo-EM structures of the apo-PfNCR1 and the PfNCR1-MMV009108 (Compound 1) complex. **A.** Cryo-EM map (left) and model (right) of apo-PfNCR1. Distinct domains of the protein are labelled and colour-coded. Sterol (CHS) orientation in the ECD sterol site. Side chains of the residues in ECD1 (light orange) and ECD2 (cyan) of PfNCR1 within 4 Å of the CHS molecule are shown as sticks labeled. **B.** Comparison between the cryo-EM densities and the MD-based densities of CHS, in gray mesh representation of the cryo-EM density, together with the suggested pose of the ligand in the stick representation, while the MD-based ligand densities are shown as a blue volume representation. The colour scale reflects the percent occupancy of the respective position in space. **C.** Details of the neck site with Compound 1 bound, including the key residues Compound 1 bound to the neck site. **D.** A comparison of the apoand Compound 1-bound conformations of PfNCR1 shows a slight shift of the residues in the neck-forming helices (K61-Y67 and K1104-F1109), indicated by arrows. The carbon atoms in apo-PfNCR1 are light-grey, in Compound 1-bound PfNCR1 – dark grey. **E.** Comparison between the cryo-EM densities and the MD densities of Compound 1, in gray mesh representation of the cryo-EM density together the suggested pose of the ligand in the stick representation, while the MD-based densities are shown as a blue volume representation reflecting percent occupancy. **F.** A comparison of the Compound 1-bound conformation of PfNCR1 in blue and the published 8V1G in gray shows overall very low RMSD, although Compound 1 in 8V1G was placed upside down compared to our MD simulation-backed model.

### Structure of the Compound 1-bound PfNCR1

Cryo-EM analysis of the Compound 1-bound PfNCR1 revealed a binding site in the “neck” region and the mutation A1108T previously shown to disrupt the Compound 1 binding site in PfNCR1 ^18^ maps to the neck site (**Figure 2C**). Binding of Compound 1 does not lead to substantial conformational changes, with only small adjustments of the residues Y1105-F1109 and I63-Y64 in the short α-helical stretches that form the “neck” site accommodating the ligand (**Figure 2D**). The resolution of the cryo-EM density map was insufficient to unambiguously model the bound Compound 1. The MD simulations carried out for two possible orientations of Compound 1 found that pose 1 was more stable (RMSD of pose 1: 2.66±0.41 Å; pose 2; 3.67±0.47 Å) and consistent with the cryo-EM density (**Figure 2E)** and 2D RMSD plots (Figure S15). Interestingly, the recently determined structure of the PfNCR1 bound to Compound 1 ^20^ featured the compound bound to this site in the orientation that appears less favourable based on our MD simulations, highlighting the importance of independently evaluating ambiguous ligand density assignment of cryo-EM data (**Figure 2F**).

### Structure of the Compound 2-bound PfNCR1

In stark contrast to Compound 1, the reconstruction of Compound 2 showed that this ligand completely displaces the CHS molecule in the ECD sterol site, while occupying a partially overlapping but distinct binding pocket, the “ecto site” (**Figure 3A**). The conformation of the neck region was not affected by Compound 2 binding to the ecto site, and the overall conformation was similar to apo-PfNCR1. Like Compound 1, the mutations conferring a change in sensitivity of *P. falciparum* to Compound 2, M398I and A1208E ^18^, map to the ecto site in proximity to the bound Compound 2 (**Figure 3B**). The cavity between the two lobes of the ECD is large and could, in principle, accommodate a wide range of ligands, including Compound 2, which is quite bulky. Modeling of Compound 2 in the ecto site was guided by MD simulations, revealing that only one of two potentially fitting ligand poses was stable (RMSD pose 1: 4.23±1.43 Å; pose 2: 4.19±0.66 Å) (**Figure 3C)** and 2D RMSD plots (Figure S15).

**Figure 3.**
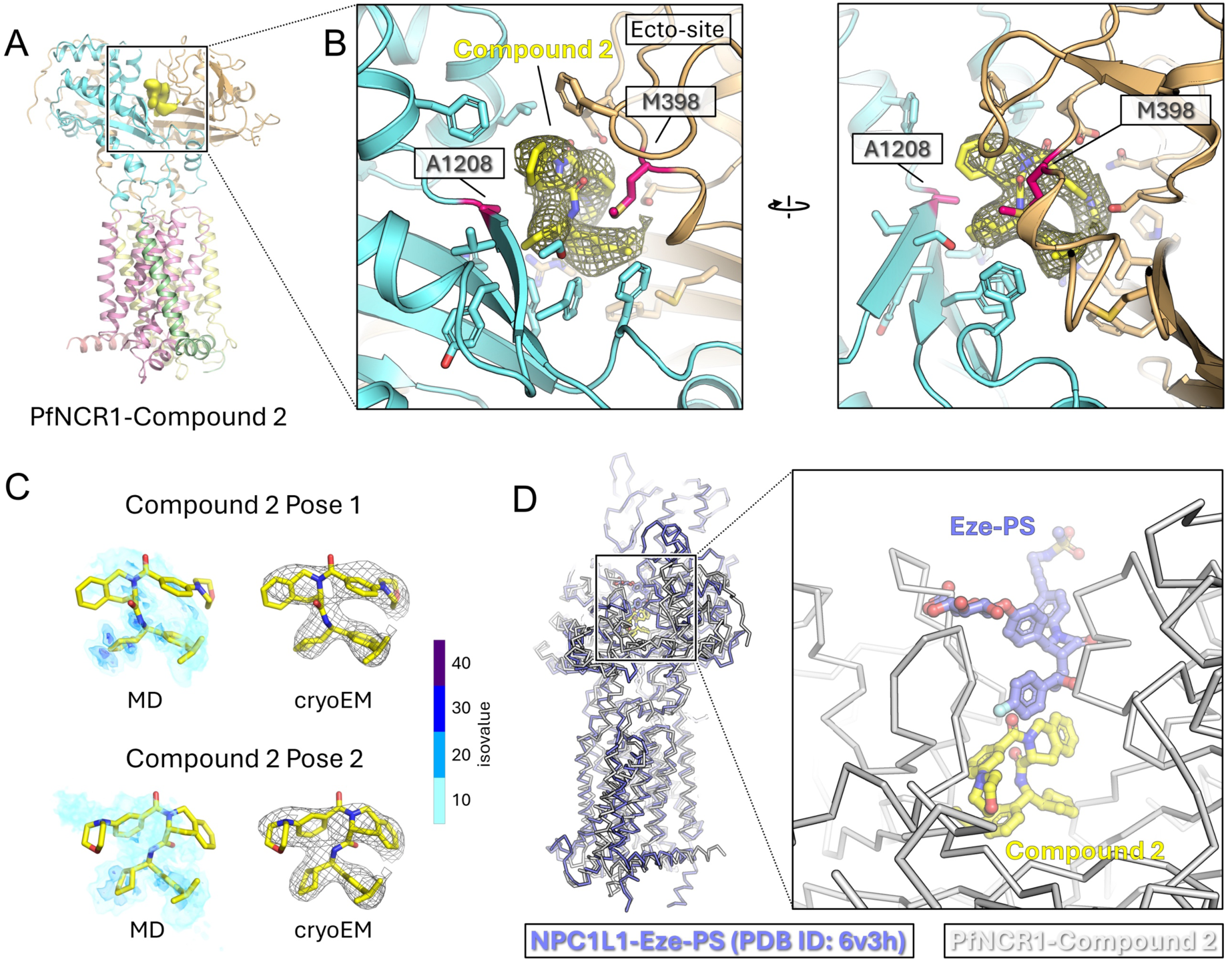
Cryo-EM structure of the PfNCR1-MMV028038 (Compound 2) complex. **A.** Model of the Compound 2-PfNCR1 complex and the map corresponding to the bound Compound 2. The ectodomain-bound sterol is competitively displaced by Compound 2. **B.** Details of the binding site; the residues corresponding to the inactivating mutations M398I and A1208E are coloured pink. **C.** Comparison between the cryo-EM densities and the MD-based densities of Compound 2, in gray mesh representation of cryo-EM density showing the suggested pose of the ligand in the stick representation, while the MD-based ligand densities shown as a blue volume representation reflecting percent occupancy. **D.** Comparison with the previously determined structure of an RND transporter with a drug bound in the ecto site: NPC1L1 bound to an ezetimibe analogue, Eze-PS (PDB ID: 6v3h) ^32^. The binding site of Eze-PS in NPC1L1 is topologically similar to PfNCR1, although without a direct overlap.

The binding site of Compound 2 bears some remote similarity with the ligand-bound state of NPC1L1 in the presence of the ezetimibe analogue (ezetimibe-PS) (PDB ID: 6v3h) ^32^. In PfNCR1, the corresponding residues forming the ecto site are located closer to the membrane than those in NPC1L1, and the relative ligand binding site locations do not overlap (**Figure 3D**).

### Structure of the Compound 3-bound PfNCR1

The cryo-EM reconstruction of PfNCR1 bound to Compound 3 revealed several striking aspects of ligand binding. First, the ligand is bound both to the ecto site and the neck site (**Figure 4A**). Remarkably, binding to the ecto site does not displace the sterol molecule. Instead, Compound 3 and the sterol bind to the protein together in proximity, suggesting cooperative interactions and a tight connection between the sterol and the ecto site (**Figure 4B**). The binding pockets of Compound 2 and Compound 3 within the ecto site are lined by the overlapping set of residues from the ECD1 (R90, S363, M398, V426) and ECD2 (F1129, F1132, F1266). Some of the ECD2 residues in close contact with Compound 2, such as A1208, F1213, L1246, directly interact with the sterol in the Compound 3-bound structure.

**Figure 4.**
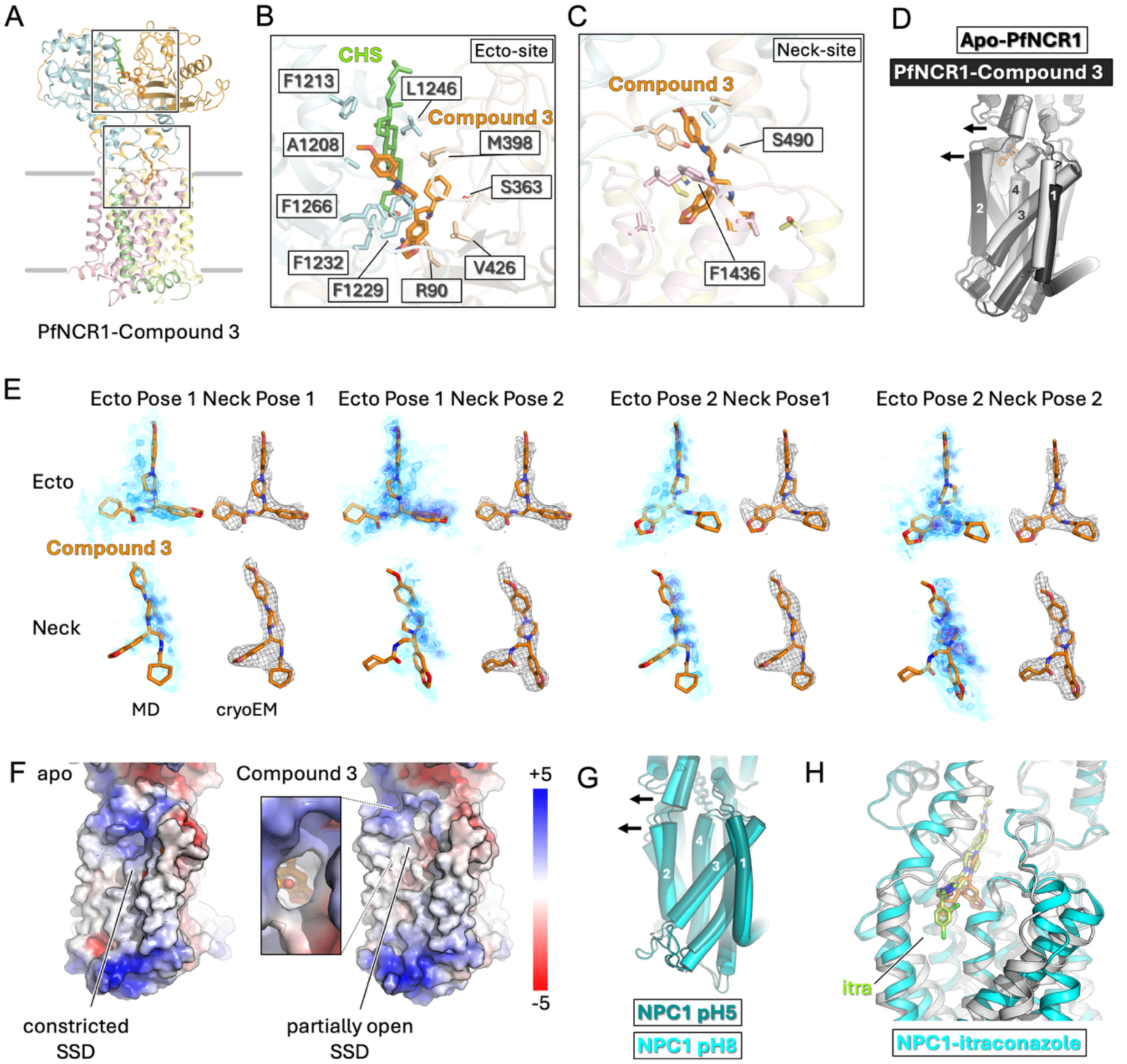
Cryo-EM structure of the PfNCR1-MMV019662 (Compound 3) complex. **A.** Model of PfNCR1-Compound 3. **B.** Close-up look at the ecto site where we can see cooperative binding of a CHS molecule in green and the Compound 3 in orange with neighboring residues. **C.** Close-up look of the neck site with Compound 3 in orange and the mutation sites S490 and F1436 labeled. **D.** The views of apo-PfNCR1 (white tubes) and Compound 3-bound. Arrows indicate the remodeling of the SSD in the presence of drug due to the shift in the TM2 and the neck site forming helices. **E.** Comparison between the cryo-EM densities and the MD-based densities of Compound 3, showing in gray mesh representation the cryo-EM density, highlighting the suggested pose of the ligand in the stick representation, while the MD-based densities are shown as a blue volume representation reflecting percent occupancy. **F.** Electrostatic surface map comparison of apo-and Compound 3-bound PfNCR1 and NPC1 shows the radical change in the properties of the lipid bilayer-exposed SSD surfaces of the two proteins (the red-blue gradient bar shows a scale of-5 to +5 kT/e). **G**. Cryo-EM structures of NPC1 in GDN at pH 8 (teal, PDB ID: 6w5s) ^24^, pH 5.5 (cyan, PDB ID: 6w5t) ^24^ show pH-dependent SSD remodeling; the surface of SSD (right panels) shows “open” SSD at pH 8 and “constricted” SSD at pH 5. Numbers 1-4 indicate TM helices 1-4 that participate in the SSD protein-lipid interface. **H.** Comparison of PfNCR1-Compound 3 to the NPC1-itraconazole complex (PDB ID: 6uox) ^38^ shows overlapping and distinct modes of neck site targeting.

As was the case for Compound 1, the mutations altering the sensitivity of PfNCR1 to Compound 3, S490L and F1436I ^18^, perfectly match the neck binding site (**Figure 4C**). Modeling of Compound 3 was also guided by MD simulations, which revealed that only one of two potentially fitting ligand poses in the ecto and neck sites, respectively, was stable (**Figure 4E).** This is highlighted in the 2D RMSD plots (Figure S16-17), which shows only one combination having low difference between the repeats. The individual RMSD were less informative (between 3.02 and 4.89 Å), because of the confined space limited by PfNCR1, restraining possible deviations.

### Compound 3 induces conformational changes in the SSD

Binding of Compound 3 to the neck site is accompanied by a lateral rearrangement of the TM2 relative to the apo-protein, leading to an opening in the SSD (**Figure 4D**). The change leads to a substantial modification of the surface of the SSD that is presumed to interact with the sterols, creating a lateral opening that connects the neck site with the outer leaflet of the membrane (**Figure 4F**). These conformational changes also expose the sidechain of E558 to the membrane and produce a notable alteration in the electrostatic surface potential at the putative entry site for lipids or cholesterol at the SSD surface corresponding to the lateral opening formed upon displacement of TM2 and the neck site. Such a strong reduction of the surface potential could modify the local membrane environment by changing interactions with the lipid, must importantly modifying changing the attraction of the phosphate moieties of the lipid headgroups and contribute to the ability of SSD to serve as a conduit for the passage of hydrophobic molecules (cholesterol or other sterols).

The Compound 3-induced changes in PfNCR1 may reflect the intrinsic plasticity of its SSD, hinting at potential similar rearrangements that may occur in other SSD-containing membrane proteins. Indeed, a similar mode of structural changes in the SSD was proposed for NPC1 in response to pH changes (**Figure 4G**). NPC1 has been suggested to shift from an “open” SSD conformation at high pH (TM2-out, PDB ID: 6w5s), to a “constricted” SSD at pH 5.5 (TM2-in, PDB ID: 6w5t) ^24^. This reorganization was suggested to be linked to pH-dependent steps of cholesterol translocation ^24^. Interestingly, the exact nature of the conformational changes in NPC1 may be more complex than previously suggested, as the first X-ray structure of NPC1 lacking TM0 and the N-terminal domain was determined at low pH and featured an open SSD (PDB ID: 5u73) ^37^.

A cryo-EM structure of NPC1 in the presence of itraconazole (PDB ID: 6uox) ^38^ showed an open SSD occupied by an elongated inhibitor molecule essentially connecting all the way from the protein-lipid interface to the neck site (**Figure 4H**). The Compound 3-bound state is analogous to this state and to the open SSD state of the NPC1, whereas apo-PfNCR1 and the Compound 1-and Compound 2-bound structures feature the constricted SSD as seen in the cryo-EM structures of NPC1 at low pH ^24^.

### *In silico* search for new compounds targeting PfNCR1

Building on the successful structure elucidation, we selected four commercially available analogs of Compound 3 for testing (Table S2). We argue that chemical similarity to Compound 3, a small molecule that is targeted to both the ecto and the neck site, could manifest in altered ligand binding properties that could be evaluated to further characterize these two binding sites (Figure S18A).

Additionally, we docked over 5 million compounds purchasable from MolPort (www.molport.com) to the ecto site and the neck site of apo and holo PfNCR1 structures with the aim to find potential additional ligands (see Materials and Methods for details). After considering the physicochemical properties and the predicted protein-ligand interactions, we selected 12 top-ranked compounds from the hit lists for purchasing and experimental evaluation (Figure S18A, Table S2).

To validate the interactions of the selected compounds with PfNCR1, we assessed their ability to thermostabilize the purified protein. The single-concentration thermostability assays showed that some of the compounds have a slight stabilizing effect (∼1-2°C), whereas others show an apparent destabilizing effect (Figure S18B). Tryptophan fluorescence quenching-based binding assays showed that all these compounds were capable, to a varied degree, of quenching the intrinsic Trp fluorescence of PfNCR1 (Figure S18C).

In addition to testing the effects on protein stability and Trp quenching, we carried out biological activity tests to prioritize the compounds and to determine their ability to kill *P. falciparum*. Growth inhibition assays were performed identically as for the three MMV compounds. Six out of 12 molecules impaired *P. falciparum* asexual growth (Figure S18D): G856-4236, G856-4269, F2573-0329 and F2573-0310 (selected based on ligand similarity), and V029-3356 and STL441565 (selected based on compound library docking). All the compounds showed EC_50_ values in the micromolar range (comprised between 3.56 and 13.40 µM). Thus, these six compounds were selected for further characterization by cryo-EM.

### Structures of PfNCR1 bound to new compounds

The grids with PfNCR1 in complex with the selected compounds were prepared as described for the MMV compounds and subjected to cryo-EM data collection and analysis. The cryo-EM datasets allowed for the reconstruction of cryo-EM maps for six PfNCR1 samples in the presence of the selected compounds (**Figure 5A**). The resolution of each of these datasets was limited to ∼4-6 Å. In five of the 3D reconstructions, the only ligand observed was CHS bound to the ECD sterol site. In contrast, for the G856-4236 (hereafter referred to as Compound 4), we observed a clear density reminiscent of Compound 3 (**Figure 5B**, Figure S19-20).

**Figure 5.**
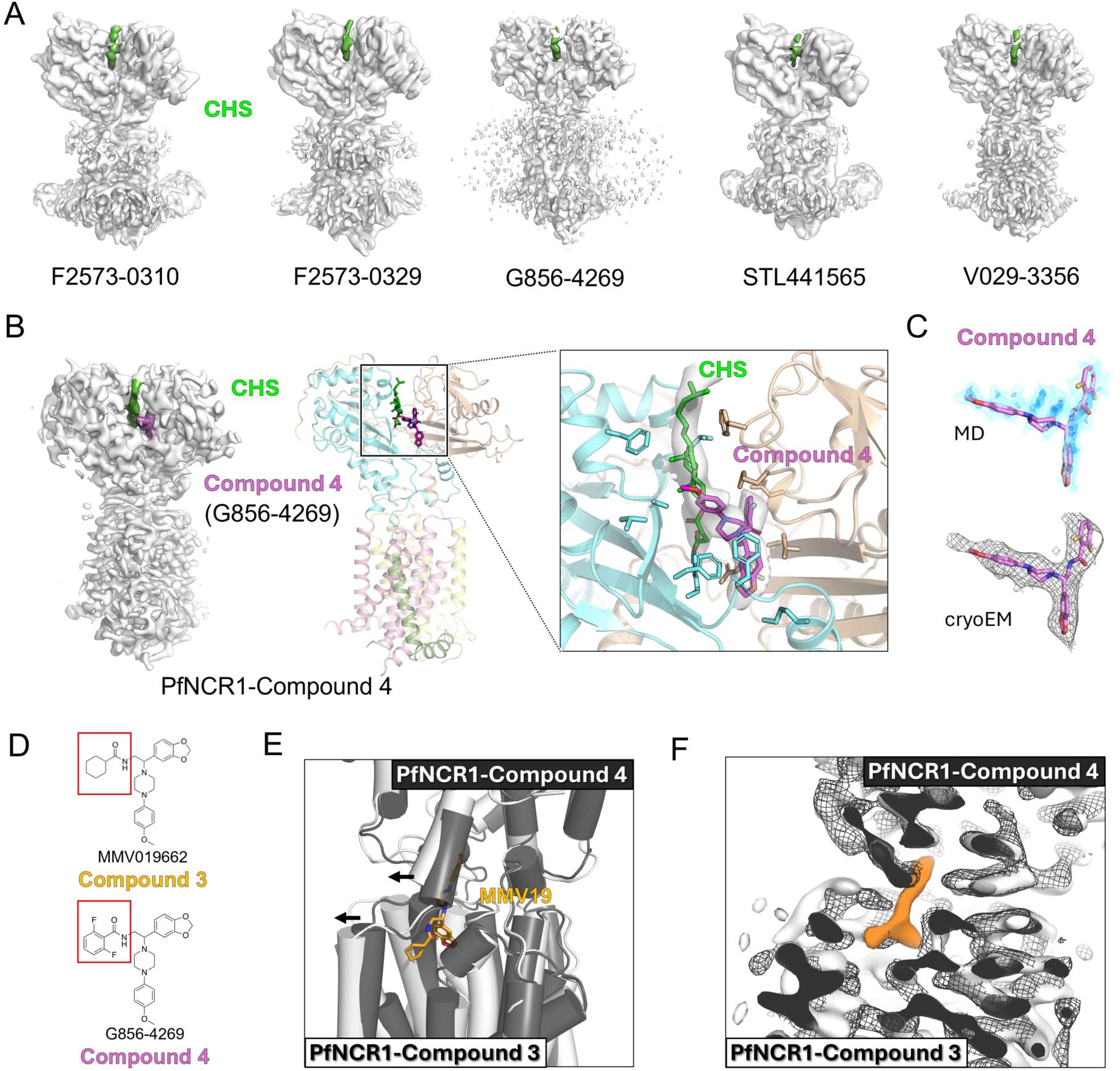
Cryo-EM reconstructions of PfNCR1 in the presence of the six new compounds with biological activity. **A.** Maps of the five complexes with highlighted CHS densities. **B.** Map and model of PfNCR1-G856-4236 (Compound 4) that showed an extra density in the ecto domain. Close-up look of the cooperative binding of the CHS molecule in green and Compound 4 in purple. **C.** Comparison between the cryo-EM density and the MD-based density of Compound 4, showing in gray mesh representation the cryo-EM density, the suggested pose of the ligand in the stick representation, while the MD-based density is shown as a blue volume representation reflecting percent occupancy. **D.** Comparison of the chemical structures of Compound 3 and Compound 4, highlighting the moiety that differs. **E.** Comparison of the neck site of PfNCR1-Compound 3 in gray and PfNCR1-Compound 4 in black showing that Compound 4 does not induce SSD remodeling. **F.** Comparison of the neck site of PfNCR1-Compound 3 map in gray surface and PfNCR1-Compound 4 in black mesh showing that Compound 4 does not bind in the neck site.

As in the case of the Compound 3, the new compound was bound to the ecto site without displacing the sterol (**Figure 5B**). The cryo-EM map was sufficiently well-resolved to allow us to propose a single orientation for Compound 4, which was subsequently confirmed by MD simulations (**Figure 5C**). The geometry of the ligand binding site is very similar, with all the key residues forming a tight pocket around the compound (RMSD: 2.85±0.80 Å). Interestingly, the single modified moiety in Compound 4 (relative to Compound 3) (**Figure 5D**) dips into the groove formed by residues E396, M398, N361, S363, V426, P428. Unlike Compound 3, Compound 4 does not enter the neck site and does not remodel the SSD (**Figure 5E, F**).

We could observe clear density only for one of the six candidate compounds, while all six showed effects in the biophysical and activity assays. The lack of observed cryo-EM density could be due to a variety of factors, from suboptimal conditions for protein-ligand interactions *in vitro*, to low affinity of these compounds insufficient to compete the CHS out of its binding site, to high dynamics of binding hampering direct detection of interaction by cryo-EM.

### Molecular dynamics (MD) simulations of the PfNCR1 cryo-EM structures

To complement the structural analysis, we performed molecular dynamics (MD) simulations of the cryo-EM structure of PfNCR1 in complex with CHS and compounds 1-4. For each ligand we evaluated all possible poses as inferred from the cryo-EM densities, performing 300 ns long simulations of 5 repeats for each pose. A comparison of the explored spatial volume of the ligand as probability density allowed to directly compare the ligand dynamics with the density determined by cryo-EM, revealing the correct binding poses of PfNCR1-bound compounds (**Figure 2-5**). To measure the openness of the SSD domain we analyzed the distance between residues Cα 559 and Cα 494. This revealed that the compound 3-bound structure consistently exhibited an increased distance compared to the apo and other compound-bound systems, indicating a more open SSD conformation (**Figure 6A, B**). To characterize the global motions of PfNCR1 with the largest amplitude and determine the impact of bound compounds on these motions, we performed principal component analysis (PCA). Combining all simulations allow identifying the common largest motions and to directly compare all simulation within a common PCA framework. The first two principal components (PC1 and PC2) accounted for over 60% of the total variance (Figure S21A) and revealed two main motions: the bending and opening motion corresponding to PC1 and the twisting motion corresponding to PC2 (Figure S21B). Projecting the simulations for each system onto the common PC1 and PC2 (**Figure 6C**) revealed that compound 1 and 2 showed motions that are overlapping with the dynamics of the CHS-only bound PfNCR1. The cryo-EM structure of compound 3 bound PfNCR1 (**Figure 4**) revealed that binding of this compound induced a conformational change, and consistently, MD simulations showed the largest amplitude for compound 3 and the exploration of a conformational space representing a more open conformation in comparison to the CHS-only bound PfNCR1 (**Figure 6C**). Interestingly, the trajectories of compound 4-bound PfNCR1 structures showed a mixed dynamics in which motions of the ECD1 sampled both the open-like and the closed conformation, while the SSD did not show an opening as observed for compound 3, indicating that compound 4 might uncouple motions of the SSD and of the ECD1.

**Figure 6.**
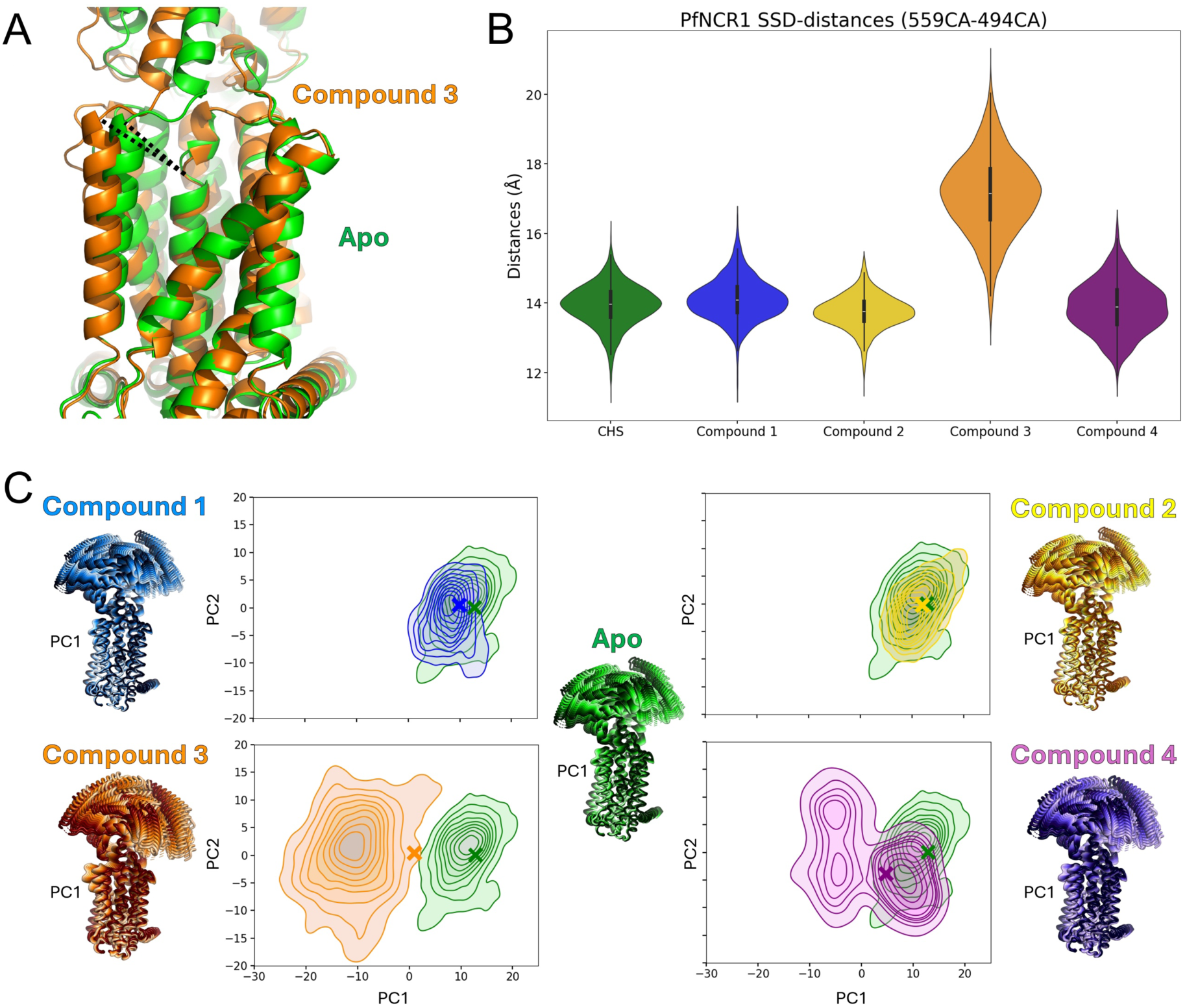
Molecular dynamics simulations of PfNCR1 bound to CHS and compounds 1-4. **A.** Comparison of cryo-EM structures of PfNCR1 when bound to CHS (green) and when bound to compound 3 (orange), focusing on the SSD domain and the distance between residues 494 and 559. **B.** Violin plots of the SSD-distances (distance between Cα atoms residues 559 and 494) during the simulations of PfNCR1 bound to different ligands. **C.** The bending motion described by PC1 for PfNCR1 and PCA 2D plots of the MD simulations of PfNCR1 bound to different ligands, comparing the dynamics of bound-CHS PfNCR1 (in green) and binding of compound 1 (in blue), compound 2 (in yellow), compound 3 (in orange) and compound 4 (in purple).

### Conformation dependency of residue protonation

The cryo-EM structures of the PfNCR1 revealed two conformations showing an open SSD domain as well as a closed SSD domain. A key difference between these two structures is the opening of a small gap between TM2 and TM4. The small gap indicates a potential entry site for cholesterol accessing a conduit leading to the upper end of the ECD domain, where we identified bound CHS (**Figure 2-5**). The inhibitors binding to the neck domain obstruct the initial part of the conduit, while inhibitors bound to the ECD bind to the distal part of the conduit. In the closed SSD conformation this entry gate to the conduit is sealed towards the membrane and also the initial part of the conduit is obstructed by the repositioning of helix TM2 and of the short helices in the neck region immediately following TM1 and preceding TM2. Importantly, in the beginning of TM4 at its extracellular side, E558 changes its orientation between the 2 conformation states. In the open SSD state, E558 is exposed towards the membrane and contributes to the overall negative electrostatic potential observed in this region (**Figure 1H**). In the closed SSD state, the same E558 is oriented towards TM12 and forms a salt bridge with K1434. This salt bridge stabilizes the position of TM4, thereby contributing to maintain a closed entry gate for cholesterol.

In order to obtain mechanistic insights linking the conformational change of the SSD domain to the proton motive force energizing the transport of cholesterol, we carried out pKa calculations using PROPKA ^39^ (**Table S3**). These predictions suggest that D571 on TM4 would be protonated in both conformations. Interestingly, the protonation states of D570 and H1387 are predicted to be conformation sensitive. In the open SSD conformation, the pKa calculations predicted D570 to be negatively charged, while H1387 was predicted to be double protonated and therefore positively charged. In contrast, in the closed state of the SSD domain, D570 is predicted to be protonated and H1387 single protonated, indicating that both residues would be neutral, therefore suggestion that a proton is transferred between D570 and H1387 with the conformational change. The cryo-EM structure revealed that the largest difference is a change in the side chain orientation of D570. In the open conformation, D570 forms a salt bridge with H1387, while in the closed conformation, the protonated side chain of D570 rotates towards TM11 and forms a hydrogen bond with T1422. These three residues are found in the core of the transmembrane domain approximately halfway through the membrane.

To verify these predictions we carried out 4 sets of simulations: we used for both conformational states (open SSD and closed SSD) both protonation state of D570 and H1387. For each configuration we carried out 5 independent simulations (**Figure 7**) using apo structure of the CHS only bound PfNCR1 for closed SSD state and the apo structure of the Compound 3 bound PfNCR1 for the open SSD state. The simulations revealed that in the open SSD state, D570 formed a stable hydrogen bond with T1422 in the protonated form. This side orientation and interaction is consistent with the conformation observed in the cryo-EM structure of the closed SSD PfNCR1. In the charged form, D570 changed orientation away from T1422 towards H1387 and remained dynamics. Similarly, in the closed SSD conformations, only one protonation state was stable and consistent with the cryo-EM structure. The negatively charged D570 and the positively charged H1387 formed a stable salt bridge, consistent with the side chain orientations observed in the closed SSD cryo-EM structure, while in the second protonation state (double neutral), both D570 and H1387 were very dynamic.

**Figure 7.**
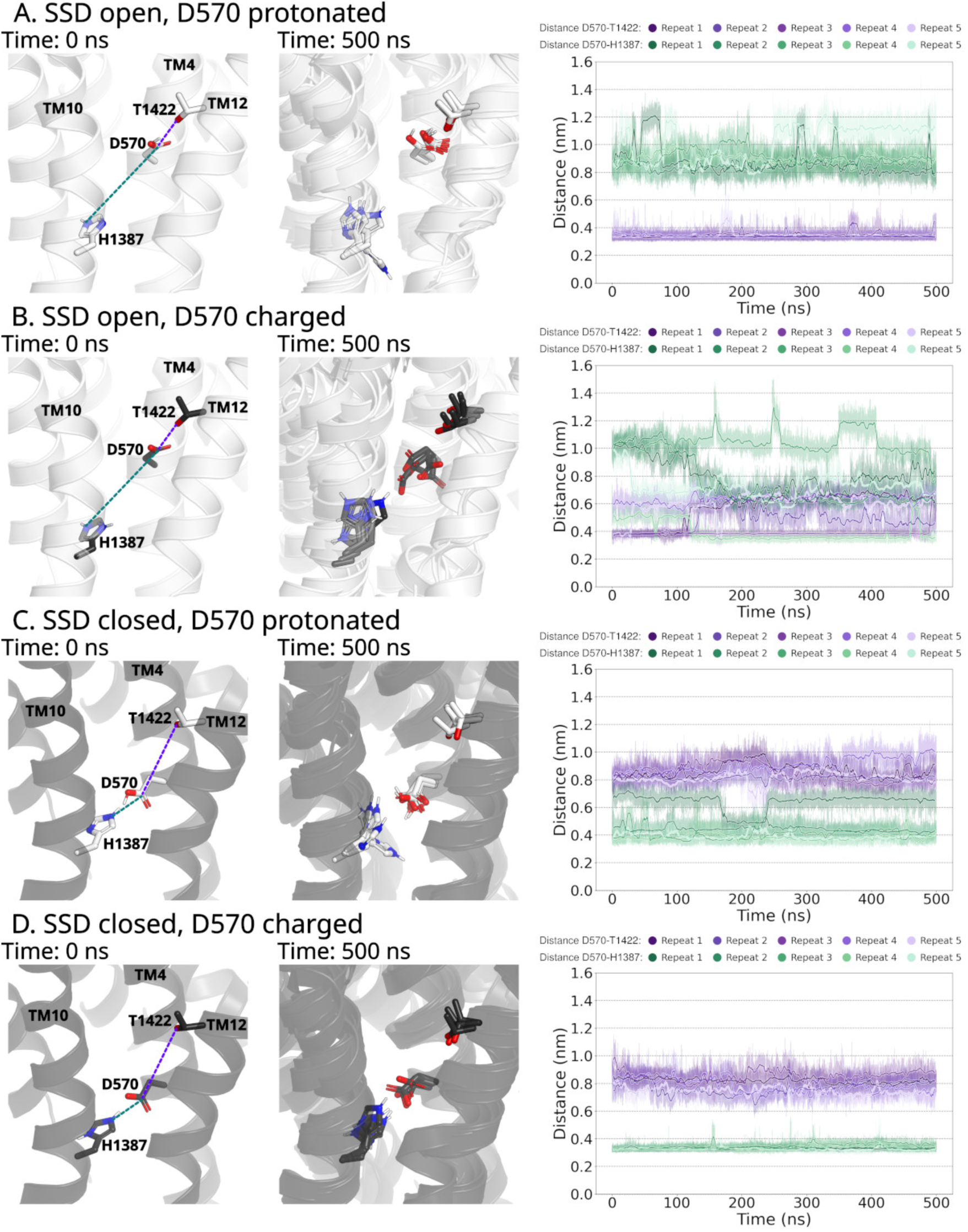
Conformation dependency of D570 and H1387 protonation: **A**. Simulations starting from the open SSD conformation in the apo state using the structure of Compound 3-bound PfNCR1 structure. Residue D570 is protonated, while H1387 is neutral. At 0 ns, the cryo-EM structure-derived starting conformation is shown, at 500 ns - the overlay of 5 independent repeats. The measured distances are indicated by dashed lines. On the right, the distance from the CG carbon of the carboxylate moiety of D570 and the oxygen of the hydroxyl group of T1422 is shown in violet, distance to the εN nitrogen of H1387 is shown in green. **B**. Simulations starting from the open SSD conformation of PfNCR1 in the apo state using the Compound 3-bound state as the starting structure. Residue D570 is charged, while H1387 is double protonated and charged. Distances are shown as reported in A. **C**. Simulations starting from the closed SSD conformation of PfNCR1 in the apo state using the structures of PfNCR1 bound to CSH only. Residue D570 is protonated, while H1387 is neutral. Distances are shown as reported in A. **D**. Simulations starting from the closed SSD conformation of PfNCR1 in the apo state using the structure only bound to CHS as the starting structure. Residues D570 is charged, while H1387 is double protonated and also charged. Distances are shown as reported in A.

Importantly D570 and E558 are both on TM4, suggesting a potential mechanism linking residue protonation in the core of the transmembrane domain of PfNCR1 with the opening of the cholesterol translocation conduit. Our simulations revealed increased dynamics of TM4, where non-matching protonation state of PfNCR1 led to secondary structure instabilities and slight local rotation of TM4 (Figure S22).

## Discussion

The structures of the ligand-bound forms of PfNCR1 point to a remarkable variety of interaction modes that can be exploited to target an RND transporter (**Figure 8**). The presence of multiple ligand binding sites in PfNCR1 is reminiscent of the observations reported for NPC1 and NPC1L1, where distinct ligand interaction modes could be found for very similar drug molecules (for example, ezetimibe binding in the neck site of NPC1L1 ^40^, or ezetimibe-PS binding to the equivalent of an ecto site in another study ^32^). In our case, PfNCR1 as a model system provides a unique testing ground for exploring the molecular pharmacology of RND transporters, with four unique compounds showing very distinct properties. Even more remarkable is the link between PfNCR1 inhibition and the survival of the parasite. It is clear that the activity of PfNCR1 is tightly linked to the fitness of *P. falciparum*, and our findings provide a strong foundation for the rational development of novel antimalarial agents. The observation that compounds 1–3 also target gametocytogenesis further highlights the importance of this pathway, as these non-replicative stages remain particularly difficult to address within current antimalarial drug pipelines. However, additional studies will be required to elucidate their precise mechanism of action at this stage, as well as to define the cellular requirements for PfNCR1 during gametocytogenesis.

**Figure 8.**
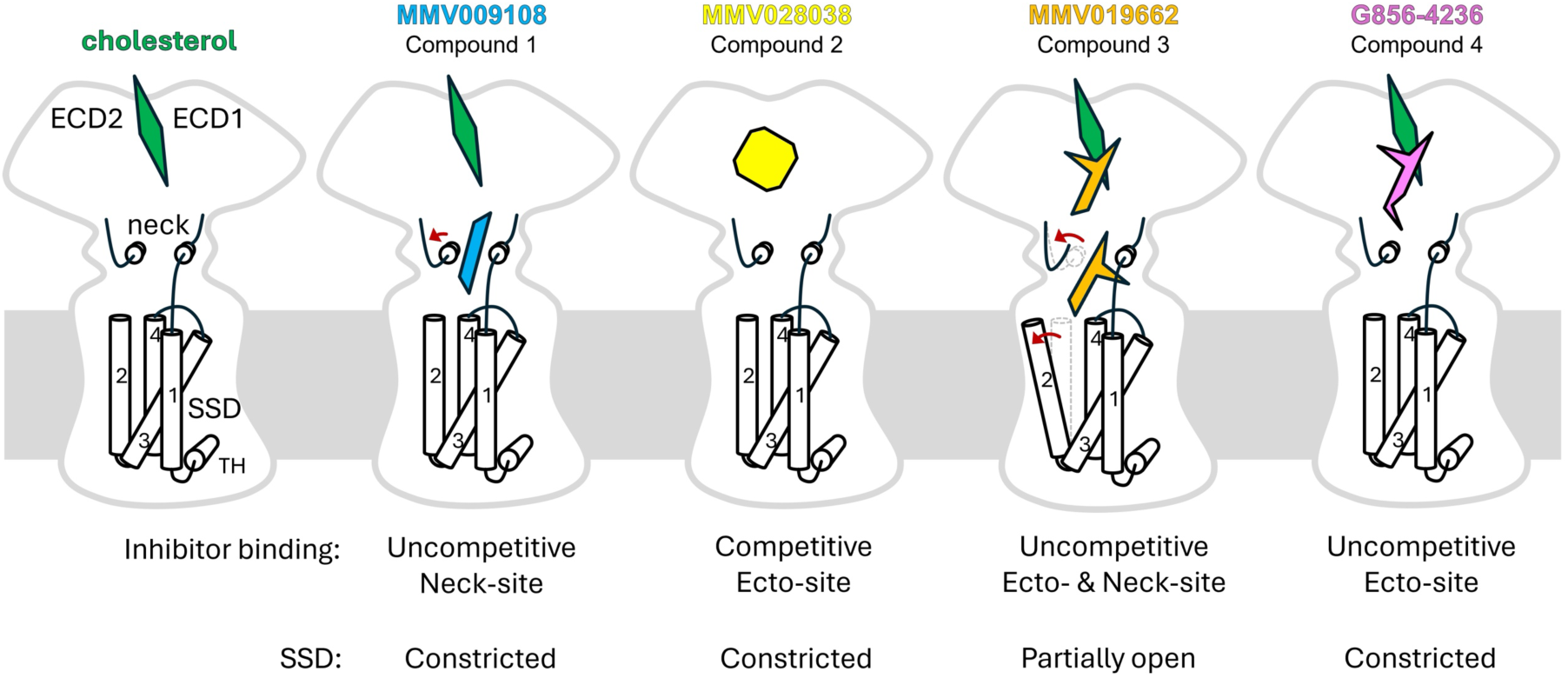
Model of PfNCR1 inhibition via ecto and neck sites, potential role of the SSD remodeling in the inhibition mechanism, potential for drug discovery targeting PfNCR1 and related RND transporters. The sterol bound to the ECD is green, the three MMV compounds are coloured blue (MMV009108, Compound 1), yellow (MMV028038, Compound 2) and orange (MMV019662, Compound 3). The newly identified inhibitor G856-4236 (Compound 4) is shown as a magenta-coloured shape bound close to the sterol. The changes induced by drug binding in the neck-forming residues and in the SSD are illustrated with red arrows.

It is worth noting that the previously determined structures of PfNCR1 in apo-and MMV009108-bound states ^20^ are generally consistent with the structures reported here. For the MMV009108 (here, Compound 1), the MD simulations in our case showed a slight preference of the inverted drug pose, relative to the previously published one. It is likely that a cryo-EM structure and a higher resolution than we and others have been able to achieve will be required to unambiguously define the orientation of MMV009108 in the binding site. Importantly, the previously reported mutations disrupting the effects of Compound 1, as well as Compounds 2 and 3 ^18^, are perfectly in line with the observed cryo-EM densities described here, as detailed in the results section for each of the ligands. The specific molecular effects that each of the mutations exert on the PfNCR1 conformation landscape, as well as on PfNCR1-ligand interactions, will require careful experimental and computational investigation.

The conformational plasticity of the SSD in PfNCR1 puts a spotlight on the SSDs in the vertebrate homologues. Our observations of Compound 3-induced opening of the SSD are paralleled by the studies of NPC1 that revealed a potential pH-dependent switch of the SSD conformation. It appears that interaction with Compound 3 alone (but not its close relative identified in our study, Compound 4) is sufficient to induce the open SSD rearrangement, linking the neck site with the SSD surface exposed to the lipids. The transition between the open and the closed SSD conformation is associated with a change of side chain orientation of D570, interacting with H1387 on TM10 in the open SSD conformation and with T1422 on TM11 in the closed SSD conformation. This change in the interaction pattern is associated with a transfer of a proton between D570 and H1387, hinting towards a link between the conformational transition of the SSD, the opening of the cholesterol conduit and the proton motive force energizing transport.

It is conceivable, that the local structural changes of D570 propagate along TM4 towards the extracellular side, where E558 forms a salt bridge with K1434, contributing to stabilize the closed SSD state, while in the open SSD state, E558 orients toward the membrane, potentially destabilizing phospholipids juxtaposed to the entry gate of the cholesterol transfer tunnel. Consistently, we find that the conformational transition is accompanied by a dramatic change in the surface properties of the SSD, essentially collapsing the local positive electrostatic surface potential. This may be directly linked to the mode of sterol translocation by PfNCR1 and other RND proteins, such as Patched/PTCH1, Dispatched/DISP1, NPC1 and NPC1L1. For NPC1, the conduit formed by the SSD opening at high pH has been interpreted as part of the translocation pathway for sterols ^24^. It remains unclear whether the same mechanism would apply to other vertebrate RND transporters, for which a variety of the SSD conformations have not yet been observed. In the case of PfNCR1, the opening of the SSD by Compound 3 binding may be related to one of two potential scenarios: (i) Compound 3 may be mimicking a sterol entering the neck site and opening up the SSD by “pushing” the TM2 aside; (ii) Compound 3 may induce a rearrangement of the SSD structure that perturbs the movement of the sterol at the SSD surface. The ability of Compound 1 to block PfNCR1 activity by acting on the neck site without perturbing the SSD lends some support to the first option. However, because the path of cholesterol movement remains to be determined in PfNCR1 as well as in other RND proteins, neither one of the two scenarios should be excluded.

It is noteworthy that the equivalent residues in other RND transporters, from *E. coli* AcrB ^41^, to *S. cerevisiae* NCR1 ^42^, to human NPC1 ^24^, have been shown to be important for proton coupling of sterol transport (in the case of AcrB, for multi-drug transport activity). Remarkably, the integrated analysis including cryo-EM structures, MD simulations and PROPKA-based protonation predictions provides unbiased evidence for direct coupling between the protonation state of the D570-H1387 pair in the TM core with the open/closed conformation of the SSD. Despite being agnostic to the published literature on pH-dependent RND transporters, the PROPKA analysis pinpointed two residues in the TM5-TM10 region PfNCR1 that precisely match established proton motive force-responsive residues in RND transporters involved in sterol or drug transport (Figure S23). Achieving this with relatively simple tools as described here is a highly encouraging and suggests that the emerging machine learning approaches will soon yield profound mechanistic insights into mechanisms of pH-dependent membrane proteins.

It is conceptually easier to understand how drugs targeting the ecto site operate. Compound 2 displaces the sterol molecule and would thus prevent the passage of sterols, while Compound 3 and Compound 4 seem to cooperatively bind alongside the sterol molecule at their target binding site, likely stabilizing the sterol and thus forming a roadblock in the sterol translocation pathway. This latter mode of inhibition has not been observed in RND transporters to date. It could provide an important hint to the drug discovery efforts by suggesting a novel mode of engagement with the target transporter of general applicability. The mode of inhibitor binding in close proximity to the sterol binding site or in direct contact with the bound sterol could be potentially relevant not only to PfNCR1, but also to the human RND transporters, e.g. PTCH1 or NPC1L1, as drug discovery efforts aimed at identifying inhibitors of these proteins in cancer (PTCH1) and in hypercholesterolemia (NPC1L1) are underway. The possibility of exploiting the novel inhibition modality described here, involving a small molecule that binds to its binding site cooperatively with a sterol molecule, may provide new avenues for therapeutic development.

## Acknowledgements

We thank Thorsten Blum (PSI) for assistance with cryo-EM data analysis. We also thank Greta Assmann and Joao Pedro Agostinho de Sousa (PSI) for support in high-performance computing. We thank Dominique Soldati for the anti-actin antibody and Daniel E. Goldberg for the PfNCR1-GFP parasites. This work was supported by the Swiss National Science Foundation (CRSII5_198545 and 320030-236267) to MB, as well as Swiss National Science Foundation grant 184951 to VMK. CB is supported by a SNSF Swiss Postdoctoral Fellowship #217028. MAM is supported by the FWF grant PAT9543323 to TS.

## References

1 World malaria report 2025: addressing the threat of antimalarial drug resistance. Geneva: World Health Organization; 2025. Licence: CC BY-NC-SA 3.0 IGO.

2 Venugopal, K., Hentzschel, F., Valkiunas, G. & Marti, M. Plasmodium asexual growth and sexual development in the haematopoietic niche of the host. Nat Rev Microbiol 18, 177–189 (2020). 10.1038/s41579-019-0306-2

3 Duffy, P. E., Gorres, J. P., Healy, S. A. & Fried, M. Malaria vaccines: a new era of prevention and control. Nat Rev Microbiol 22, 756–772 (2024). 10.1038/s41579-024-01065-7

4 Siqueira-Neto, J. L. et al. Antimalarial drug discovery: progress and approaches. Nat Rev Drug Discov 22, 807–826 (2023). 10.1038/s41573-023-00772-9

5 Hamerly, T. et al. NPC1161B, an 8-Aminoquinoline Analog, Is Metabolized in the Mosquito and Inhibits Plasmodium falciparum Oocyst Maturation. Front Pharmacol 10, 1265 (2019). 10.3389/fphar.2019.01265

6 Birkholtz, L. M., Alano, P. & Leroy, D. Transmission-blocking drugs for malaria elimination. Trends Parasitol 38, 390–403 (2022). 10.1016/j.pt.2022.01.011

7 Wong, W. et al. Mefloquine targets the Plasmodium falciparum 80S ribosome to inhibit protein synthesis. Nat Microbiol 2, 17031 (2017). 10.1038/nmicrobiol.2017.31

8 Fry, M. & Pudney, M. Site of action of the antimalarial hydroxynaphthoquinone, 2-[trans-4-(4’-chlorophenyl) cyclohexyl]-3-hydroxy-1,4-naphthoquinone (566C80). Biochem Pharmacol 43, 1545–1553 (1992). 10.1016/0006-2952(92)90213-3

9 Ribbiso, K. A. et al. Artemisinin-Based Drugs Target the Plasmodium falciparum Heme Detoxification Pathway. Antimicrob Agents Chemother 65 (2021). 10.1128/AAC.02137-20

10 Wang, J. et al. Haem-activated promiscuous targeting of artemisinin in Plasmodium falciparum. Nat Commun 6, 10111 (2015). 10.1038/ncomms10111

11 Bridgford, J. L. et al. Artemisinin kills malaria parasites by damaging proteins and inhibiting the proteasome. Nat Commun 9, 3801 (2018). 10.1038/s41467-018-06221-1

12 Mbengue, A. et al. A molecular mechanism of artemisinin resistance in Plasmodium falciparum malaria. Nature 520, 683–687 (2015). 10.1038/nature14412

13 Bailly, C. Pyronaridine: An update of its pharmacological activities and mechanisms of action. Biopolymers 112, e23398 (2021). 10.1002/bip.23398

14 Carrington, H. C., Crowther, A. F., Davey, D. G., Levi, A. A. & Rose, F. L. A metabolite of paludrine with high antimalarial activity. Nature 168, 1080 (1951). 10.1038/1681080a0

15 Thapar, M. M., Gupta, S., Spindler, C., Wernsdorfer, W. H. & Bjorkman, A. Pharmacodynamic interactions among atovaquone, proguanil and cycloguanil against Plasmodium falciparum in vitro. Trans R Soc Trop Med Hyg 97, 331–337 (2003). 10.1016/s0035-9203(03)90162-3

16 Srivastava, I. K. & Vaidya, A. B. A mechanism for the synergistic antimalarial action of atovaquone and proguanil. Antimicrob Agents Chemother 43, 1334–1339 (1999). 10.1128/AAC.43.6.1334

17 Ross, L. S. & Fidock, D. A. Elucidating Mechanisms of Drug-Resistant Plasmodium falciparum. Cell Host Microbe 26, 35–47 (2019). 10.1016/j.chom.2019.06.001

18 Istvan, E. S. et al. Plasmodium Niemann-Pick type C1-related protein is a druggable target required for parasite membrane homeostasis. Elife 8 (2019). 10.7554/eLife.40529

19 Garten, M. et al. Contacting domains segregate a lipid transporter from a solute transporter in the malarial host-parasite interface. Nat Commun 11, 3825 (2020). 10.1038/s41467-020-17506-9

20 Zhang, Z. et al. The Plasmodium falciparum NCR1 transporter is an antimalarial target that exports cholesterol from the parasite’s plasma membrane. Sci Adv 10, eadq6651 (2024). 10.1126/sciadv.adq6651

21 Kobylka, J., Kuth, M. S., Muller, R. T., Geertsma, E. R. & Pos, K. M. AcrB: a mean, keen, drug efflux machine. Ann N Y Acad Sci 1459, 38–68 (2020). 10.1111/nyas.14239

22 Doughty, D. M. et al. The RND-family transporter, HpnN, is required for hopanoid localization to the outer membrane of Rhodopseudomonas palustris TIE-1. Proc Natl Acad Sci U S A 108, E1045–1051 (2011). 10.1073/pnas.1104209108

23 Gong, X. et al. Structural Insights into the Niemann-Pick C1 (NPC1)-Mediated Cholesterol Transfer and Ebola Infection. Cell 165, 1467–1478 (2016). 10.1016/j.cell.2016.05.022

24 Qian, H. et al. Structural Basis of Low-pH-Dependent Lysosomal Cholesterol Egress by NPC1 and NPC2. Cell 182, 98–111 e118 (2020). 10.1016/j.cell.2020.05.020

25 Long, T., Liu, Y., Qin, Y., DeBose-Boyd, R. A. & Li, X. Structures of dimeric human NPC1L1 provide insight into mechanisms for cholesterol absorption. Sci Adv 7 (2021). 10.1126/sciadv.abh3997

26 Zhang, Y. & Beachy, P. A. Cellular and molecular mechanisms of Hedgehog signalling. Nat Rev Mol Cell Biol 24, 668–687 (2023). 10.1038/s41580-023-00591-1

27 Qi, C., Di Minin, G., Vercellino, I., Wutz, A. & Korkhov, V. M. Structural basis of sterol recognition by human hedgehog receptor PTCH1. Sci Adv 5, eaaw6490 (2019). 10.1126/sciadv.aaw6490

28 Burke, R. et al. Dispatched, a novel sterol-sensing domain protein dedicated to the release of cholesterol-modified hedgehog from signaling cells. Cell 99, 803–815 (1999). 10.1016/s0092-8674(00)81677-3

29 Wu, X., Yan, R., Cao, P., Qian, H. & Yan, N. Structural advances in sterol-sensing domain-containing proteins. Trends Biochem Sci 47, 289–300 (2022). 10.1016/j.tibs.2021.12.005

30 Yang, T. et al. Crucial step in cholesterol homeostasis: sterols promote binding of SCAP to INSIG-1, a membrane protein that facilitates retention of SREBPs in ER. Cell 110, 489–500 (2002). 10.1016/s0092-8674(02)00872-3

31 Kuwabara, P. E. & Labouesse, M. The sterol-sensing domain: multiple families, a unique role? Trends Genet 18, 193–201 (2002). 10.1016/s0168-9525(02)02640-9

32 Huang, C. S. et al. Cryo-EM structures of NPC1L1 reveal mechanisms of cholesterol transport and ezetimibe inhibition. Sci Adv 6, eabb1989 (2020). 10.1126/sciadv.abb1989

33 Cannac, F. et al. Cryo-EM structure of the Hedgehog release protein Dispatched. Sci Adv 6, eaay7928 (2020). 10.1126/sciadv.aay7928

34 Lasonder, E. et al. Integrated transcriptomic and proteomic analyses of P. falciparum gametocytes: molecular insight into sex-specific processes and translational repression. Nucleic Acids Res 44, 6087–6101 (2016). 10.1093/nar/gkw536

35 Silvestrini, F. et al. Protein export marks the early phase of gametocytogenesis of the human malaria parasite Plasmodium falciparum. Mol Cell Proteomics 9, 1437–1448 (2010). 10.1074/mcp.M900479-MCP200

36 Jumper, J. et al. Highly accurate protein structure prediction with AlphaFold. Nature 596, 583–589 (2021). 10.1038/s41586-021-03819-2

37 Li, X. et al. 3.3 A structure of Niemann-Pick C1 protein reveals insights into the function of the C-terminal luminal domain in cholesterol transport. Proc Natl Acad Sci U S A 114, 9116–9121 (2017). 10.1073/pnas.1711716114

38 Long, T. et al. Structural basis for itraconazole-mediated NPC1 inhibition. Nat Commun 11, 152 (2020). 10.1038/s41467-019-13917-5

39 Olsson, M. H., Sondergaard, C. R., Rostkowski, M. & Jensen, J. H. PROPKA3: Consistent Treatment of Internal and Surface Residues in Empirical pKa Predictions. J Chem Theory Comput 7, 525–537 (2011). 10.1021/ct100578z

40 Hu, M. et al. Structural insights into the mechanism of human NPC1L1-mediated cholesterol uptake. Sci Adv 7 (2021). 10.1126/sciadv.abg3188

41 Murakami, S., Nakashima, R., Yamashita, E. & Yamaguchi, A. Crystal structure of bacterial multidrug efflux transporter AcrB. Nature 419, 587–593 (2002). 10.1038/nature01050

42 Frain, K. M. et al. Conformational changes in the Niemann-Pick type C1 protein NCR1 drive sterol translocation. Proc Natl Acad Sci U S A 121, e2315575121 (2024). 10.1073/pnas.2315575121

