## Supplementary Information for "Structural basis for multiple binding modes of antimalarial ligands targeting *Plasmodium falciparum* NCR1"

### Materials and Methods

#### Protein expression and purification

*Plasmid Constructs:* Plasmid constructs were synthesized and cloned by Genewiz (Azenta Life Sciences). The target gene sequences were codon-optimized for expression in human cells and synthesized de novo. Cloning was performed into the pAYST vector, which includes a C-terminal 3C protease cleavage site, a yellow fluorescent protein (YFP) tag, and a twin-strep tag for downstream purification and detection.

*GFP-nanobody:* The expression and purification of anti-GFP nanobody was done as previously described [1]. In brief, the anti GFP nanobody plasmid was transformed in E. coli BL21(DE3) cells and grown in terrific broth (TB) medium supplemented with 100 µg/mL ampicillin at 37°C. The expression was induced at OD 600 of 0.8 using 500ul of 1mM IPTG at 20 °C for 16 hours. Cells were harvested at 4000 g at 4 °C for 20 minutes and stored at -80 °C. For purification, cell pellets were thawed and resuspended in 20mM HEPES, pH 7.5, 150mM NaCl, 5% glycerol, and 1mM PMSF. Cells were lysed using a microfluidizer LM10 at 18000 psi for three cycles. Cell debris was removed by centrifuging at 19000 rpm at 4 °C for 45 minutes. The supernatant was collected and incubated with pre-equilibrated Ni-NTA resin for 30 minutes. 8 ml of bed volume for every 4 liters of culture was used. The Ni-NTA resin was equilibrated with 5 CV of 20mM HEPES pH 7.5, 150mM NaCl. Beads were washed with 10 CV of 20mM HEPES pH 7.5, 150mM NaCl, 20mM imidazole, followed by 10 CV washes with 20mM HEPES pH 7.5, 150mM NaCl, 40mM imidazole. Elution was done by adding 1.5 CV of 20mM HEPES pH 7.5, 150mM NaCl, 250mM imidazole, 10% glycerol for 3-5 times. The elution fractions were pooled and concentrated using a 10kDa cutoff centrifugal filter. Following Ni-NTA purification, GFP-NB is subjected to SEC in aliquots of 5ml at 18-20mg/ml. SEC is done on a Superdex 75 column using 20mM HEPES, pH 7.5, 150mM NaCl. The corresponding fractions were collected, 10% glycerol was added, and the purified protein was flash frozen in aliquots and stored at -80 °C. For coupling, 0.8g of CnBr-activated Sepharose 4B and 10ml of 1mM HCl were added to a 50ml Falcon tube and left rotating until resin beads were homogenously swollen and resuspended. The beads were collected in a 50ml Bio-Rad column and washed with 20 ml of 1mM HCl and 20ml of 0.1M $NaHCO_3$ and 0.5M NaCl at pH 8.5. The beads were resuspended in 15ml of 0.1M $NaHCO_3$, 0.5M NaCl, pH 8.5, and 10mg of anti-GFP NB solution was added and left rolling overnight at 4 °C. The next day, the beads were collected and washed twice with 20 mL of 0.2M glycine-NaOH at pH 8.0 and then washed with 20ml of PBS. Finally, the beads were resuspended in 20ml of PBS.

*HRV3C:* The HRV3C plasmid was expressed in BL21(DE3) cells cultivated in TB medium. Induction was carried out at an OD 600 of 0.6 by adding 100 µL of 1 mM IPTG, followed by incubation at 20°C for 16 hours. The cells were harvested by centrifugation at 4500 rpm for 20 minutes and stored at -80°C.

For purification, the collected cell pellets were resuspended in a buffer containing 25 mM HEPES at pH 7.5, 500 mM NaCl, 10% glycerol, 5 mM β-mercaptoethanol, and 50 mM imidazole. Lysis was performed using a microfluidizer LM10 at 18,000 psi for three cycles. The lysate was cleared by centrifugation at 15,000 rpm for 30 minutes at 4°C. The resulting supernatant was incubated with pre-equilibrated Ni-NTA resin for 30 minutes, utilizing 8 mL of resin bed volume per 4 liters of culture. The Ni-NTA resin was equilibrated with 4 column volumes (CV) of 20 mM HEPES at pH 7.5, 150 mM NaCl. The resin was then washed with 4 CV of a buffer containing 25 mM HEPES at pH 7.5, 500 mM NaCl, 10% glycerol, 5 mM β-mercaptoethanol, and 50 mM imidazole. Elution was performed using 1.5 CV of 25 mM HEPES pH 7.5, 500 mM NaCl, 10% glycerol, 5 mM β-mercaptoethanol, and 500 mM imidazole. The eluted fractions were pooled and desalted using a HiPrep 26/10 Desalting column to exchange the buffer to 25 mM HEPES at pH 7.5, 500 mM NaCl, 10% glycerol, and 5 mM β-mercaptoethanol, achieving a final concentration of approximately 1 mg/mL. Aliquots were flash-frozen and stored at -80°C.

*Adherent cell culture:* Protein expression was done in HEK293F adherent cells maintained in 15cm cell culture plates in Dulbecco's Modified Eagle's Medium (DMEM, BioConcept) medium, complemented with 1% Penicillin-Streptomycin (PenStrep) and 10% Fetal Calf Serum (FCS) at 37°C and 5 % $CO_2$. Before transfection, the medium was exchanged to DMEM supplemented with 2% FCS and 1% PenStrep. Transfection was done using branched polyethylene imine (PEI) using 50 µg of construct DNA per plate and 100 µg of branched PEI per plate (DNA:PEI=1:2) in non-supplemented DMEM medium. The mixture was incubated for 10 minutes to allow DNA-PEI complex formation and then added to the plates in a drop-wise fashion. 5mM NaBu was added 24h post transfection and the cells were harvested by scraping and centrifugation at 1000 g at 4°C for 10 minutes. The cell pellets were frozen and stored at -80°.

*Suspension cell culture:* Protein expression was done in suspension HEK293F cells maintained in 2l Erlenmeyer flasks in Freestyle medium complemented with 1% Penicillin-Streptomycin (PenStrep) and 2% Fetal Calf Serum (FCS) at 37° and 5 % $CO_2$. Transfection was done using linear polyethylene imine (PEI) using 1 g of construct DNA per liter of culture at a density of 1 million cells and 3 g of branched PEI per liter of culture at a density of 1 million (DNA:PEI=1:3) in non-supplemented Freestyle medium. The mixture was incubated for 10 minutes to allow DNA-PEI complex formation and then added to the flasks. 5mM NaBu was added 24h post transfection and the cells were harvested by centrifugation at 1500 g at 4°C for 15 minutes. The cell pellets were frozen and stored at -80°.

*Protein purification YFP- tag*: Cell pellets were thawed on ice and resuspended in buffer A: 50mM Tris-HCl pH 8.0, 150mM NaCl, 10% glycerol, Roche anti-protease inhibitor tablet, 1mM PMSF. Cells were homogenized with a dounce homogenizer (20 strokes). A membrane prep was done by ultracentrifugation for 35 minutes at 40000rpm using a Beckmann Ti45 rotor. The supernatant was discarded, and the pellet was resuspended in Buffer A. For solubilization 1%DDM/0.2%CHS was added and incubated for 60 minutes at 4°C while rolling. Another ultracentrifugation step was done for 40 minutes at 40000 rpm using a Ti45 rotor. The supernatant was collected and then mixed with 0.8g of prepared GFP-NB coupled to Sepharose beads. Incubated for 60 minutes to allow binding at 4 degrees, while occasionally mixing by hand. The solution was collected in a 50ml Bio-Rad column and washed with 130ml of buffer B: 50mM Tris-HCl at pH=8, 150mM NaCl, 10% glycerol, 0.02% GDN. For cleavage, 10ml of buffer B and 0.8mg of HRV3C protease were incubated for 2h on a roller. The flow-through was collected and eluted again with 2ml of buffer B. The elution was concentrated using a 100kDa cutoff centrifugal concentrator by using 2900g 5 minute until a volume of 1ml was reached. A 3-minute high-speed spin was done to remove large aggregates. The concentrated protein solution was injected into a Superose6 column run with buffer C: 50mM Tris-HCl, pH=8, 150mM NaCl, 0.02% GDN (**Figure S1**).

*Protein purification twin-strep tag:* Procedure is very similar to the purification using YFP-tag previously described. Here Strep-tactin XT 4flow resin was used. Instead of cleaving the tag elution with 3CV of buffer 50mM Tris-HCl pH=8, 150mM NaCl, 10% glycerol, 0.02% GDN, 50mM biotin was used.

**Cryo-EM sample preparation and data collection**

*Cryo-EM sample preparation:* Freshly purified protein in 0.02% GDN was concentrated using a 100kDa cutoff centrifugal filter to a final concentration of 4 mg/ml for the Apo sample and 8mg/ml for the drug samples. MMV009019 (Enamine, Compound 1), MMV028038 (Enamine, Compound 2), MMV019662 (Enamine, Compound 3), F2573-0310 (Life Chemicals Inc.), F2573-0329 (Life Chemicals Inc.), STL441565 (Vitas M Chemical Limited), V029-3356 (ChemDiv Inc.), G856-4269 (ChemDiv Inc.), G856-4236 (ChemDiv Inc., Compound 4) were added to separate samples at a final concentration of 1mM and incubated for 10 min on ice. An aliquot of 3.5 µl of sample was placed on the glow-discharged cryo-EM grid (Quantifoil R1.2/1.3 cu 200), blotted, and plunge-frozen in liquid ethane using a Mark IV Vitrobot instrument maintained at 100% humidity with blot force 10 and blot time of 3 seconds. The grids were cryo-transferred for storage in liquid nitrogen.

*Data collection:* The cryo-EM datasets were obtained at the SCOPEM facility at ETHZ using a 300 kV Titan Krios electron microscope (FEI) equipped with a K3 direct electron detector (Gatan) with a pixel size of 0.66 Å/pix (in super-resolution mode) for the apo dataset and 0.65 Å/pix for the drug datasets. The defocus range was set from -0.5 µm to -3 µm, with the movies being dose fractionated into 40 frames. The total dose for the apo dataset was 55 e/Å2, for the Compound 1 dataset it was 53 e/Å2, for the two Compound 2 datasets it was 53 e/Å2 and 57 e/Å2, for the two Compound 3 datasets it was 60 e/Å2, and for the two Compound 4 datasets it was 58 e/Å2 and 50 e/Å2.

#### Cryo-EM data analysis and model building

Data processing was performed in Relion [2] and CryoSPARC [3], with initial steps performed in Relion. Micrographs were motion corrected using MotionCor2 [4]. Micrographs were CTF corrected using CTFFIND4.1 [5]. Particles were autopicked using templates from manual picking. Several 2D classification rounds were performed, followed by 3D classification. The particles from the best 3D classes were used for 3D auto-refinement. A no-align 3D classification job was used to separate more bad particles. After that, another 3D auto refinement followed by CTF refinements and polishing was done. From there, the resulting particles were transferred to CryoSPARC, where non-uniform and local refinement jobs were used to finalize the apo map to 3.04 Å (**Figure S2**). Model building was performed in Coot. The model was refined using real_space_refine in Phenix [6].

The MMV molecule-containing samples were processed in a very similar way (**Figure S3-10**). This resulted in an Compound 1-bound map of 3.89 Å, an Compound 2-bound map of 3.69 Å, and an Compound 3- map of 3.73 Å.

#### Tryptophan quenching

The protein was diluted to a final concentration of 2 µM in a buffer containing 50 mM Tris-HCl at pH 8.0, 150 mM NaCl, 0.02% GDN, and 10% glycerol. The solution was centrifuged at 24,000 g for 5 minutes before being transferred to a quartz glass cuvette. Fluorescence spectra were recorded from 280 nm to 500 nm using a Cary Eclipse fluorescence spectrophotometer, with an excitation wavelength of 280 nm. Titration of the three MMV drugs was performed by sequential addition of 0.2 µL aliquots of increasing drug dilutions, resulting in final concentrations ranging from 85 nM to 238.5 µM for each drug. Emission values at 333 nm were used to determine fluorescence quenching. The resulting data was analyzed using GraphPad Prism.

#### Thermal unfolding

PfNCR1 at a concentration of 0.15 mg/ml was mixed with a serial dilution of small molecules, with concentrations ranging from 98 nM to 100 µM, in a buffer containing 50 mM Tris-HCl pH 8.0, 150 mM NaCl, and 0.02% GDN. The prepared mixtures were loaded into Prometheus NT.48 series nanoDSF Grade High Sensitivity capillaries (Nanotemper) and analyzed for thermal unfolding using the Prometheus Panta system (Nanotemper). The temperature was gradually increased at a rate of 1°C per minute from 25°C to 95°C. The data were processed using Panta Analysis Software to determine the first derivative of the fluorescence ratio (350 nm/330 nm) as a function of temperature. The unfolding transition temperature ($T_m$) was identified as the peak of the first derivative curve for each titration. The resulting Tm values were plotted against the corresponding compound concentrations using GraphPad Prism for further analysis.

***P. falciparum* cultures, gametocyte and growth inhibition assays**

*Parasite lines:* Unless indicated otherwise, the *Plasmodium falciparum* line NF54/DiCre [7] was used. In addition, *P. falciparum* 3D7 with C-terminally GFP-tagged NCR1 was kindly provided by the Goldberg lab [8]. These clonal parasites were maintained with 5 nM WR99210 (Jacobus).

*Parasite cultures: Plasmodium falciparum* parasites were cultured *in vitro* as routinely performed, at 4% hematocrit, in RPMI 1640 / 25 mM HEPES / L-Glutamine (Gibco) complemented with AlbuMAX II (0.5%; Gibco), gentamicin (40 µg/ml; Gibco), hypoxanthine (200 µM; Sigma), and choline chloride (2 mM; Sigma). Parasites were grown in human erythrocytes (obtained from the Bern Transfusion Center), at 37°C, under shaking conditions, in a specific gas mixture (3% $O_2$, 4% $CO_2$ in $N_2$). Ring stage parasites were synchronized with a sorbitol treatment: cells were incubated for 10 min in 5% sorbitol at 37°C, then washed and resuspended in complete RPMI media.

**Gametocytogenesis inhibition assay**

*Gametocytogenesis induction*: Synchronized cultures of ~1-3% old rings (26-30 hours post-invasion) were induced with minimal fatty acid medium (RMPI 1640 / 25 mM HEPES / L-Glutamine (Gibco) complemented with BSA 390 mg/100 mL (AppliChem), and 30 µM Oleic acid (Sigma-Aldrich), 30 µM Palmitic acid (Sigma-Aldrich)). Twenty-four hours after induction (referred to as day 0), parasites were grown with serum medium (RMPI 1640 / 25 mM HEPES / L-Glutamine (Gibco) complemented with 200 µM Hypoxanthine (Sigma), 10% human AB serum, and 40 µg/mL gentamycin (Gibco)). The day after (referred to as Gametocyte day 1, Gamday1), serum medium supplemented with 50 mM N-acetylglucosamine (serum/GlucNac) was used for seven consecutive days to eliminate asexual parasites. On Gamday8, gametocytes were grown with serum medium until Gamday 13, when they are fully mature. Gametocyte cultures were kept on a heating plate to maintain a temperature of 37°C during medium changes.

*Inhibition assay*: Two independent inductions were performed, each containing 10 ml of gametocyte cultures. On GamDay 2, inductions were split into a 24-well plate, with 2 mL per well, and fed with serum/GlucNac and compounds were added at a concentration of 3 x $EC_{90}$ (MMV009108, Compound 1: 3 µM; MMV028038, Compound 2: 15 µM; MMV019662, Compound 3: 15 µM). DMSO at 0.15% served as the control. Gametocytes were treated with these drugs until Gamday13 and gametocytaemia was counted daily from Gamday3 until Gamday13 using Hemacolor-stained thin blood smears using a Leica DM750 bright-field microscope. For each induction, a total of 2,000 cells were counted by two independent researchers. The total gametocytaemia represents the average gametocytaemia from Gamday3 to Gamday13 across both inductions.

**Expression of NCR1 in gametocytes**

*Flow cytometry:* Three cultures of stage V gametocytes (induced as described above) of either NF54 or NCR1-GFP were analysed by flow cytometry to detect GFP fluorescence. In parallel, three cultures of asexual NF54 or NCR1-GFP were also analysed: those were synchronised to capture the different stages (rings, trophozoites and schizonts). 5 µl of cells were incubated 20 min in Hoechst 33342 (Invitrogen; 1:2'000 in incomplete RPMI) at 37°C. Cells were analysed on a CytoFLEX S, with gating as follows: red blood cells were gated on FSC-H / SSC-H, infected cells on Hoechst (PB450-A) / SSC-A, singlets on FSC-Width / FSC-A, and finally histograms of GFP (FITC-A) are shown, with average values indicated. Gating strategies are explained in Figure S2. Note that for the asexual blood stages, gating of ring- and trophozoite-infected cells were the same, and gating of schizont-infected cells was different (higher Hoechst signal). Data was analysed on FlowJo v10.10.1.

*Western blot:* Asynchronous asexual blood stages of NF54 or NCR1-GFP were isolated from their host red blood cell with a lysis in 0.1% saponin in PBS (supplemented with protease inhibitor cocktail, Roche) 10 min on ice. Stage V gametocytes (NF54 or NCR1-GFP) were similarly isolated, but with two sequential lyses in 0.03% saponin. Following washes of the parasite pellets in PBS to remove haemoglobin, parasites were lysed in RIPA (150 mM NaCl, 5 mM EDTA pH 8, 50 mM Tris pH 7.5, 1% NP40, 0.5% sodium deoxycholate, 0.1% SDS, protease inhibitor cocktail) 30 min on ice, followed by water bath sonication (3 x 30 sec, Diagenode Bioruptor Pico). Lysates were centrifuged 30 min at 21'300 g at 4°C. Supernatants were collected, LDS sample buffer (NuPAGE) and 50 mM DTT were added. Samples were loaded on a NuPAGE 4-12% pre-cast gel without being boiled (for better detection of transmembrane proteins). Electrophoresis was done in Tris-Glycine running buffer, samples were transferred to a nitrocellulose membrane (Cytivia). Primary anti-GFP antibody was incubated overnight at 4°C (Amsbio TP401; 0.1 mg/ml in 5% milk-PBST) while anti-actin (hybridoma provided by the Soldati lab; 1:10 in 5% milk-PBST) was incubated 1h at room temperature. Secondary antibodies were probed 1h at room temperature (anti-rabbit-HRP and anti-mouse-HRP respectively, Invitrogen). Blots were imaged on a qTouch imager (RWD Life Science) in presence of Immobilon ECL ultra (Millipore).

**Asexual growth inhibition assay**

*Compounds:* Compounds tested were MMV009108, (Enamine, Compound 1), MMV028038 (Enamine, Compound 2), MMV019662 (Enamine, Compound 3), AR003UJV (Aaron Chemicals), AR00CYV7 (Aaron Chemicals), AR002OQ4 (Aaron Chemicals), AR00IMC2 (Aaron Chemicals), F2573-0310 (Life Chemicals Inc.), F2573-0329 (Life Chemicals Inc.), STL441565 (Vitas M Chemical Limited), STL055711 (Vitas M Chemical Limited), V029-3356 (ChemDiv Inc.), G856-4269 (ChemDiv Inc.), G856-4236 (ChemDiv Inc.), and 5102735 (ChemBridge Corporation).

Ring-stage parasites were exposed for 72 h to a serial dilution of compounds. For the three MMV compounds, the two-fold serial dilution started at 25 µM; for the twelve compounds identified *in silico*, a first assay was performed starting at 25 µM. The six compounds (F2573-0310, F2573-0329, STL441565, V029-3356, G856-4269 and G856-4236) that showed growth inhibition at the maximum concentrations were further tested starting at 60 µM to calculate their $EC_{50}$ accurately. Assays were carried out in a 384-well plate (culture of 50 µl, with a maximum of 0.6% DMSO), in technical duplicates and biological triplicates. Starting cultures were diluted to 0.2% ring-stage parasitemia in 2% hematocrit. After 72 h, following a freeze-thaw cycle, 10 µl of culture was incubated with 25 µl of Malstat solution (0.1 M Tris pH 9.0, 20 g/L lactic acid, pH 7.5, 0.2% Triton X-100, 0.5 g/L acetylpyridine adenine dinucleotide (APAD, Sigma), 200 µg/mL nitroblue tetrazolium (NBT; Sigma), 1 µg/mL phenozine ethosulfate (PES; Sigma)). The reaction reveals the presence of parasitic Lactate Dehydrogenase (LDH) – indicative of parasite growth - with a change in colour, measured by absorption at 650 nm on a SpectraMax Paradigm plate reader. Technical replicate averages were then normalized with the DMSO control and uninfected red blood cell (uRBC) culture controls using the formula below. Values thereby obtained were plotted against the logarithm of compound concentrations, together with a non-linear regression was calculated using GraphPad Prism to obtain the half inhibitory concentrations ($EC_{50}$).

$$\left(1 - \frac{|(test)| - |(uRBC)|}{|(DMSO)| - |(uRBC)|}\right) * 100$$

**Molecular dynamics**

*Membrane assembly*: Following the established procedures [9, 10], the experimentally solved cryo-EM structure of PfNCR1 bound to cholesterol hemisuccinate (CHS) was uploaded to the Position of Proteins in Membranes (PPM) server [11] to determine its position in the membrane. The experimentally not resolved loops and domains were also not included in the model. The martinize2 pipeline [12] was used to convert the PfNCR1 into a coarse-grained representation employing the Martini 3 forcefield [13]. The insane.py script [14] was used to embed PfNCR1 in a lipid bilayer with the following ratio: POPC:POPE:CHOL = 10:15:25. The system was solvated in water and salt was added to a concentration of 150 mM. Molecular dynamics (MD) simulations were carried out using GROMACS version 2024.1 [15-17]. This procedure was performed 5 times to independently assemble 5 systems of membrane-embedded PfNCR1.

To equilibrate the membranes, a 1 µs MD simulation was conducted in the presence of position restraints on the protein, with a force constant of 1000 kJ $mol^{-1}$ $nm^{-2}$ at a temperature of 310 K maintained using the v-rescaling thermostat [18]. A pressure of 1 bar was maintained using the c-rescaling barostat [19]. The Reaction-Field algorithm was used for electrostatic interactions with a cut-off of 1.1 nm, and a single cutoff of 1.1 nm was used for Van der Waals interactions. A time step of 0.02 ps was used. The equilibrated systems were converted to an all-atom representation (backmapping) using a version of the backward.py script [20], which was modified to work on Martini 3 structures.

*Preparation of ligand-bound PfNCR1 systems*: to avoid infrequent structural impressions induced by the coarse-grained to all-atom conversion, the protein was replaced with the original cryo-EM PfNCR1 complexes. Possible atom overlaps between the inserted protein and the equilibrated membrane were relaxed using the membed procedure [21] as previously described [22]. Protonation states were defined according to the hydrogen bonding pattern of the cryo-EM conformation. Five repeats for every system were assembled.

*Preparation of ligand-free PfNCR1 systems:* The closed and open SSD conformations of PfNCR1 were extracted from the cryo-EM structures of PfNCR1 bound to CHS and Compound 3, respectively. The pKa of the ionizable groups was estimated using PROPKA [23]. D571 and D1383 were protonated in all apo simulations, while the protonation state of D570 and H1387 was modified according to the conformation dependent pKa predictions. In the open SSD conformation D570 was predicted to be protonated and H1387 neutral, while in the closed SSD conformation, D570 was predicted to be negatively charged and H1387 double protonated and therefore positively charged. In a second set of simulations, to probe the impact of protonation, the protonation state of D570 and H1387 was defined as predicted for the alternative conformations. The differently protonated PfNCR1 structures were prepared and parametrized using GROMACS then embedded in the assembled membranes using the membed procedure. Five repeats for every system were assembled.

*All-atom simulation*: We used the amber99sb-ildn force field [24] to describe the protein, ions, and solvent, and Slipid [25, 26] for the lipid membrane. The topology parameters for the ligands were generated using the AnteChamber PYthon Parser interfacE (ACPYPE) server [27, 28] according to the Generalised Amber Force Field 2 (GAFF2) [29].

The assembled systems were energy-minimized and equilibrated in four steps of 2.5 ns each, with gradually releasing the position restraints (1000, 100, 10, 1 kJ $mol^{-1}$ $nm^{-2}$) applied on the Cα atoms and the atoms of the ligands [9, 30]. Then a production run (with no position restraints) of 300 ns was carried out for the ligand-bound PfNCR1 systems, while a 500 ns production simulation was carried out for the ligand-free PfNCR1 systems. The temperature was maintained at 310 K using a v-rescale ($\tau$ = 0.5 ps) thermostat [18], while separately coupling protein, membrane, and solvent. The pressure was maintained at 1 bar using the Parrinello-Rahman barostat [31] in a semi-isotropic manner and applying a coupling constant of 20.1 ps. Long-range electrostatic interactions were described using the smooth particle mesh Ewald method [32], applying a cutoff of 0.9 nm. The van der Waals interactions were described using the Lennard-Jones potentials, applying a cutoff of 0.9 nm. Long-range corrections for energy and pressure were applied.

*Analysis*: The trajectories were processed and analysed using the GROMACS package. To generate the ligand densities from the MD trajectories the GROMACS spatial tool was used. For the principal component analysis (PCA): all the trajectories from all the ligands in different starting poses were concatenated, and the covariance matrix was calculated from this concatenated trajectory using GROMACS covar tool. Then the trajectories corresponding to each system were projected on principal component 1 and 2 using GROMACS anaeig tool. To calculate RMSDs and distances of the structures in the MD simulations we used python scripts employing the MD Analysis package, v2.9.0 [33, 34]. For visualization, we used VMD [35] v1.9.4 and Pymol version 2.5 (**Figures S12-15**).

**In silico screening**

*Ligand-based virtual screening:* MMV019662 (Compound 3) was used as the query ligand to search for structurally similar compounds in the MolPort website (https://www.molport.com/shop/find-chemicals), and the top-ranked compounds by similarity were visually inspected to select the promising analogs to purchase.

*Structure-based virtual screening:* All compounds in the MolPort all-stock library with a molecular weight below 600 Da were prepared with Schrödinger's LigPrep [36] (Schrödinger version 2022-4) with the following settings: the OPLS4 force field [37] for geometry optimization and energy minimization, protonation state generation at pH 0.0 using Epik [38], and stereoisomer generation was limited to a maximum of four per ligand. These settings ensured a comprehensive yet computationally efficient representation of ligand conformational and protonation variability (**Figure S16)**.

The protein structures were first prepared with Schrödinger’s Protein Preparation Wizard [39, 40] (Schrödinger, LLC). The process included assigning bond orders, adding hydrogen atoms, and optimizing the hydrogen-bonding network. Protonation states of ionizable residues were predicted at physiological pH (~7.4) using the Epik module. The structure was then energy-minimized using the OPLS4 force field. The prepared structures were then loaded into Hemes to reassign hydrogen atoms and define the binding pocket with the selected ligands.

All prepared compounds from the MolPort library were docked into each binding pocket using the GOLD docking algorithm [41] (version 5.8.1; Cambridge Crystallographic Data Centre, Cambridge, UK) with the GoldScore scoring function. To accelerate the calculations, the search efficiency was set to 0.3, and each ligand was docked into each binding site three times. The top-ranked compounds for each pocket, based on docking scores, were subsequently filtered to retain only those with fewer than 10 rotatable bonds and a calculated logP value below 6.5.

Following filtration, the remaining top-ranked poses were visually inspected for key interactions, such as hydrogen bonding and hydrophobic contacts.

#### Data availability

The MD simulation trajectories and data have been deposited to Zenodo (10.5281/zenodo.22214685). All coordinates and cryo-EM density maps have been deposited to the Protein Data Bank (9THA, https://doi.org/10.2210/pdb9THA/pdb; 9THB, https://doi.org/10.2210/pdb9THB/pdb; 9THC, https://doi.org/10.2210/pdb9THC/pdb; 9THD, https://doi.org/10.2210/pdb9THD/pdb; 9THE, https://doi.org/10.2210/pdb9THE/pdb;) and Electron Microscopy Data Bank (EMD-55920, https://www.ebi.ac.uk/pdbe/entry/emdb/EMD-55920; EMD-55921, https://www.ebi.ac.uk/pdbe/entry/emdb/EMD-55921; EMD-55922, https://www.ebi.ac.uk/pdbe/entry/emdb/EMD-55922; EMD-55923, https://www.ebi.ac.uk/pdbe/entry/emdb/EMD-55923; EMD-55924, https://www.ebi.ac.uk/pdbe/entry/emdb/EMD-55924;). All other data generated in this study are provided in the Supplementary Information/Source Data file.

### Supplementary Figures

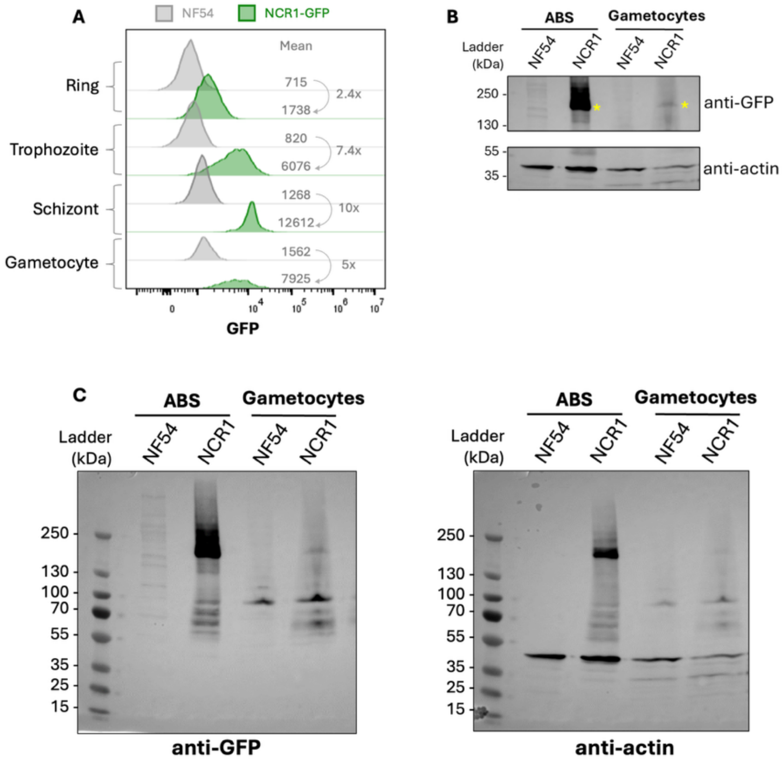


**Figure S1:** NCR1 is expressed in both the asexual blood stages (ABS) and in gametocytes. Parasites expressing NCR1-GFP [8] were analysed by flow cytometry and western blot to detect expression of NCR1 at different stages. **A.** GFP signal was detected in all stages of NCR1-GFP parasites (green) compared to the NF54 control (grey). In particular, GFP signal increased throughout the ABS (rings < trophozoites < schizonts). GFP signal was also detected in NCR1-GFP gametocytes (one representative histogram out of three biological replicates, shown in Figure S2). Mean GFP fluorescence is indicated for each condition, as well as fluorescence fold-change of NCR1-GFP compared to the corresponding NF54 control. **B.** Parasite extracts of asynchronised ABS or stage V gametocytes of either NF54 or NCR1-GFP parasites were analysed by western blot. GFP antibody detected a band corresponding to NCR1-GFP (~ 197 kDa; yellow star) in both ABS and gametocytes (although fainter) of NCR1-GFP parasites. Loading control: anti-actin (~ 42 kDa). **C.** Full western blots corresponding to B.

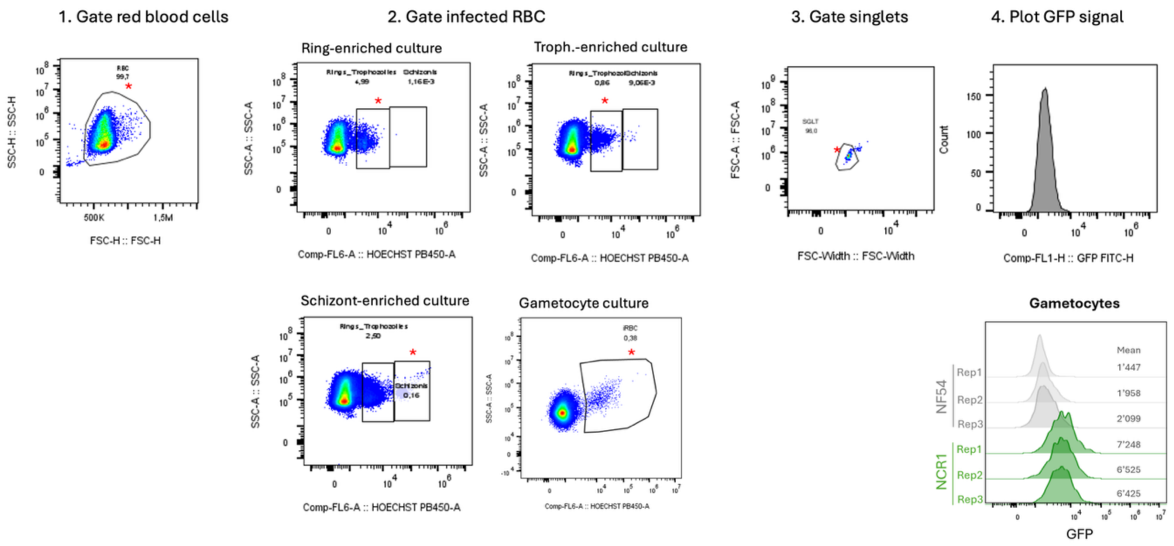


**Figure S2:** Flow cytometry gating strategy. Following staining with Hoechst 33342, cultures of asexual blood stages or gametocytes (either NF54 or NCR1-GFP parasites) were analysed with a CytoFLEX S. At least 5'000 infected red blood cells were acquired. First, red blood cells were gated (FSC-H / SSC-H), then infected cells were gated based on Hoechst content (same gate for rings and trophozoites, separate gate for schizonts, and for gametocytes). Singlets were gated on FSC-Width / FSC-A. Finally, GFP signal was plotted and mean value indicated. Here the three biological replicates of gametocyte cultures are shown. Red star indicates the population selected for the next step. One example of NCR1-GFP parasites is shown (4 for the gating of infected RBCs). The same gatings were used for NF54 parasites. Acquisition gains were: FSC: 120, SSC: 100; FITC: 180; PB450: 68.

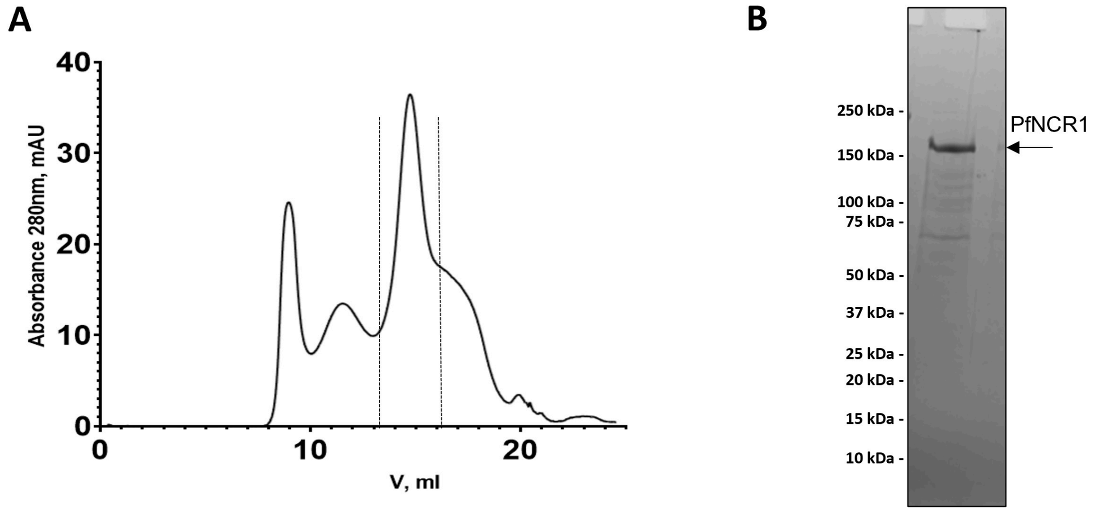


**Figure S3:** Expression and purification of PfNCR1**. A.** Size-exclusion chromatography (SEC) profile following GFP-NB affinity purification. **B.** SDS-PAGE analysis with Coomassie staining.

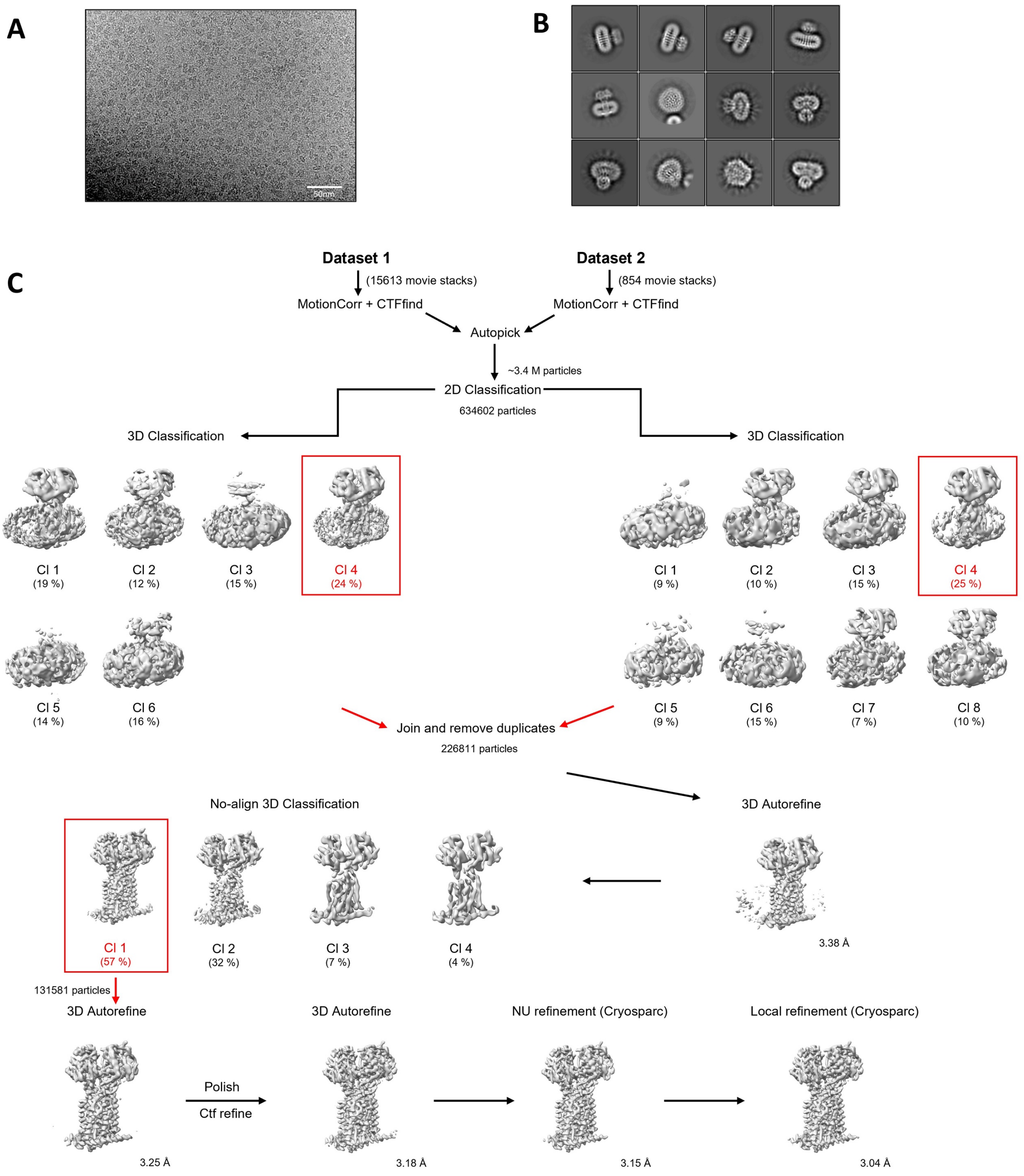


**Figure S4:** Cryo-EM image processing pipeline of apo-PfNCR1 **A.** A representative cryo-EM micrograph of apo-PfNCR1 dataset (scale bar 50 nm). **B.** Selected classes after 2D classification. The box size is 400 Å. **C.** Overview of the cryo-EM apo-PfNCR1 processing pipeline. Selected 3D classes for further processing are indicated with red boxes.

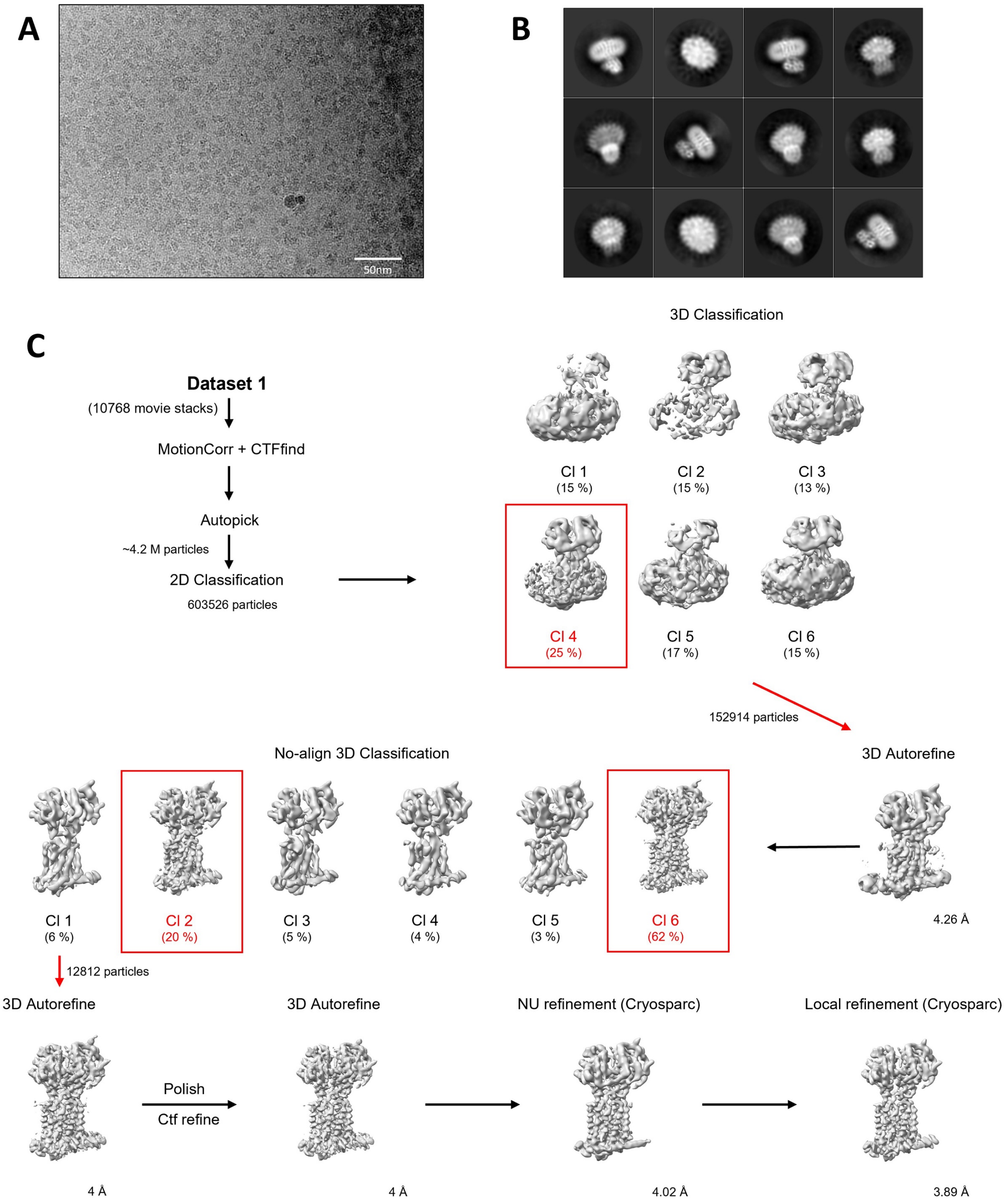


**Figure S5:** Cryo-EM image processing pipeline of MMV009108 (Compound 1)-PfNCR1 **A.** A representative cryo-EM micrograph of Compound 1-PfNCR1 dataset (scale bar 50 nm). **B.** Selected classes after 2D classification. The box size is 400 Å. **C.** Overview of the cryo-EM Compound 1-PfNCR1 processing pipeline. Selected 3D classes for further processing are indicated with red boxes.

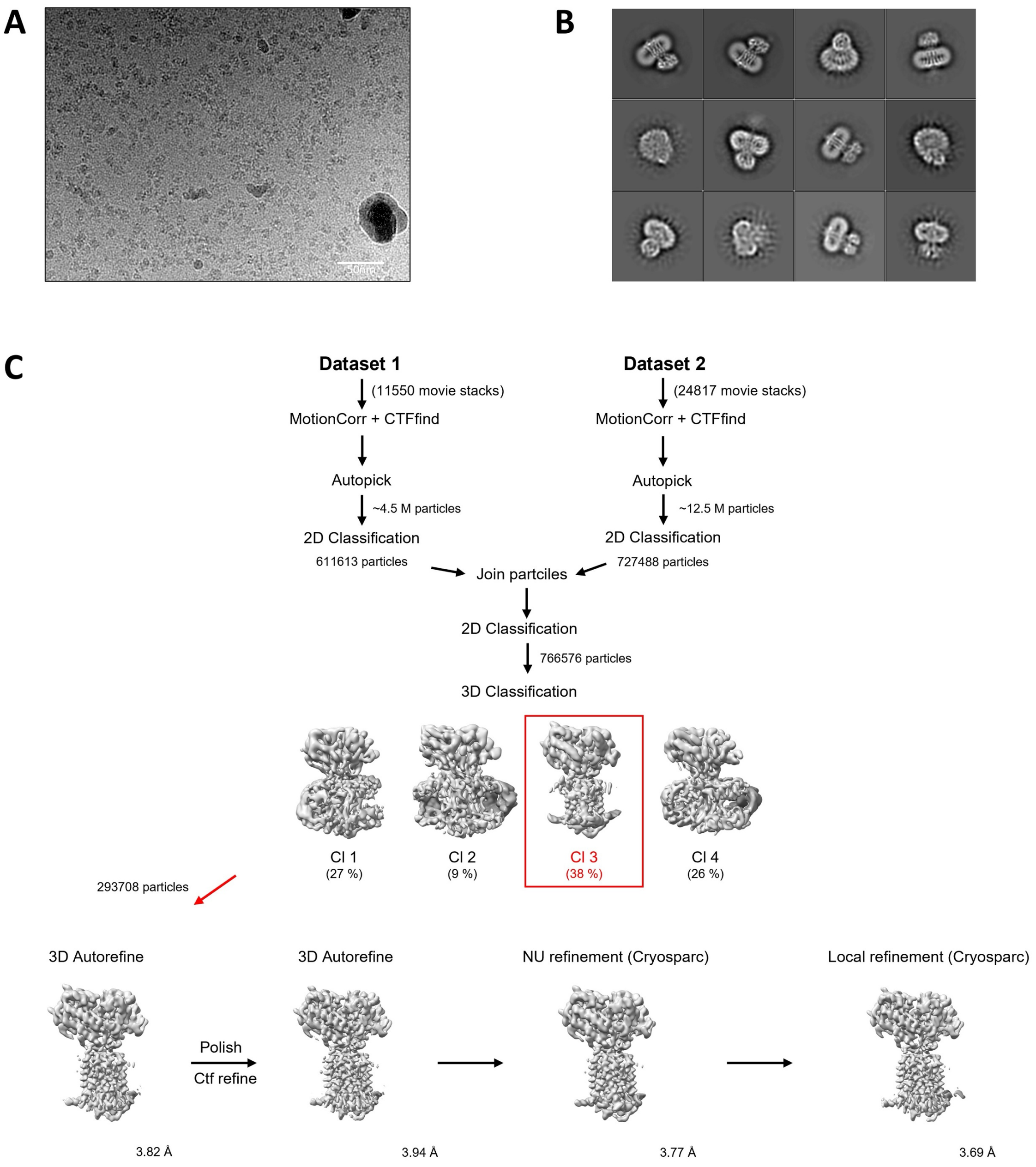


**Figure S6:** Cryo-EM image processing pipeline of MMV028038 (Compound 2)-PfNCR1 **A.** A representative cryo-EM micrograph of Compound 2-PfNCR1 dataset (scale bar 50 nm). **B.** Selected classes after 2D classification. The box size is 400 Å. **C.** Overview of the cryo-EM Compound 2-PfNCR1 processing pipeline. Selected 3D classes for further processing are indicated with red boxes.

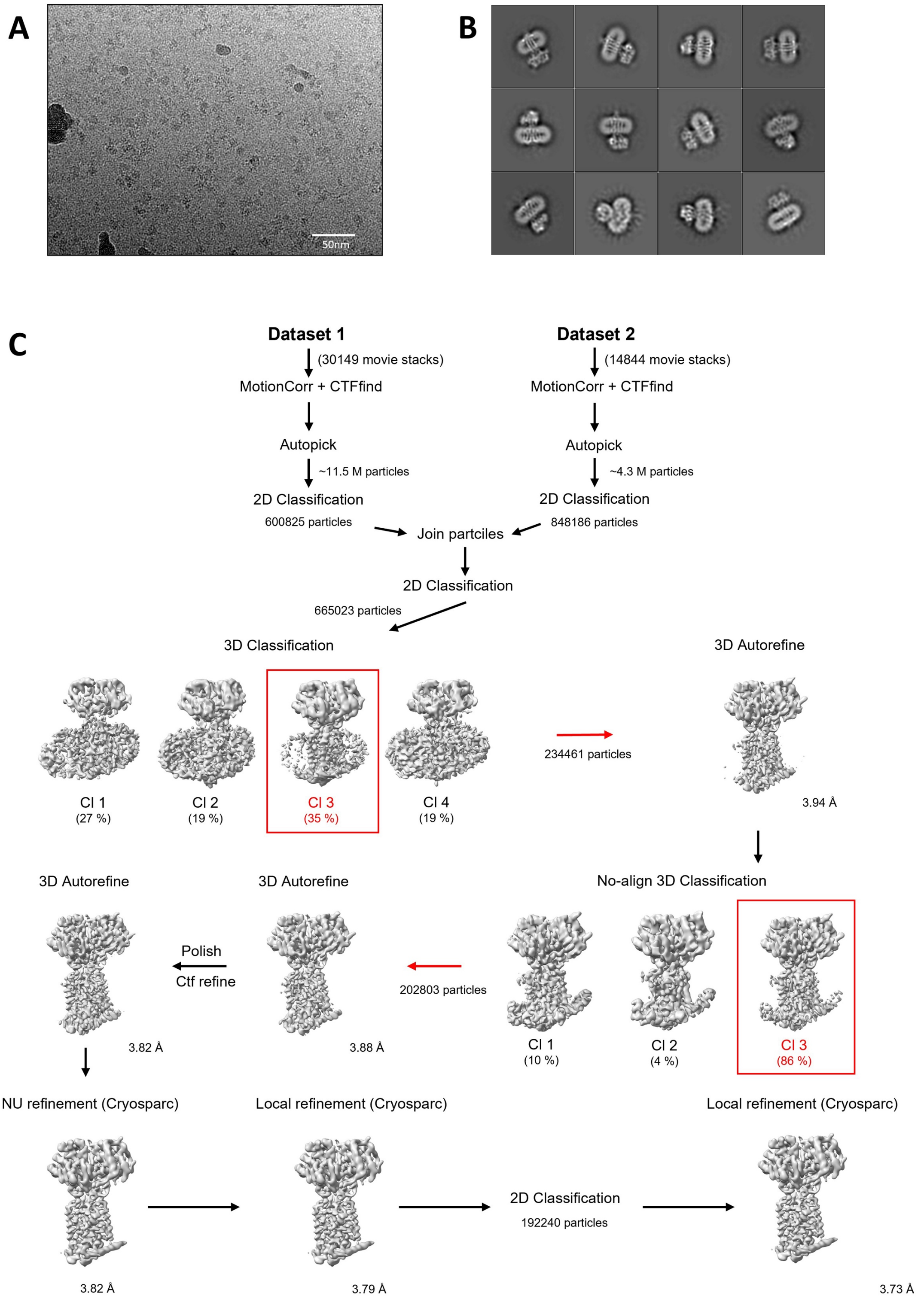


**Figure S7:** Cryo-EM image processing pipeline of MMV019662 (Compound 3)-PfNCR1 **A.** A representative cryo-EM micrograph of Compound 3-PfNCR1 dataset (scale bar 50 nm). **B.** Selected classes after 2D classification. The box size is 400 Å. **C.** Overview of the cryo-EM Compound 3-PfNCR1 processing pipeline. Selected 3D classes for further processing are indicated with red boxes.

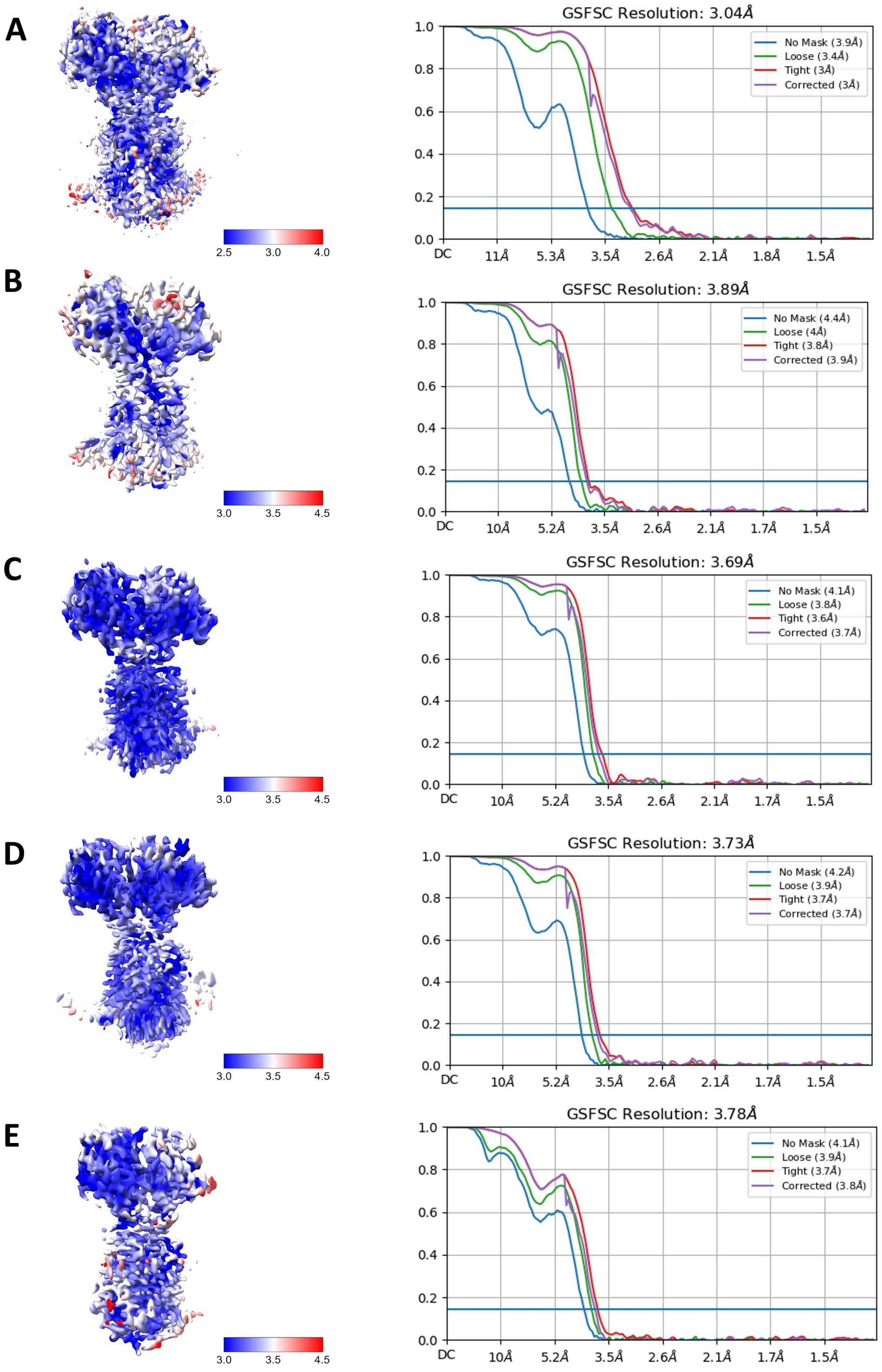


**Figure S8:** Cryo-EM data analysis of PfNCR1 **A.** Local resolution and FSC curve for apo-PfNCR1. **B.** Local resolution and FSC curve for MMV009108-PfNCR1 (Compound 1). **C.** Local resolution and FSC curve for MMV028038-PfNCR1 (Compound 2). **D.** Local resolution and FSC curve for MMV019662-PfNCR1 (Compound 3). **E.** Local resolution and FSC curve for G856-4236-PfNCR1 (Compound 4).

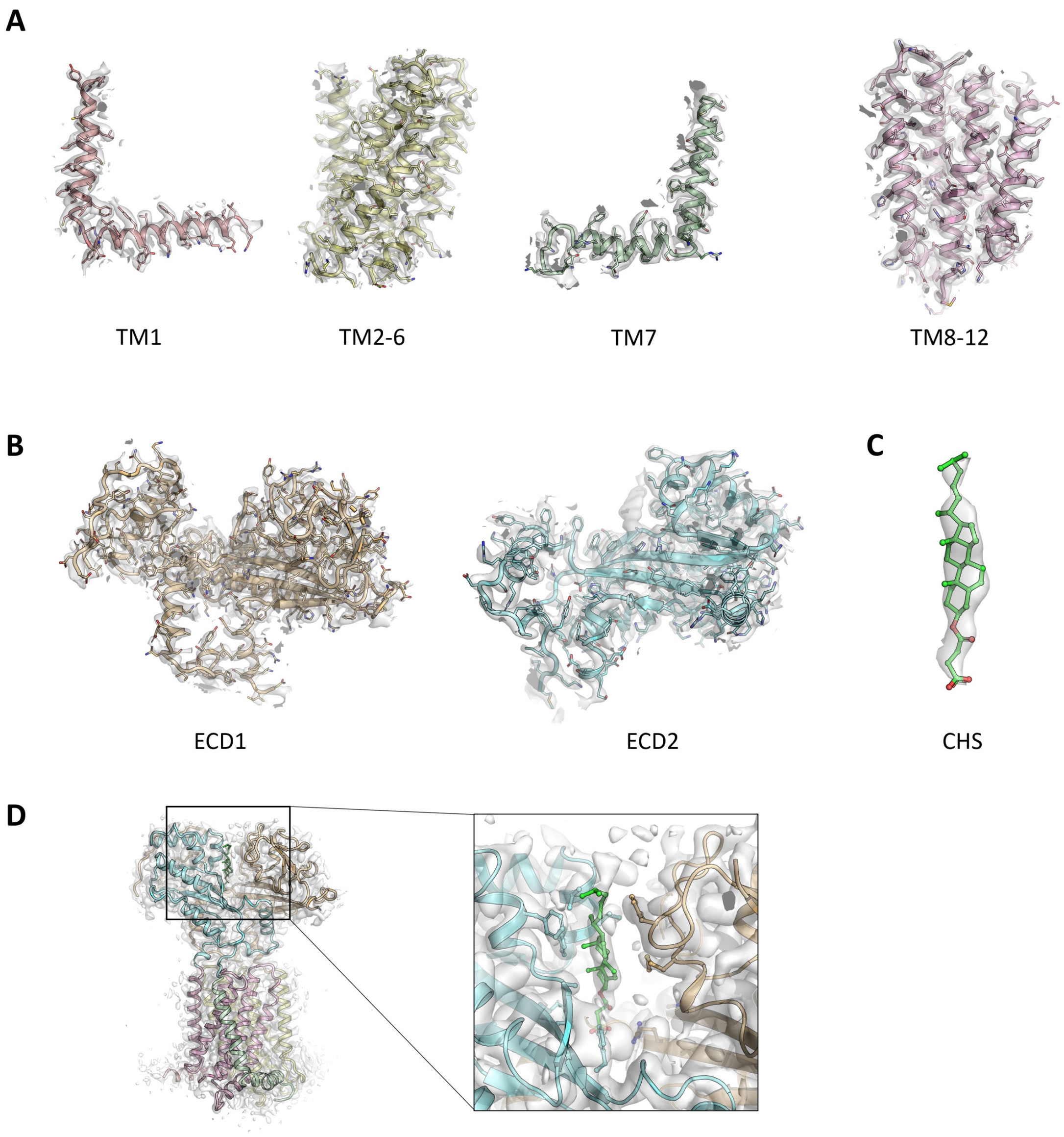


**Figure S9:** Quality of the PfNCR1-apo cryoEM density **A.** Cryo-EM density features of the TM domains. **B.** Cryo-EM density features of the ECD domains. **C.** Cryo-EM density features of CHS. **D.** Cryo-EM density features of the CHS binding site.

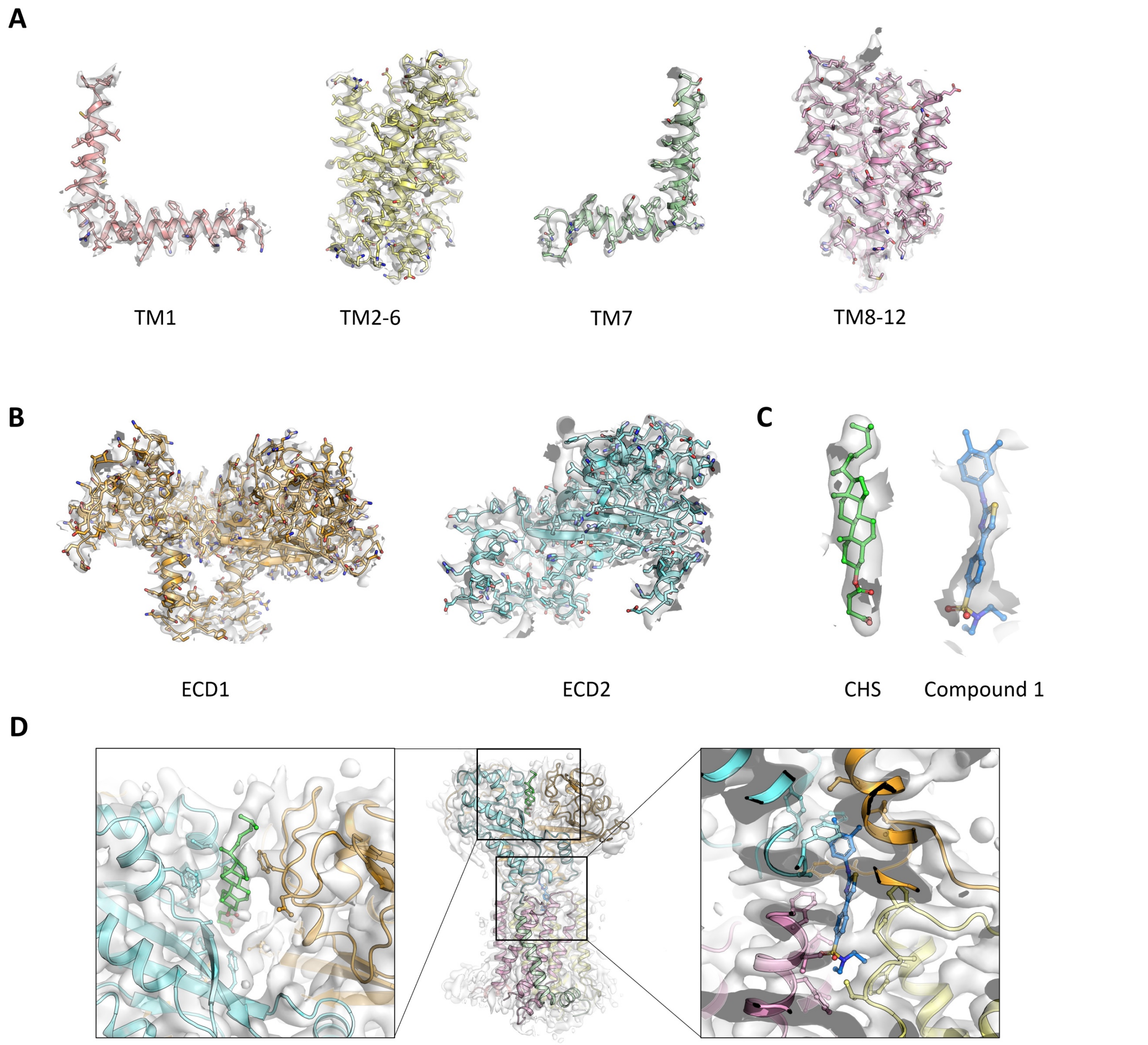


**Figure S10:** Quality of the PfNCR1-MMV009108 (Compound 1) cryoEM density **A.** Cryo-EM density features of the TM domains. **B.** Cryo-EM density features of the ECD domains. **C.** Cryo-EM density features of CHS in the ecto site and Compound 1 in the neck site. **D.** Cryo-EM density features of the CHS binding site and of the Compound 1 binding site.

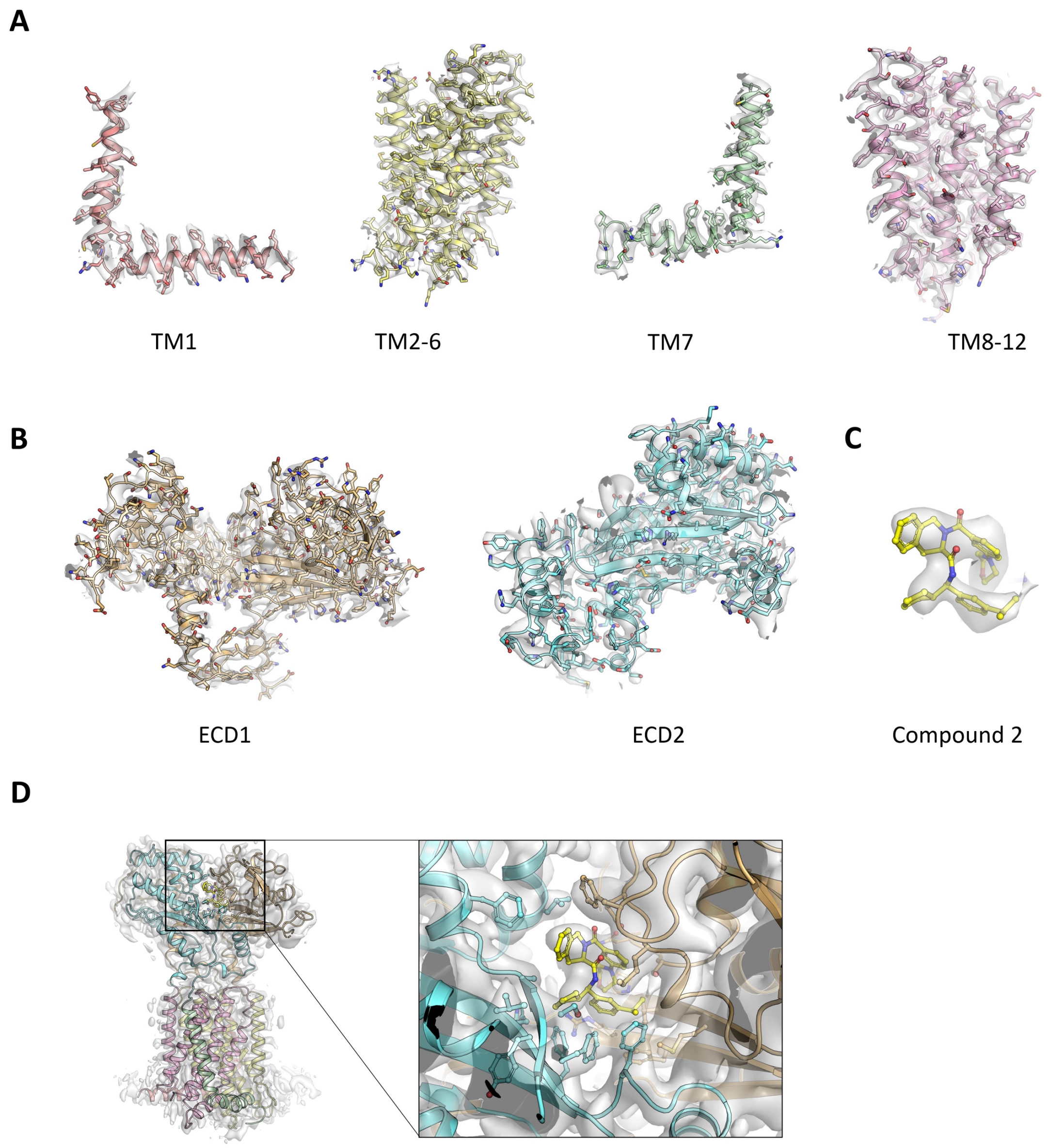


**Figure S11:** Quality of the PfNCR1-MMV028038 (Compound 2) cryoEM density **A.** Cryo-EM density features of the TM domains. **B.** Cryo-EM density features of the ECD domains. **C.** Cryo-EM density features of Compound 2 in the ecto site. **D.** Cryo-EM density features of the Compound 2 binding site.

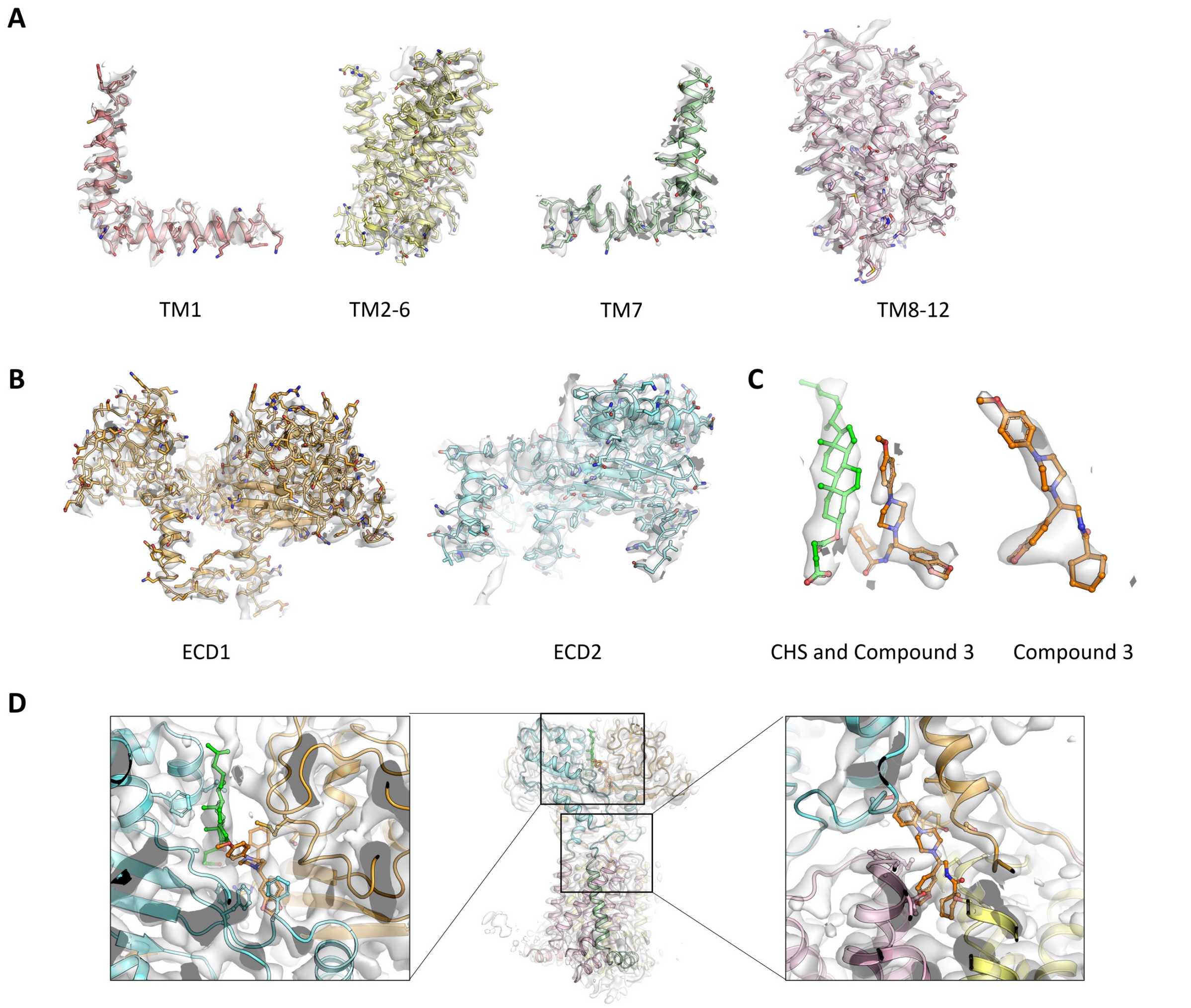


**Figure S12:** Quality of the PfNCR1-MMV019662 (Compound 3) cryoEM density **A.** Cryo-EM density features of the TM domains. **B.** Cryo-EM density features of the ECD domains. **C.** Cryo-EM density features of CHS and Compound 3 in the ecto site and Compound 3 in the neck site. **D.** Cryo-EM density features of the ecto binding site and of the neck binding site.

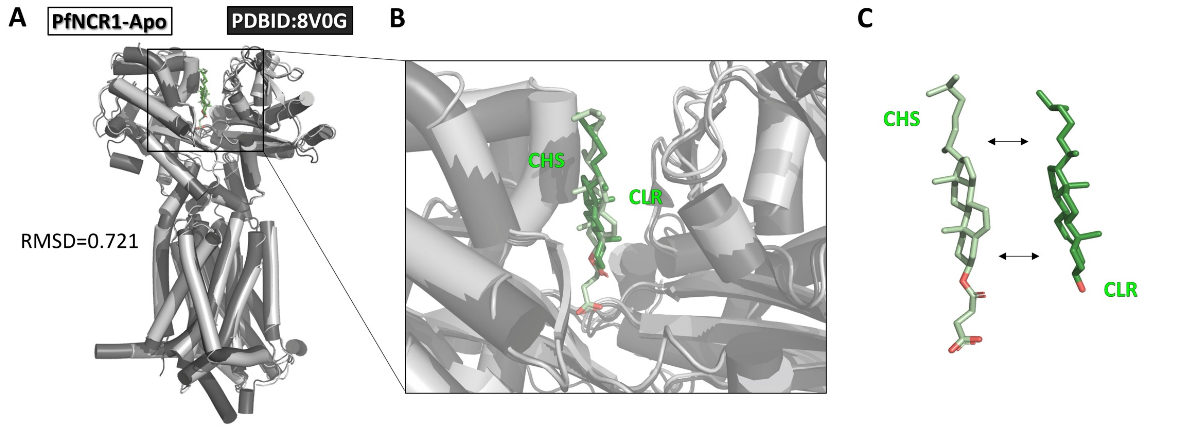


**Figure S13:** Comparison of PfNCR1-Apo and published apo 8V0G. **A.** Our PfNCR1-Apo in grey and 8V0G [42] in black show low RMSD overall. **B.** Close up look on the ecto site, in light green our modeled CHS and in dark green their modeled CLR. **C.** CHS and CLR are shown separated for clarity. CHS was modeled rotated by 180° around the z-axis relative to the previously published CLR orientation, based on our MD simulation results.

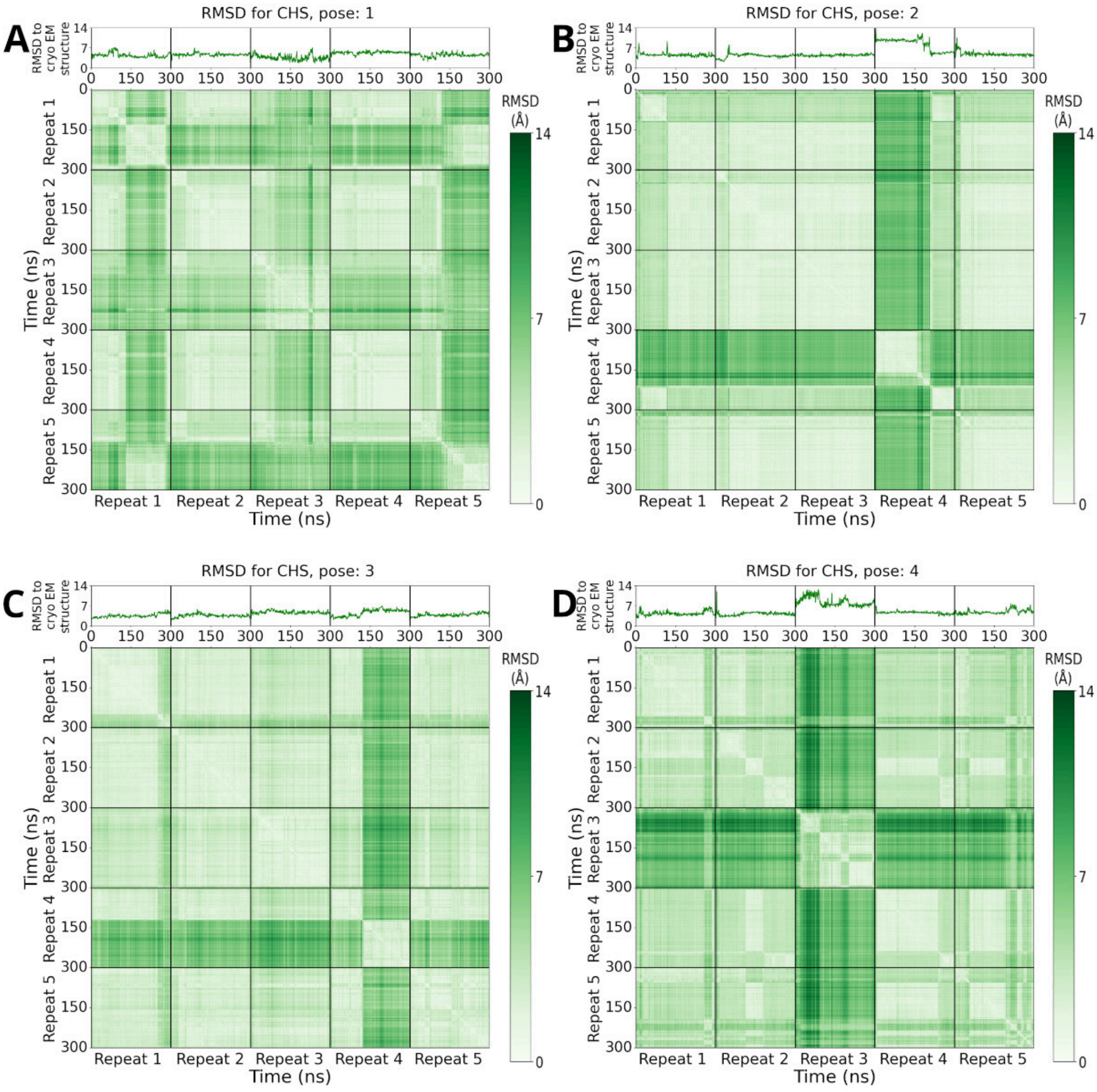


**Figure S14:** The RMSD for the poses suggested for CHS **A. B. C.** and **D.** show the RMSD changes during the simulations of CHS in poses 1, 2, 3 and 4, respectively. For each figure, we see on the top: The 1D RMSD of the ligand during the course of the five repeats of 300 ns simulations compared to the original starting pose suggested from the cryo-EM. On the bottom: we see the 2D RMSD of the ligand during the course of the five repeats of 300 ns simulations compared to itself.

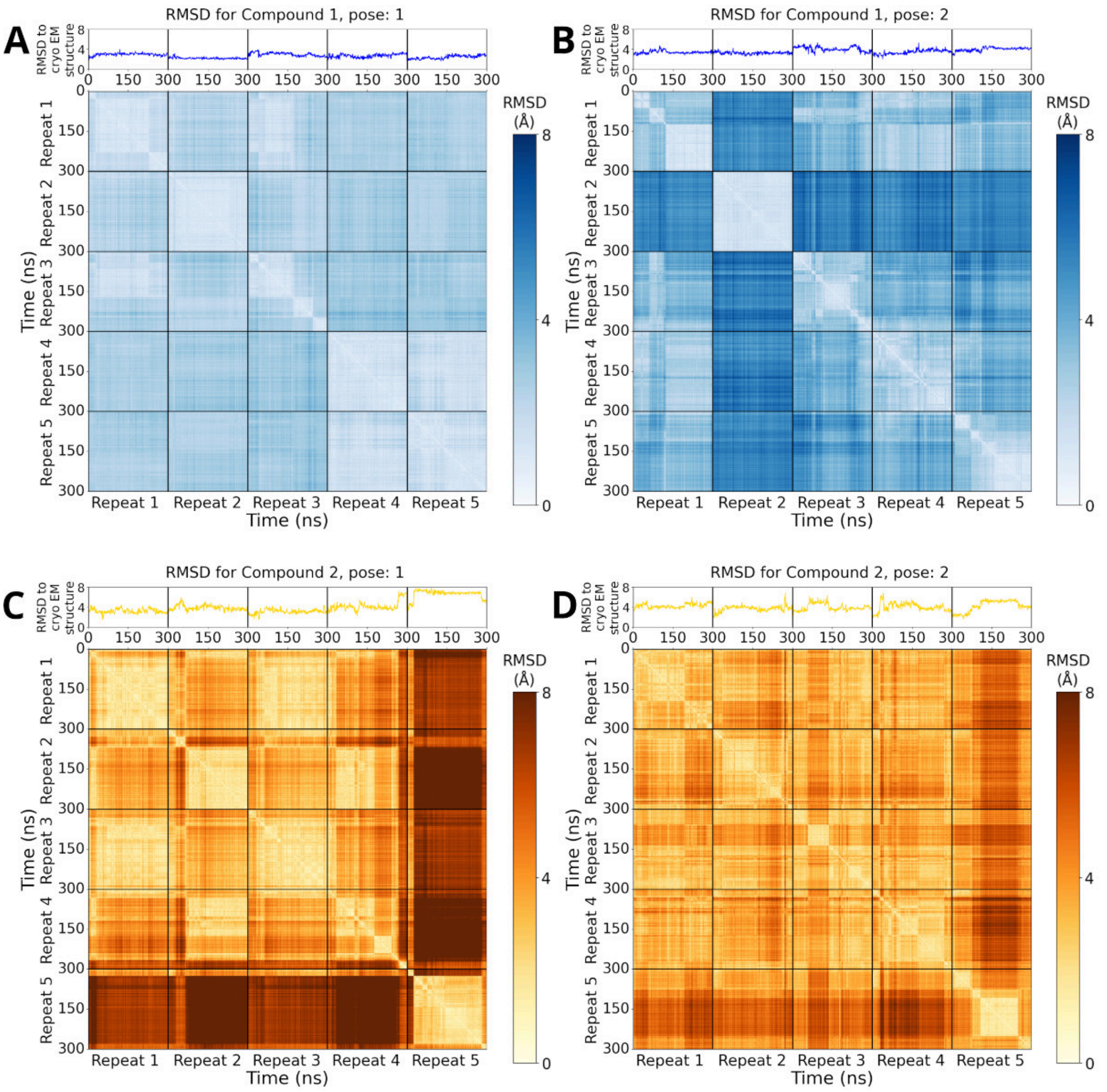


**Figure S15:** The RMSD for the poses suggested for MMV009108 (Compound 1) and MMV028038 (Compound 2) **A.** and **B.** show the RMSD changes during the simulations of Compound 1 in pose 1 and 2 respectively, while **C.** and **D.** show the RMSD changes during the simulations of Compound 2 in pose 1 and 2, respectively. For each figure, we see on top: The 1D RMSD of the ligand during the course of the five repeats of 300 ns simulations compared to the original starting pose suggested from the cryo-EM. On the bottom: we see the 2D RMSD of the ligand during the course of the five repeats of 300 ns simulations compared to itself.

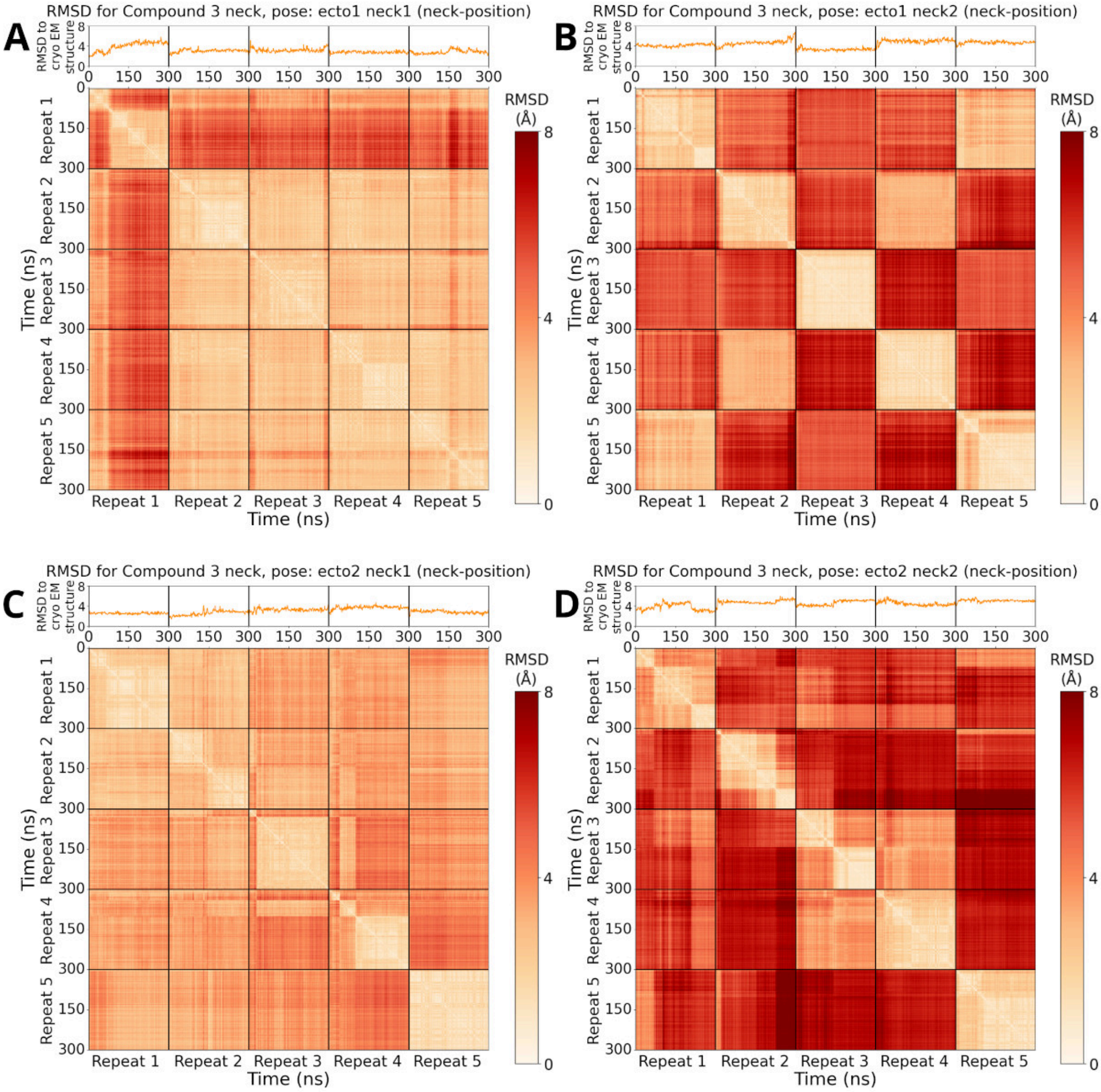


**Figure S16:** The RMSD for the poses suggested for MMV019662 (Compound 3) at the neck location **A. B. C.** and **D.** show the RMSD changes during the simulations of Compound 3 in poses ecto1_neck1, ecto1_neck2, ecto2_neck1 and ecto2_neck2, respectively. For each figure, we see on top: The 1D RMSD of the ligand during the course of the five repeats of 300 ns simulations compared to the original starting pose suggested from the cryo-EM. On the bottom: we see the 2D RMSD of the ligand during the course of the five repeats of 300 ns simulations compared to itself.

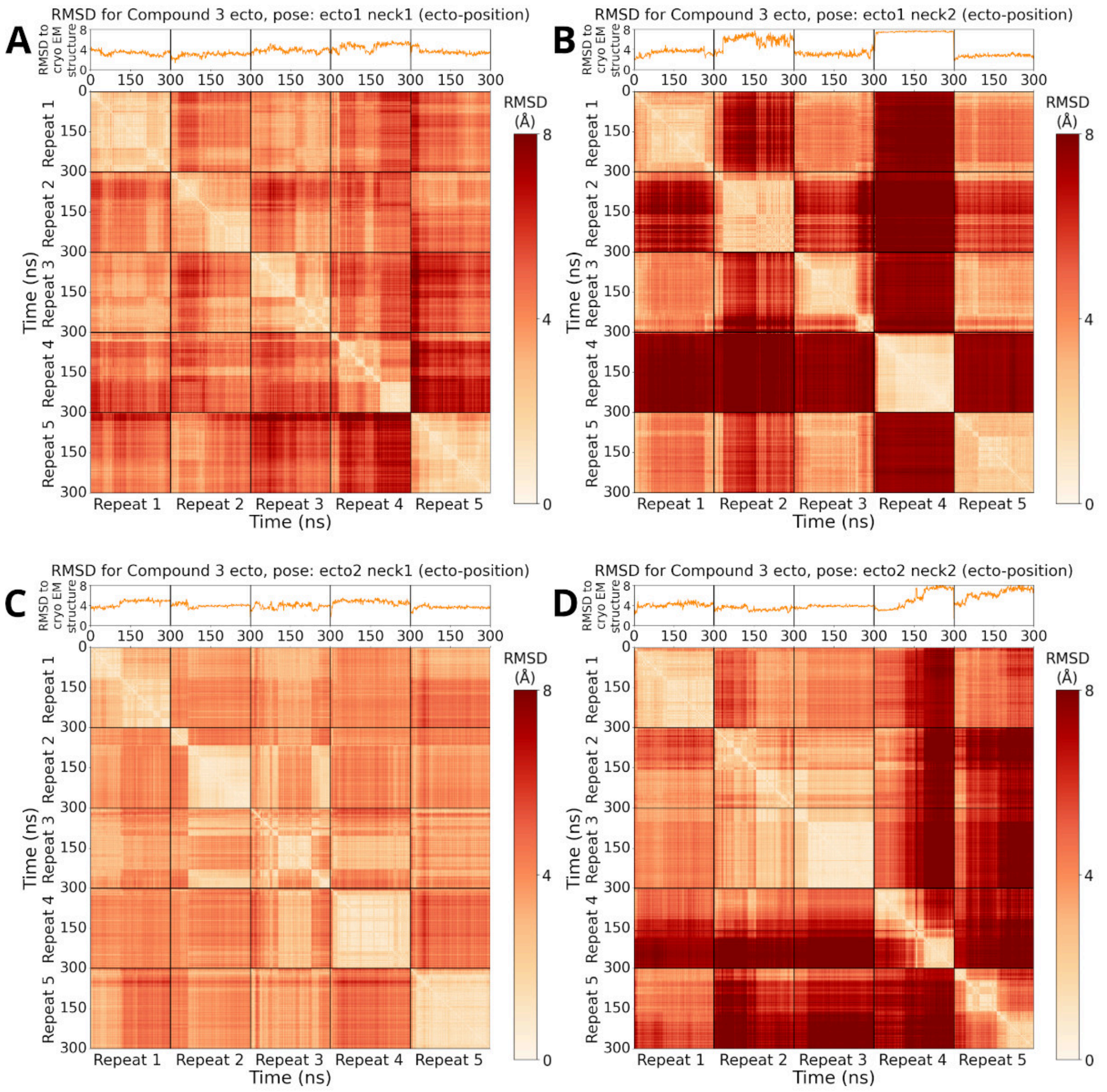


**Figure S17:** The RMSD for the poses suggested for MMV019662 (Compound 3) at the ecto location **A. B. C.** and **D.** show the RMSD changes during the simulations of Compound 3 in poses ecto1_neck1, ecto1_neck2, ecto2_neck1 and ecto2_neck2. respectively. For each figure, we see on top: The 1D RMSD of the ligand during the course of the five repeats of 300 ns simulations compared to the original starting pose suggested from the cryo-EM. On the bottom: we see the 2D RMSD of the ligand during the course of the five repeats of 300 ns simulations compared to itself.

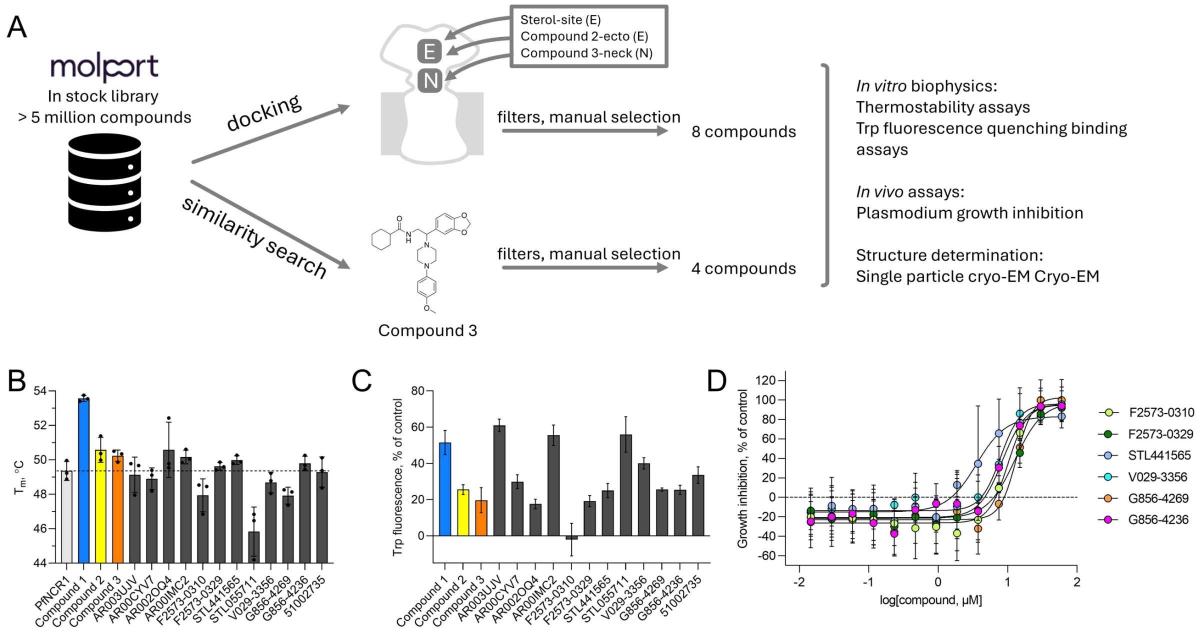


**Figure S18.** Search for new compounds targeting PfNCR1. **A.** Workflow for the computer-guided identification of the new PfNCR1 modulators. **B-C.** Biophysical characterization of compound-PfNCR1 interactions, based on thermostability (B) and Trp fluorescence quenching (C). **D.** *P. falciparum* growth inhibition assays reveal six compounds with biological activity (data for the remaining compounds that showed no effect within this concentration range – not shown). $EC_{50}$ = 10.11, 12.69, 3.56, 7.92, 13,40 and 8.42 μM respectively.

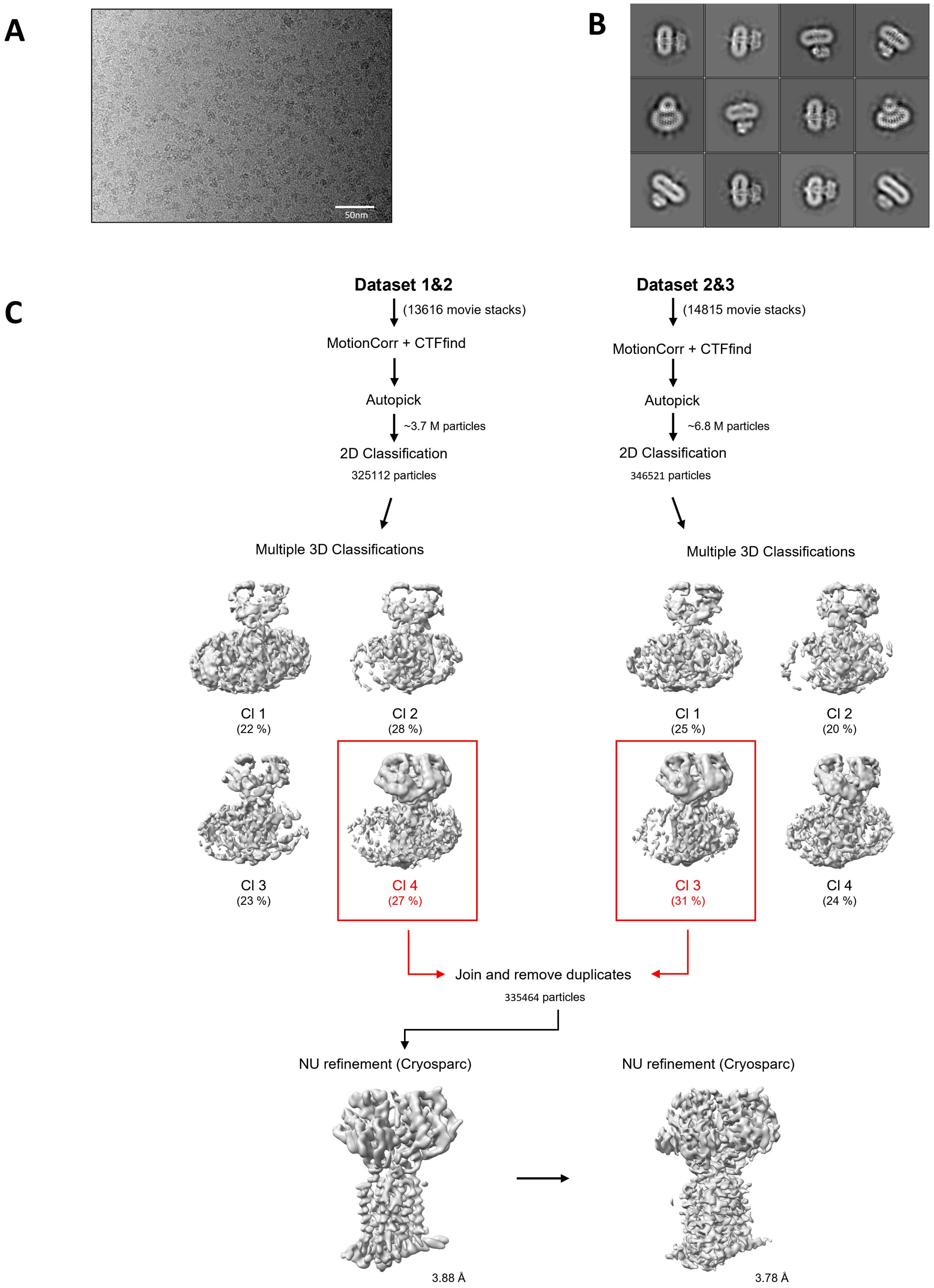


**Figure S19:** Cryo-EM image processing pipeline of G856-4236-PfNCR1 **A.** A representative cryo-EM micrograph of G856-4236-PfNCR1 dataset (scale bar 50 nm). **B.** Selected classes after 2D classification.

The box size is 400 Å. **C.** Overview of the cryo-EM G856-4236-PfNCR1 processing pipeline. Selected 3D classes for further processing are indicated with red boxes.

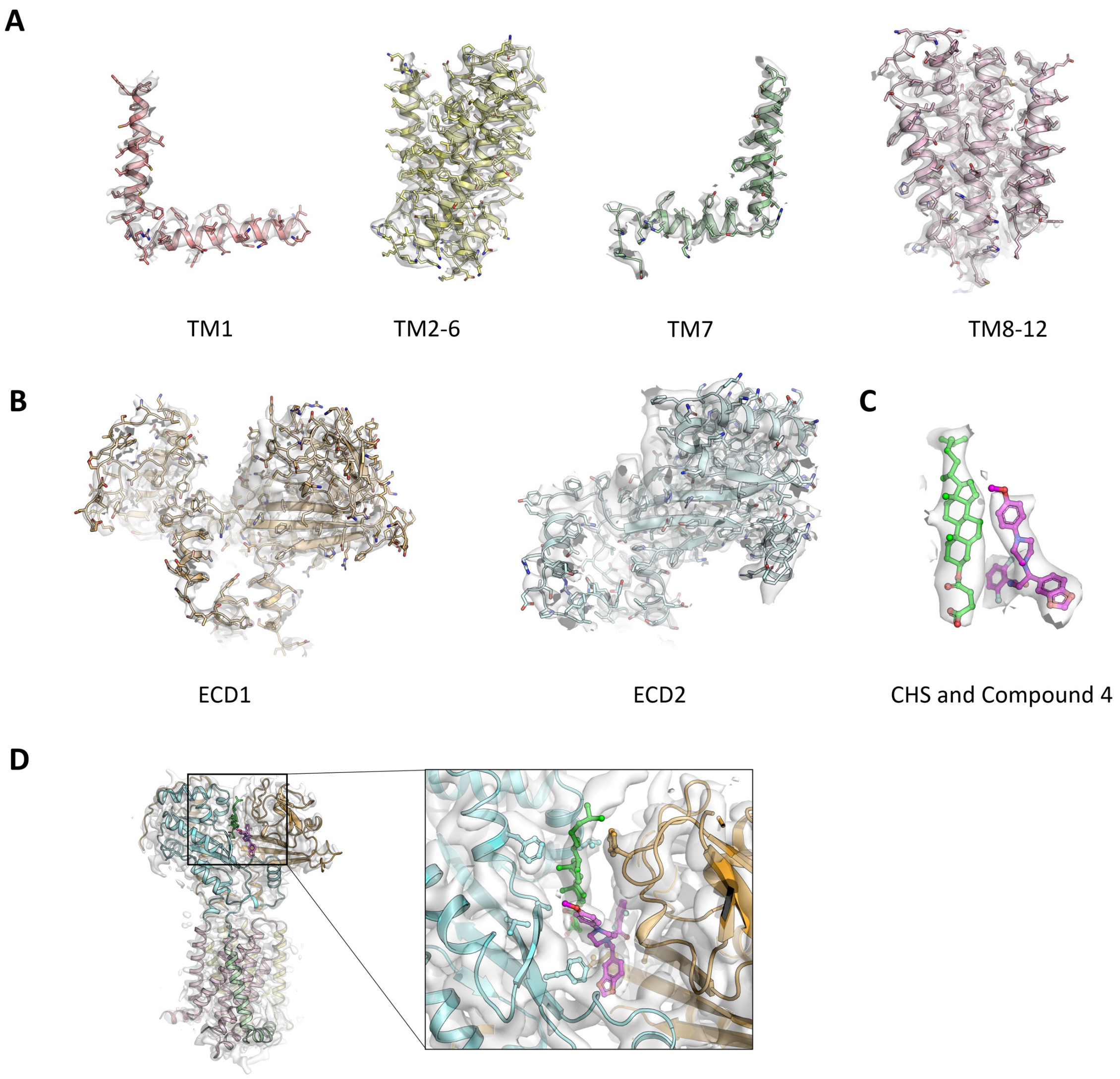


**Figure S20:** Quality of the PfNCR1-G856-4236 (Compound 4) cryoEM density **A.** Cryo-EM density features of the TM domains. **B.** Cryo-EM density features of the ECD domains. **C.** Cryo-EM density features of CHS and G856-4236 in the ecto site. **D.** Cryo-EM density features of the Compound 4 binding site.

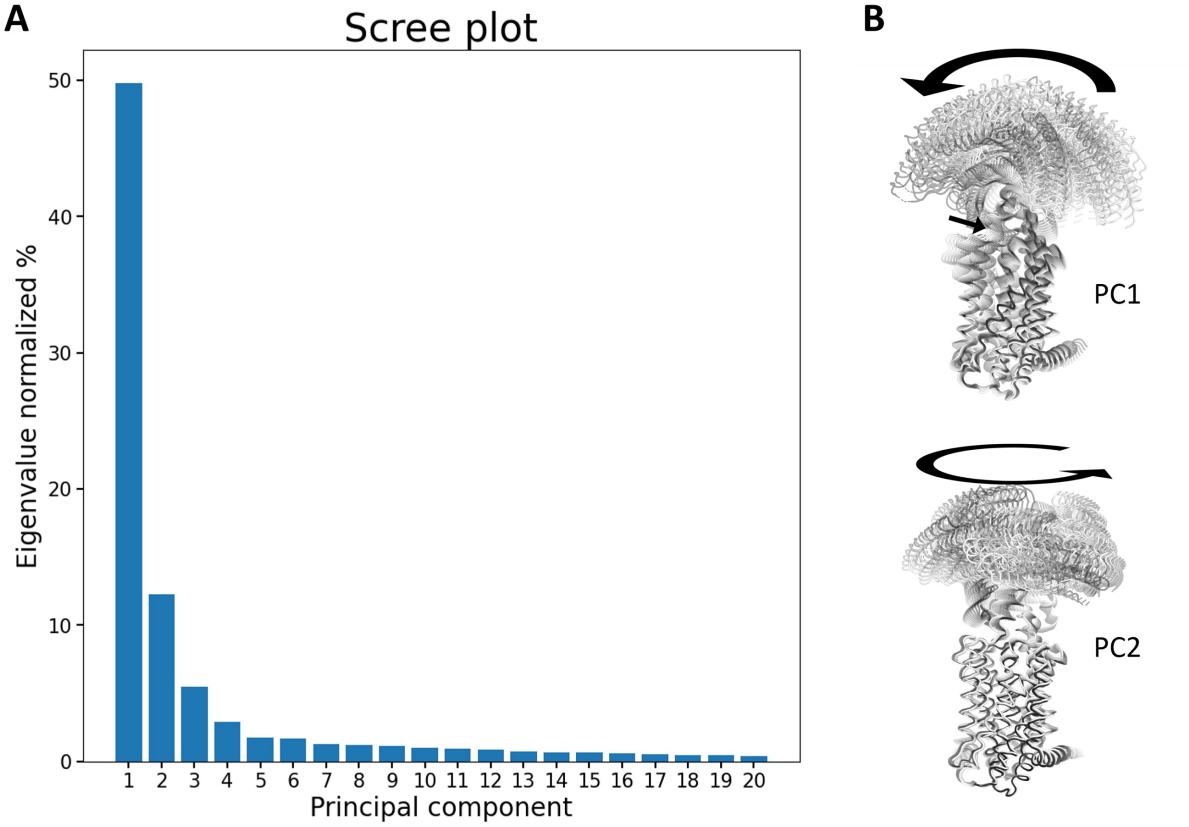


**Figure S21:** Principal component analysis of PfNCR1. **A.** The scree plot showing the eigenvalues of the principal components (normalized to 100). **B.** Bending and opening motion corresponding to PC1 and Twisting motion corresponding to PC2.

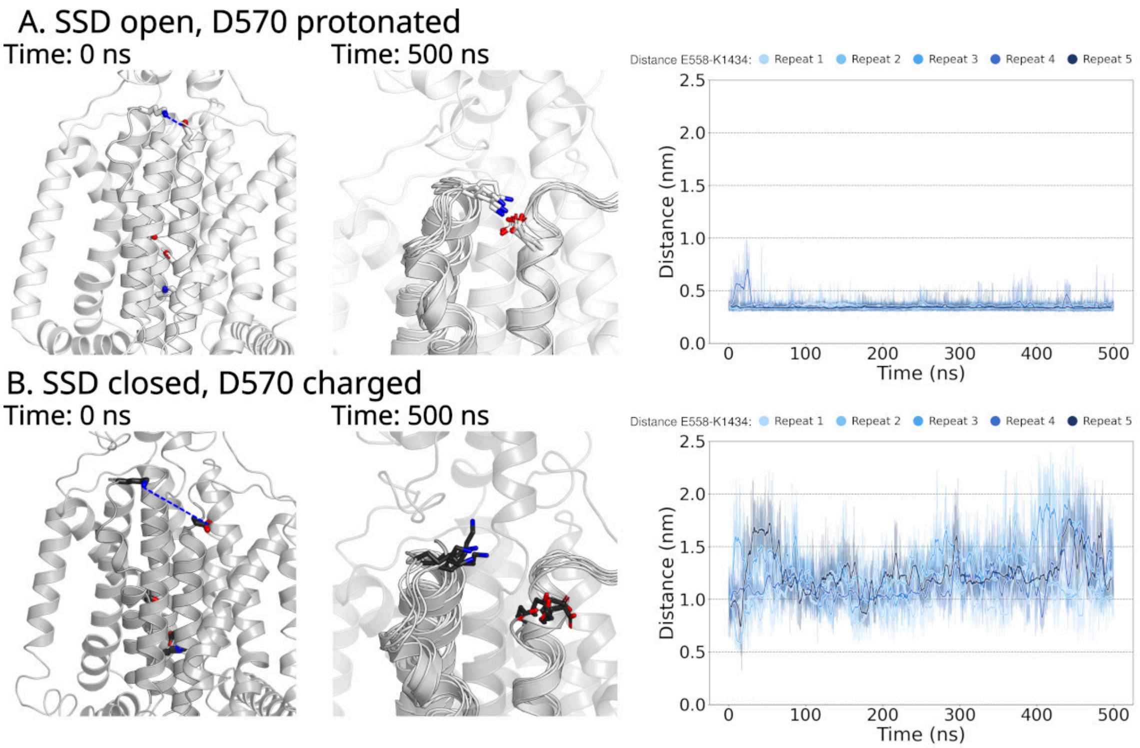


**Figure S22: A.** Simulations of apo PfNCR1 in the open SSD conformation. D570 is protonated. On the left, the starting structure of the simulation is shown, highlighting residues E558 and K1434 near the entry gate to the cholesterol conduit. In the center of the transmembrane domain, D570, H1387 and T1422 are shown. The measure distance shown on the right is indicated by a blue dashed line. In the middle, a zoom in on E558 and K1434 is shown, showing an overlay of the end structures of 5 independent simulations. On the right, the time evolution of the distance between the carboxylate carbon CD of the glutamate sidechain of E558 and the nitrogen NZ atom of the lysine sidechain of K1434 is shown. **B.** Same data as in A, but simulations of apo PfNCR1 in the closed SSD conformation with charged D570 and double protonated H1387. The starting structure is shown on the left, in the middle the overlay of the end structures of the 5 simulations and on the right the distance between the carboxylate carbon CD of the glutamate sidechain of E558 and the nitrogen NZ atom of the lysine sidechain of K1434.

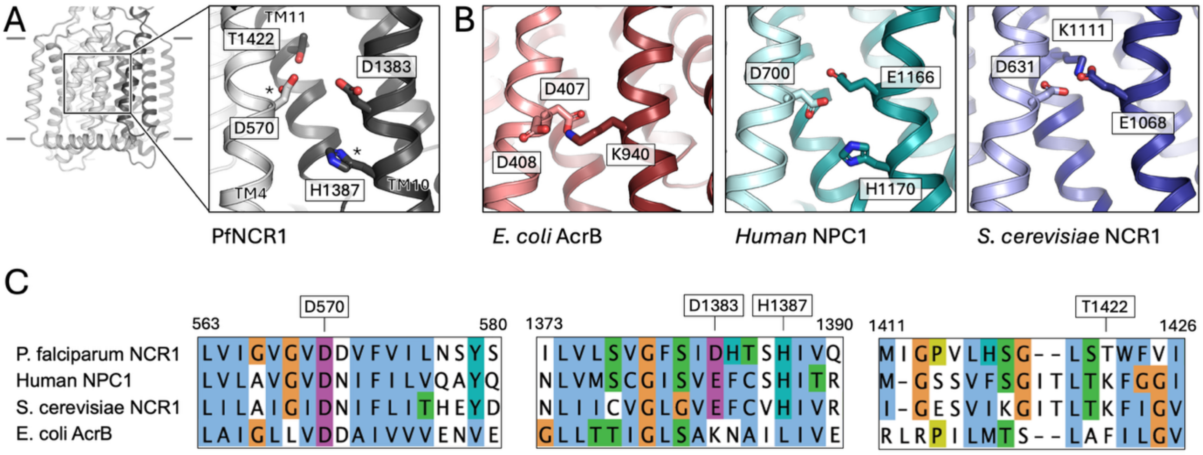


**Figure S23:** Residues involved in proton coupling in selected RND transporters. **A.** The key residues (D570 & H1387 marked with an asterisk) in the core of PfNCR1 revealed to be involved in protonation/deprotonation in distinct SSD conformations. **B.** Equivalent residues in human NPC1 (PDB ID: 6W5T [43]), *S. cerevisiae* NCR1 (PDB ID: 8QED [44]), and in *E.coli* AcrB (PDB ID: 1IWG [45]). The residues shown for each protein have been suggested to play a role in proton coupling for each of the corresponding proteins. **C.** Sequence alignment of the selected transporters, focusing on the regions of interest shown in B.

**Table S1**. Cryo-EM analysis and statistics of PfNCR1.

| **PfNCR1** | **Apo** | **MMV009108** Compound 1 | **MMV028038** Compound 2 | **MMV019662** Compound 3 | **G856-4236** Compound 4 |
|---|---|---|---|---|---|
| Instrument | FEI Titan Krios / Gatan K3 Summit | | | | |
| Magnification | 130000 | | | | |
| Voltage (kV) | 300 | | | | |
| Data-set 1 | 55 e-/Å$^2$ | 53 e-/Å$^2$ | 60 e-/Å$^2$ | 53 e-/Å$^2$ | 58 e-/Å$^2$ |
| Data-set 2 | 55 e-/Å$^2$ | - | 60 e-/Å$^2$ | 57 e-/Å$^2$ | 50 e-/Å$^2$ |
| Defocus range (µm) | -0.5 to -3.0 | | | | |
| Pixel size (Å) | 0.66 | 0.65 | 0.65 | 0.65 | 0.65 |
| Number of particles | 131581 | 125812 | 192240 | 293678 | 335464 |
| Map symmetry | C1 | | | | |
| Model resolution at FSC threshold 0.143 | 3.04 Å | 3.89 Å | 3.73 Å | 3.69 Å | 3.78 Å |
| Map CC | 0.83 | 0.78 | 0.79 | 0.73 | 0.77 |
| Protein residues/ligands | 910/1 | 910/2 | 910/3 | 910/1 | 910/2 |
| Bond length (r.m.s.d) | 0.004 | 0.004 | 0.003 | 0.004 | 0.004 |
| Bond angle (r.m.s.d) | 0.547 | 0.647 | 0.607 | 0.730 | 0.668 |
| MolProbity score | 1.70 | 2.16 | 2.07 | 2.18 | 2.31 |
| Clashscore | 12.96 | 22.56 | 18.26 | 22.96 | 24.74 |
| Rotamer outlier (%) | 0.00 | 0.00 | 0.00 | 0.00 | 0.12 |
| Favored (%) | 97.65 | 95.53 | 95.53 | 95.3 | 93.51 |
| Allowed (%) | 2.35 | 4.47 | 4.47 | 4.7 | 6.49 |
| Disallowed (%) | 0 | 0 | 0 | 0 | 0 |

**Table S2**. Compounds selected from virtual screening for experimental evaluation.

| Molport ID | Supplier ID | Supplier Name | SMILES | in silico method |
|---|---|---|---|---|
| MOLPORT-047-776-970 | 5102735 | ChemBridge Corporation | [Na+].Nc1cc(cc2cc(c(\N=N\c3ccc(cc3)S(O)(=O)=O)c(O)c12)S([O-])(=O)=O)S(O)(=O)=O | docking based on apo-PfNCR1 |
| MOLPORT-047-973-035 | AR002OQ4 | Aaron Chemicals LLC | [Na+].[Na+].[Na+].Oc1c(N=Nc2ccc(cc2)S(O)(=O)=O)c(cc2cc(cc([O-])c12)S([O-])(=O)=O)S([O-])(=O)=O | docking based on apo-PfNCR1 |
| MOLPORT-051-541-661 | AR003UJV | Aaron Chemicals LLC | [NaH].Oc1cc(cc2cc(c(\N=N\c3cccc4ccccc34)c(O)c12)S(O)(=O)=O)S(O)(=O)=O | docking based on apo-PfNCR1 |
| MOLPORT-051-435-288 | AR00IMC2 | Aaron Chemicals LLC | [Na+].[Na+].[O-]S(=O)(=O)C1=Cc2cc(cc(Nc3nc(Cl)nc(Cl)n3)c2C(=O)C1=NNc1ccccc1)S([O-])(=O)=O | docking based on apo-PfNCR1 |
| MOLPORT-047-660-117 | AR00CYV7 | Aaron Chemicals LLC | Oc1ccc(\N=N\c2cc(cc3cc(cc(O)c23)S(O)(=O)=O)S(O)(=O)=O)c(O)c1 | docking based on apo-PfNCR1 |
| MOLPORT-028-883-617 | V029-3356 | ChemDiv, Inc. | COc1ccc(C)cc1N1C(=O)C(Nc2ccccc2)=C(Sc2ccc(NC(=O)c3ccc(F)cc3)cc2)C1=O | docking based on PfNCR1-MMV028038 |
| MOLPORT-039-343-000 | STL441565 | Vitas M Chemical Limited | CN1C(=O)\C(=C\c2cn(CCOc3ccc(cc3)C(C)(C)C)c3ccccc23)c2cc(ccc12)S(=O)(=O)N1CCOCC1 | docking based on PfNCR1-MMV019662 (neck) |
| MOLPORT-000-779-126 | STL055711 | Vitas M Chemical Limited | CC(C)c1ccc(cc1)C1N(Cc2cc[nH+]cc2)C(=O)C(=O)\C1=C(\[O-])c1ccc(OCc2cccc(C)c2)cc1 | docking based on PfNCR1-MMV019662 |

| | | | | |
|---|---|---|---|---|
| | | | | (neck) |
| MOLPORT-003-137-985 | F2573-0329 | Life Chemicals Inc. | COc1ccc(cc1)N1CCN(CC1)C(CNC(=O)Oc1ccccc1)c1ccc2OCOc2c1 | similarity search |
| MOLPORT-003-138-036 | G856-4269 | ChemDiv, Inc. | Fc1ccc(cc1)N1CCN(CC1)C(CNC(=O)c1ccc(Cl)cc1)c1ccc2OCOc2c1 | similarity search |
| MOLPORT-003-373-489 | G856-4236 | ChemDiv, Inc. | COc1ccc(cc1)N1CCN(CC1)C(CNC(=O)c1c(F)cccc1F)c1ccc2OCOc2c1 | similarity search |
| MOLPORT-003-137-979 | F2573-0310 | Life Chemicals Inc. | Cc1ccc(cc1)C(=O)NCC(N1CCN(CC1)c1ccccc1)c1ccc2OCOc2c1 | similarity search |

**Table S3: pKa predictions of protonatable residues carried out using PROPKA**

| Residue | Predicted pKa for PfNCR1 in the closed conformation | Predicted pKa for PfNCR1 in the open conformation |
|---|---|---|
| GLU12 | 3.84 | 4.15 |
| ASP19 | 3.92 | 3.83 |
| ASP27 | 4.04 | 3.94 |
| ASP34 | 3.48 | 3.83 |
| GLU58 | 4.46 | 4.38 |
| HIS59 | 5.09 | 5.6 |
| GLU60 | 5.1 | 5.18 |
| ASP62 | 4.03 | 4 |
| GLU76 | 4.92 | 3.95 |
| GLU79 | 5.02 | 4.35 |
| ASP83 | 5.76 | 5.65 |
| GLU96 | 3.28 | 3.64 |
| GLU109 | 4.72 | 4.83 |
| GLU114 | 6.42 | 7.23 |
| GLU115 | 4.59 | 3.61 |
| ASP119 | 4 | 4.02 |
| GLU121 | 3.86 | 5 |
| ASP123 | 4.16 | 4.03 |
| GLU126 | 4.89 | 3.86 |
| GLU129 | 5.44 | 4.63 |
| HIS142 | 7.17 | 6.58 |
| GLU146 | 4.26 | 4.2 |

| | | |
|---|---|---|
| GLU156 | 4.81 | 4.56 |
| GLU157 | 4.59 | 3.97 |
| ASP170 | 3.88 | 3.95 |
| ASP173 | 3.07 | 3.21 |
| ASP175 | 3.93 | 3.78 |
| GLU176 | 4.39 | 4.64 |
| ASP297 | 3.89 | 3.97 |
| ASP300 | 3.69 | 3.26 |
| ASP301 | 5.28 | 4.79 |
| ASP317 | 3.6 | 3.85 |
| ASP333 | 3.43 | 2.67 |
| GLU336 | 4.76 | 4.8 |
| ASP339 | 3.78 | 3.7 |
| GLU350 | 4.08 | 4.94 |
| ASP351 | 4.37 | 4.44 |
| GLU354 | 5.49 | 5.04 |
| ASP359 | 3.47 | 3.47 |
| GLU370 | 4.47 | 4.15 |
| ASP377 | 3.32 | 3.02 |
| ASP383 | 2.93 | 3.65 |
| ASP386 | 3.87 | 4.09 |
| GLU396 | 4.52 | 6.1 |
| GLU409 | 4.26 | 4.62 |
| HIS433 | 4.96 | 5.27 |
| GLU436 | 4.53 | 5.52 |

| | | |
|---|---|---|
| GLU443 | 6.62 | 7.74 |
| ASP448 | 4.27 | 4.07 |
| ASP456 | 4.34 | 3.42 |
| ASP457 | 3.99 | 4.03 |
| ASP461 | 3.84 | 3.73 |
| GLU462 | 4.47 | 4.74 |
| GLU463 | 4.53 | 4.76 |
| ASP466 | 3.92 | 3.49 |
| HIS475 | 6.18 | 5.51 |
| ASP479 | 4.95 | 5.38 |
| GLU483 | 5.57 | 5.84 |
| ASP484 | 3.89 | 4.29 |
| GLU485 | 6.33 | 4.93 |
| ASP487 | 4.04 | 4.61 |
| ASP493 | 2.69 | 2.28 |
| GLU558 | 7.43 | 5.7 |
| ASP570 | 7.88 | 3.98 |
| ASP571 | 8.93 | 8.5 |
| ASP587 | 3.75 | 2.87 |
| ASP598 | 5.22 | 4.65 |
| GLU654 | 5.95 | 5.87 |
| GLU658 | 3.81 | 4.89 |
| GLU1051 | 3.96 | 3.87 |
| ASP1106 | 3.9 | 4.32 |
| ASP1112 | 3.72 | 3.01 |

| | | |
|---|---|---|
| ASP1128 | 3.88 | 4.29 |
| ASP1131 | 4.71 | 5.9 |
| GLU1134 | 4.37 | 5 |
| ASP1139 | 5.12 | 4.85 |
| HIS1141 | 6.04 | 6.03 |
| GLU1150 | 4.95 | 4.63 |
| HIS1181 | 7.8 | 6.56 |
| GLU1183 | 4.74 | 5.97 |
| GLU1189 | 4.52 | 4.66 |
| GLU1190 | 4.54 | 5.17 |
| HIS1195 | 5.12 | 5.05 |
| GLU1199 | 5.55 | 4.63 |
| GLU1202 | 4.65 | 4.57 |
| GLU1225 | 4.51 | 4.7 |
| GLU1229 | 4.59 | 4.55 |
| GLU1232 | 4.64 | 4.94 |
| ASP1241 | 3.77 | 3.97 |
| ASP1250 | 5.81 | 5.19 |
| HIS1264 | 2.62 | 2.54 |
| ASP1271 | 3.63 | 3.28 |
| ASP1272 | 3.99 | 4.1 |
| GLU1274 | 4.65 | 5.05 |
| GLU1290 | 4.38 | 4.36 |
| HIS1292 | 6.44 | 5.99 |
| HIS1300 | 5.54 | 4.19 |

| | | |
|---|---|---|
| GLU1307 | 5.82 | 5.32 |
| ASP1309 | 6.02 | 6.44 |
| GLU1310 | 4.58 | 4.08 |
| GLU1314 | 4.63 | 4.71 |
| ASP1352 | 6.95 | 7.28 |
| ASP1383 | 9.19 | 9.37 |
| HIS1384 | 3.42 | 4.3 |
| HIS1387 | 4.13 | 7.25 |
| HIS1394 | 5.2 | 5.09 |
| ASP1401 | 3.39 | 3.05 |
| GLU1402 | 3.69 | 3.5 |
| GLU1406 | 4.02 | 4.67 |
| HIS1409 | 5 | 5.75 |
| HIS1417 | 4.1 | 4.04 |
| ASP1435 | 4.59 | 5.16 |

#### References


1. Kubala, M.H., et al., *Structural and thermodynamic analysis of the GFP:GFP-nanobody complex.* Protein Sci, 2010. **19**(12): p. 2389-401.
2. Scheres, S.H., *RELION: implementation of a Bayesian approach to cryo-EM structure determination.* J Struct Biol, 2012. **180**(3): p. 519-30.
3. Punjani, A., et al., *cryoSPARC: algorithms for rapid unsupervised cryo-EM structure determination.* Nat Methods, 2017. **14**(3): p. 290-296.
4. Zheng, S.Q., et al., *MotionCor2: anisotropic correction of beam-induced motion for improved cryo-electron microscopy.* Nat Methods, 2017. **14**(4): p. 331-332.
5. Rohou, A. and N. Grigorieff, *CTFFIND4: Fast and accurate defocus estimation from electron micrographs.* J Struct Biol, 2015. **192**(2): p. 216-21.
6. Afonine, P.V., et al., *Real-space refinement in PHENIX for cryo-EM and crystallography.* Acta Crystallogr D Struct Biol, 2018. **74**(Pt 6): p. 531-544.
7. Tiburcio, M., et al., *A Novel Tool for the Generation of Conditional Knockouts To Study Gene Function across the Plasmodium falciparum Life Cycle.* mBio, 2019. **10**(5).
8. Istvan, E.S., et al., *Plasmodium Niemann-Pick type C1-related protein is a druggable target required for parasite membrane homeostasis.* Elife, 2019. **8**.
9. Gradisch, R., et al., *Ligand coupling mechanism of the human serotonin transporter differentiates substrates from inhibitors.* Nat Commun, 2024. **15**(1): p. 417.
10. Khanppnavar, B., et al., *Structural basis of organic cation transporter-3 inhibition.* Nat Commun, 2022. **13**(1): p. 6714.
11. Lomize, M.A., et al., *OPM database and PPM web server: resources for positioning of proteins in membranes.* Nucleic Acids Res, 2012. **40**(Database issue): p. D370-6.
12. Kroon, P.C., et al., *Martinize2 and Vermouth: Unified Framework for Topology Generation*. 2025, eLife Sciences Publications, Ltd.
13. Souza, P.C.T., et al., *Martini 3: a general purpose force field for coarse-grained molecular dynamics.* Nat Methods, 2021. **18**(4): p. 382-388.
14. Wassenaar, T.A., et al., *Computational Lipidomics with insane: A Versatile Tool for Generating Custom Membranes for Molecular Simulations.* J Chem Theory Comput, 2015. **11**(5): p. 2144-55.
15. Abraham, M.J., et al., *GROMACS: High performance molecular simulations through multi-level parallelism from laptops to supercomputers.* SoftwareX, 2015. **1-2**: p. 19-25.
16. Hess, B., et al., *GROMACS 4: Algorithms for Highly Efficient, Load-Balanced, and Scalable Molecular Simulation.* J Chem Theory Comput, 2008. **4**(3): p. 435-47.
17. Van Der Spoel, D., et al., *GROMACS: fast, flexible, and free.* J Comput Chem, 2005. **26**(16): p. 1701-18.
18. Bussi, G., D. Donadio, and M. Parrinello, *Canonical sampling through velocity rescaling.* Journal of Chemical Physics, 2007. **126**(1).
19. Bernetti, M. and G. Bussi, *Pressure control using stochastic cell rescaling.* J Chem Phys, 2020. **153**(11): p. 114107.
20. Wassenaar, T.A., et al., *Going Backward: A Flexible Geometric Approach to Reverse Transformation from Coarse Grained to Atomistic Models.* J Chem Theory Comput, 2014. **10**(2): p. 676-90.

21. Wolf, M.G., et al., *g_membed: Efficient Insertion of a Membrane Protein into an Equilibrated Lipid Bilayer with Minimal Perturbation.* Journal of Computational Chemistry, 2010. **31**(11): p. 2169-2174.
22. Szollosi, D. and T. Stockner, *Sodium Binding Stabilizes the Outward-Open State of SERT by Limiting Bundle Domain Motions.* Cells, 2022. **11**(2).
23. Olsson, M.H., et al., *PROPKA3: Consistent Treatment of Internal and Surface Residues in Empirical pKa Predictions.* J Chem Theory Comput, 2011. **7**(2): p. 525-37.
24. Lindorff-Larsen, K., et al., *Improved side-chain torsion potentials for the Amber ff99SB protein force field.* Proteins-Structure Function and Bioinformatics, 2010. **78**(8): p. 1950-1958.
25. Jämbeck, J.P.M. and A.P. Lyubartsev, *An Extension and Further Validation of an All-Atomistic Force Field for Biological Membranes.* Journal of Chemical Theory and Computation, 2012. **8**(8): p. 2938-2948.
26. Jämbeck, J.P.M. and A.P. Lyubartsev, *Another Piece of the Membrane Puzzle: Extending Slipids Further.* Journal of Chemical Theory and Computation, 2013. **9**(1): p. 774-784.
27. Kagami, L., et al., *The ACPYPE web server for small-molecule MD topology generation.* Bioinformatics, 2023. **39**(6).
28. Sousa da Silva, A.W. and W.F. Vranken, *ACPYPE - AnteChamber PYthon Parser interfacE.* BMC Res Notes, 2012. **5**: p. 367.
29. He, X., et al., *A fast and high-quality charge model for the next generation general AMBER force field.* J Chem Phys, 2020. **153**(11): p. 114502.
30. Gradisch, R., et al., *Occlusion of the human serotonin transporter is mediated by serotonin-induced conformational changes in the bundle domain.* Journal of Biological Chemistry, 2022. **298**(3).
31. Parrinello, M. and A. Rahman, *Polymorphic Transitions in Single-Crystals - a New Molecular-Dynamics Method.* Journal of Applied Physics, 1981. **52**(12): p. 7182-7190.
32. Darden, T., D. York, and L. Pedersen, *Particle Mesh Ewald - an N.Log(N) Method for Ewald Sums in Large Systems.* Journal of Chemical Physics, 1993. **98**(12): p. 10089-10092.
33. Gowers, R.J., et al. *MDAnalysis: A Python Package for the Rapid Analysis of Molecular Dynamics Simulations*. 2019. United States.
34. Michaud-Agrawal, N., et al., *Software News and Updates MDAnalysis: A Toolkit for the Analysis of Molecular Dynamics Simulations.* Journal of Computational Chemistry, 2011. **32**(10): p. 2319-2327.
35. Humphrey, W., A. Dalke, and K. Schulten, *VMD: visual molecular dynamics.* J Mol Graph, 1996. **14**(1): p. 33-8, 27-8.
36. *Schrödinger Release 2022-4: LigPrep, Schrödinger, LLC, New Tork, NY, 2022,* [www.schrodinger.com](http://www.schrodinger.com).
37. Lu, C., et al., *OPLS4: Improving Force Field Accuracy on Challenging Regimes of Chemical Space.* J Chem Theory Comput, 2021. **17**(7): p. 4291-4300.
38. Shelley, J.C., et al., *Epik: a software program for pK( a ) prediction and protonation state generation for drug-like molecules.* J Comput Aided Mol Des, 2007. **21**(12): p. 681-91.
39. *Schrödinger Release 2022-4: Protein Preparation Wizard; Epik, Schrödinger, LLC, New York, NY, 2022.*

40. Sastry, G.M., et al., *Protein and ligand preparation: parameters, protocols, and influence on virtual screening enrichments.* J Comput Aided Mol Des, 2013. **27**(3): p. 221-34.
41. Jones, G., et al., *Development and validation of a genetic algorithm for flexible docking.* J Mol Biol, 1997. **267**(3): p. 727-48.
42. Zhang, Z., et al., *The Plasmodium falciparum NCR1 transporter is an antimalarial target that exports cholesterol from the parasite's plasma membrane.* Sci Adv, 2024. **10**(51): p. eadq6651.
43. Qian, H., et al., *Structural Basis of Low-pH-Dependent Lysosomal Cholesterol Egress by NPC1 and NPC2.* Cell, 2020. **182**(1): p. 98-111 e18.
44. Frain, K.M., et al., *Conformational changes in the Niemann-Pick type C1 protein NCR1 drive sterol translocation.* Proc Natl Acad Sci U S A, 2024. **121**(15): p. e2315575121.
45. Murakami, S., et al., *Crystal structure of bacterial multidrug efflux transporter AcrB.* Nature, 2002. **419**(6907): p. 587-93.
